# Novel protein networks in motile cilia from human epithelial cells revealed by cryo-ET and proteomics analysis of cilia from PCD patients

**DOI:** 10.64898/2026.09.28.755144

**Authors:** Charlotte de Ceuninck, Tamino Cairoli, Pavel Afanasyev, Yanxun Li, Eva Markovic, Alex Jud, Alexander Leitner, Loretta Müller, Takashi Ishikawa

## Abstract

In our past study on human cilia from primary ciliary dyskinesia (PCD) patients using cryo-electron tomography and mass spectrometry, defects in the *DNAH5* gene were proven to cause the loss of the entire outer dynein arm along with other proteins. This phenomenon was not observed in the unicellular green alga *Chlamydomonas*. In this study, we examined the loss of ciliary components caused by defects in various genes, based on proteomic and structural analyses of motile cilia from multiple PCD patients. The absence of single outer arm dynein genes causes a loss or significant decrease in other inner and outer arm dyneins. Interestingly, some patients with outer dynein defects show decreased intraflagellar transport (IFT) proteins, suggesting an influence of cargo components or assembly on transport. These phenomena were not observed in *Chlamydomonas*, suggesting a more complex mechanism of ciliogenesis in humans despite the high similarity in the final 3D architecture of the 9+2 axoneme. Defects in certain central pair proteins unexpectedly reduced components on the doublets as well. Our results demonstrate protein-protein interactions during stages of human motile ciliogenesis that are more complex than those in unicellular organisms.

## Introduction

Motile cilia are beating organelles with a diameter of ∼300 nm and a length of 5–20μm. They function in human tissues such as trachea, brain ventricles, oviducts, sperm, Eustachian tubes, and embryonic nodes. Cilia across most species, from unicellular algae to humans, share a nearly identical structure called the “9+2” arrangement. In this architecture, nine doublet microtubules form a spoke-like arrangement and are connected by dynein ATPase motor proteins assembled into two complexes: outer and inner arm dyneins (ODA and IDA) (summarized in (Ishikawa, 2016)). Defects in motile cilia cause a wide variety of symptoms collectively termed primary ciliary dyskinesia (PCD) (Wallmeier et al, 2020).

The structure of motile cilia is highly conserved from the green alga *Chlamydomonas* to humans (Fig. 1). In the axoneme—the region protruding from the cell that is responsible for force generation—nine doublet microtubules with highly repetitive 96 nm units surround two singlet microtubules decorated by numerous proteins (the central pair apparatus, CPA) (Gui et al, 2022). From each 96 nm unit of the doublet, two or three T-shaped protein complexes called radial spokes (RS) extend toward the CPA. The T-shaped RS consists of head and stalk domains (Pigino et al, 2011; Gui et al, 2021). Between adjacent doublets, dynein motor proteins form two types of complexes—inner and outer dynein arms (IDA and ODA)—that enable sliding motion between doublets. In *Chlamydomonas*, there are multiple isoforms of dyneins in the IDA and ODA, designated as dyneins a, b, c, d, and e (all single-headed) and f (heterodimeric, with fα and β chains) for the IDA, and dyneins, and for the ODA (Fig. 1).

**Fig. 1:**
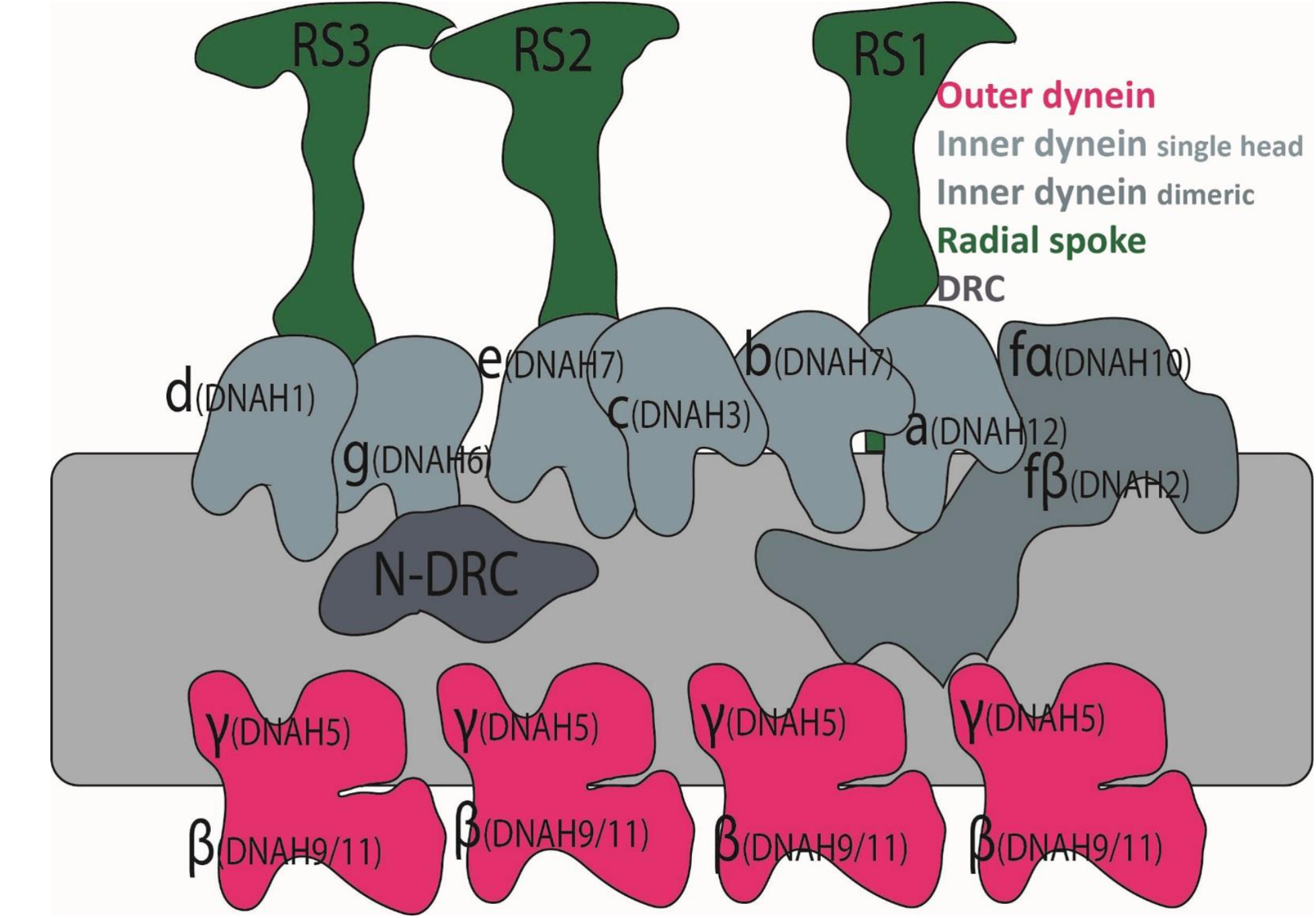
Schematic diagram of ciliary components on doublet microtubules in human motile cilia (3D layout). Major structures (outer/inner dynein arms, radial spokes, N-DRC) are color-coded. Dynein nomenclature is shown for both *Chlamydomonas* (uppercase) and human (lowercase in parentheses).

The human axoneme has a similar arrangement of dyneins, except that the ODA contains two dynein heavy chains. These are designated DNAH1–11, among which DNAH9 and DNAH11 occupy the position corresponding to *Chlamydomonas* outer dynein in the distal and proximal regions, respectively (Dougherty et al, 2016). In total, more than 400 proteins have been identified in the axoneme by mass spectrometry (MS)-based proteomics (Pazour et al, 2005). Out of these axonemal proteins, the locations and conformations of more than 100 have been revealed by single-particle cryo-EM analysis (Walton et al, 2023; Gui et al, 2022). How these proteins interact with each other to coordinate beating motion remains unclear.

Until now, mechanistic studies on cilia have been carried out mainly using model organisms. For example, the roles of outer and inner dynein arms were defined as the accelerator and the regulator, respectively, based on the phenotypes of corresponding *Chlamydomonas* mutants (Kamiya, 2002). *Chlamydomonas* mutants lacking outer arms can still swim with a proper waveform, albeit more slowly and weakly (Kamiya & Okamoto, 1985). This led to assigning the roles of force amplification/acceleration and waveform determination to the outer and inner arm dyneins, respectively. However, in human cilia, although their structural arrangement is very similar to that of *Chlamydomonas*, the loss of outer arms causes ciliary paralysis (Hornef et al, 2006; Fliegauf et al, 2007). The reason why the loss of ODA results in different phenotypes across species has not yet been elucidated.

The mechanisms underlying PCD are tightly linked not only to ciliary beating function, but also to the assembly process required to build the complex architecture of the axoneme. To date, ∼60 causal genes for PCD have been identified (Suppl. Table 1A). Approximately 50 of these encode structural ciliary components, while the remainder are considered cytoplasmic assembly factors (Fliegauf et al, 2007; Despotes et al, 2024). Diagnosis of PCD is carried out using this gene panel, alongside high-speed video microscopy (HSVM), classical transmission electron microscopy (TEM) of resin-embedded and stained thin sections, and nasal nitric oxide (nNO) measurement (Goutaki et al, 2019; Fernandez-Gonzalez et al, 2009; Kott et al, 2013).

Modern techniques, such as 3D structural analysis by cryo-TEM and MS-based proteomics, offer diagnostic insights. In our previous study (De Ceuninck Van Capelle et al, 2025), we analyzed the 3D structure and proteome of bronchial and nasal epithelial cilia from healthy individuals and PCD patients. We found that in three patients with distinct truncation mutations in *DNAH5* (near the N-terminus, in the middle, and near the C-terminus), the outer dynein arm was completely lost. This stands in stark contrast to unicellular green algae (*Chlamydomonas*), where the loss of a single outer arm dynein gene results in the loss of only the corresponding density without broader disruption, as observed by cryo-electron tomography (cryo-ET) (Ishikawa et al, 2007). Furthermore, cilia from PCD patients with *DNAH5* defects are immotile (Hornef et al, 2006), unlike *Chlamydomonas* outer arm mutants, which retain flagellar beating and can swim slowly (Kamiya, 2002).

In this study, we systematically examined protein components in nasal motile cilia from PCD patients. Proteomics by MS and structural analysis using cryo-ET with subtomogram averaging (STA) revealed distinct protein networks in human cilia. MS-based proteomics identified changes in ciliary components with statistical significance, while cryo-ET and STA demonstrated the presence or absence of specific proteins within the 96 nm periodic repeat along doublet microtubules.

## Results

In this study, we analyzed the structure and proteome of human epithelial cilia using cryo-ET/STA and MS to examine interdependencies among ciliary component proteins. First, we confirmed the structure of wild-type (WT) axonemes from healthy cilia and profiled their proteome. We then performed the same analysis on cilia from several patients exhibiting different ciliary defects. This allowed us to correlate protein losses confirmed by MS with structural changes observed via cryo-ET, as well as downstream changes in the abundance of proteins critical for ciliary function and architecture.

### WT Axoneme Structure

In the 3D reconstruction of WT (healthy) human cilia (Fig. 2A, B), density is significantly reduced at the locations of two inner dyneins: b (corresponding to the *Chlamydomonas* ortholog DHC5 (Bui et al, 2012)) and c (corresponding to *Chlamydomonas* DHC9) (Figs. 1&2B), consistent with previous reports (Lin et al, 2014) (Fig. 2C). According to high-resolution single-particle cryo-EM analysis (Walton et al, 2023), these loci in human cilia are occupied by DNAH7 and DNAH3, respectively (Fig. 2D), both of which were detected in our MS proteomics data (Suppl. Table 1B).

**Fig. 2:**
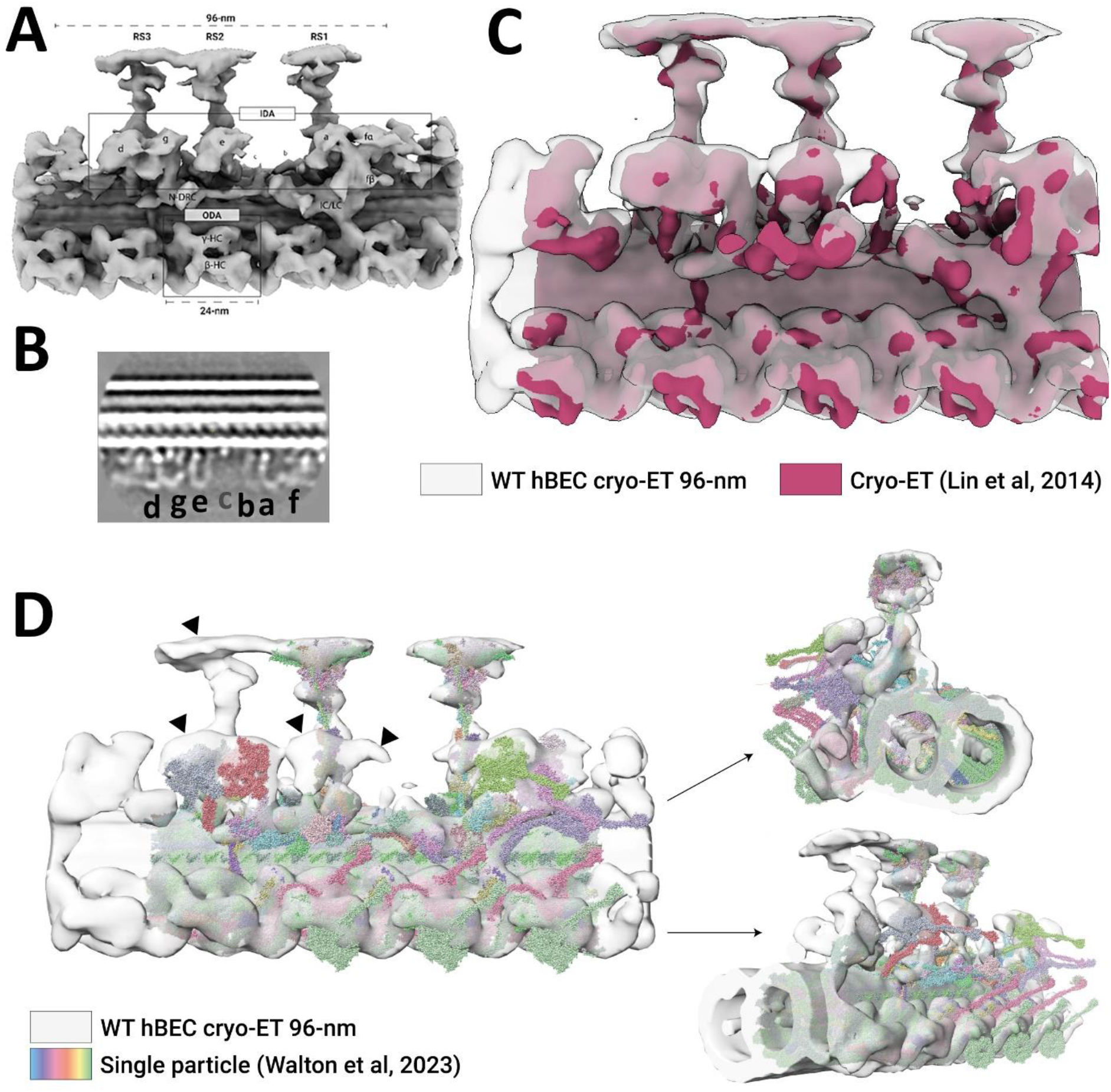
(A) 3D structure of doublet microtubules from WT human ciliated bronchial epithelial cells. (B) Section from the map including inner dyneins. Inner dyneins a – f are indicated. Since dynein c density is weak, indication of dynein c is also in grey. (C, D) Its comparison with previously reported structures; (C) Lin et al. 2014 and (D) Walton et al. 2023.

The overall structure of WT cilia is similar to the previous report of cryo-ET studies (Lin et al, 2014), with minor differences at the outer dynein arm, where outer dyneins lie closer to the inner dyneins in our structure - likely reflect dynein ATP hydrolysis and power-stroke states. Both cryo-ET structures highlight additional density (indicated by arrowheads in Fig. 2C) not present in the high-resolution single-particle cryo-EM structure (Walton et al, 2023). This likely represents flexible regions on microtubule doublets isolated from the axoneme that were averaged out during single-particle processing, suggesting the presence of unidentified proteins.

Using mass spectrometry, >1,000 proteins were identified from isolated cilia of bronchial epithelial cells (Suppl. Table 1B). This dataset served as our control baseline for analyzing cilia from PCD patients.

### Patient Summary

To investigate how genetic defects in individual ciliary components impact the assembly of motile cilia, we performed MS-based proteomics and cryo-ET/STA on nasal epithelial cilia from 15 patients. Genetic defects, motility phenotypes, diagnostic properties, and clinical symptoms are summarized in Table 1. Genetic investigations were performed using the standard PCD gene panel (Suppl. Table 1A).

**Table 1.** PCD patients, who provided cilia for this study. See the additional Excel file for the whole view.

|  | Gene of defect | Ciliary beating pattern | Other diagnostic test results |
| --- | --- | --- | --- |
| **PD122** | DNAH5[4348C→T]<br>? [13692093 13944547del] | immotile | TEM: hallmark class 1 ODA defect<br>IF: DNAH5, DNAI1, DNAI2, DNAH11, DNAH9 missing; GAS8 present, DNALI1 rather present |
| **PD441** | DNAH5[8998C→T] [799G→T] | immotile | TEM: hallmark class 1 ODA defect<br>IF: DNAH5, DNAI1, DNAI2, DNAH9 and DNAH11 missing; GAS8 present |
| **PD480** | DNAH5[13194 13197del]<br>[11761+1 11762-1) (11883+1 11884-1)del] | immotile | TEM: hallmark class 1 ODA defect<br>IF; DNAH5, DNAH9,<br>DNAH11, DNAI1 and DNAI2 missing, GAS8 and DNALI1 present. |
| **PD435** | DNAH11[4333C→T] [10523A→G] | vibrating | IF: DNAH11 missing, DNAH5, DNAH9, GAS8, DNAI1, DNAI2, DNALI1 present |
| **PD442** | N.A. | vibrating | IF: DNAH5, GAS8 and RSPH9 present, DNAH11 missing |
| **PD551** | DNAH11[7099 delA][11327T→C] | vibrating | TEM: no evidence for PCD<br>IF: DNAH11 missing, DNAH5 partially missing, DNAI1, DNAI2, DNALI1, GAS8, RSPH9 present, DNAH9 inconclusive |
| PD222 | DNAH9 cis[12177G→A][12640G→T] | normal | TEM: no evidence for PCD<br>IF: DNAH5 present, DNAH9 present in full axoneme length |
| PD272 | DNAH9 cis[12177G→A][12640G→T] | normal | TEM: class 2 ODA defect<br>IF: DNAH5 present, DNAH9 present in full axoneme length |
| PD691 | Biallelic likely pathogenic mutations in DNAH1 and DNAH11 (still under investigation) [**] | Immotile, some residual vibration | DNAH1 and DNAH11 missing, DNAH5 DNAI1, DNAI2, DNALI1, GAS8, RSPH9 present |
| PD311 | hydin [7585del] | normal | TEM: normal<br>IF: SPEF2, DNAH5, GAS8, RSPH9 present |
| PD343 | hydin[2647 2648][5956dupA] | Reduced bending and amplitude | IF: DNAH5, RSPH9, DNAH11, GAS8 present, SPEF2 missing |
| PD358 | hydin [7585del] | Normal | TEM: normal IF: SPEF2 present |
| PD403 | Not detected within the investigated PCD panel (114 genes) | Partially rotating (in cell culture 50% rotation, 50% normal beating) | TEM: not enough evidence for PCD (20% IDA, 10% Central Complex defects)<br>IF: DNAH5, GAS8, DNAH11, RSPH1, RSPH4a, RSPH9 present |
| PD428 | N.A. | Immotile, residual vibration | IF: GAS8, RSPH9, DNAI1, DNAI2, DNAH11 present, DNAH5 partially present |
| PD461 | Not detected within the investigated PCD gene panel (47 genes), biallelic variants of unknown significance in HYDIN (c.5553T>A p.(Asn1851Lys) (trio exome analysis) | Immotile, some cells with long cilia that have pathologic slow residual movement | TEM: not enough evidence for PCD, but 72% orientation distraction<br>IF: DNAH5, GAS8, RSPH9, DNAH11 present |

### Defect in a Single Outer Dynein Gene Causes Loss/Decrease of Other Dyneins and IFT Components

As observed in our previous work (De Ceuninck Van Capelle et al, 2025), defects in *DNAH5* lead to a total loss of the ODA. Here, we further analyzed PD441 (a *DNAH5* patient from our previous study) and PD551 (a *DNAH11* mutant). In both patients, outer dynein mutations also caused a significant reduction in inner arm dyneins, as detected by MS (Fig. 3A, B; Table 2, Suppl. Table 2). This reduction in inner arm dyneins is consistent with the blurred density observed in cryo-ET maps (Fig. 6D in (De Ceuninck Van Capelle et al, 2025)).

**Fig. 3:**
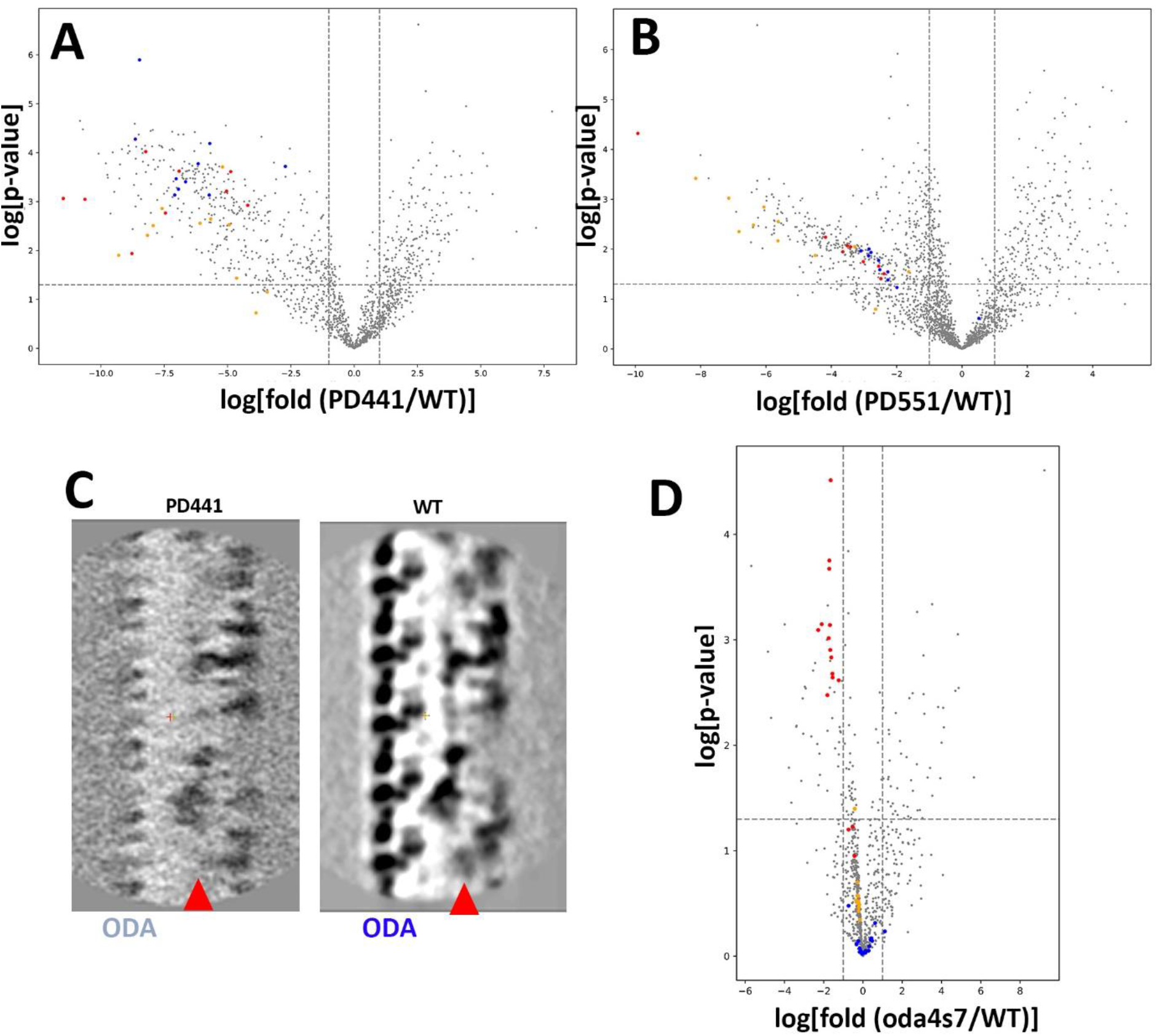
(A,B) Volcano plots from MS proteomics between data from (A) PD441 and WT and (B) between MS of PD551 and WT. (C) Structure of PD441 cilia by cryo-ET. (D) MS of cilia from the *Chlamydomonas* oda4-s7 mutant. Outer arm dyneins are shown in red dots. Inner arm dyneins in brown. IFT proteins in blue.

**Table 2.** MS data of 5 patients to show proteins which decreased significantly in the patients’ cilia compared to WT (marked by -). For full datasets of all the patients, refer Suppl. Table 2.

| Protein names | Gene names | DNAH5 PD122 | DNAH5 PD441 | DNAH5 PD480 | DNAH11 PD442 | DNAH11 PD551 |
| --- | --- | --- | --- | --- | --- | --- |
| Dynein axonemal heavy chain 1 | DNAH1 | - | - | - | - | - |
| Dynein axonemal heavy chain 10 | DNAH10 | - | - | - | - | - |
| Dynein axonemal heavy chain 11 | DNAH11 | - |  |  | - | - |
| Dynein axonemal heavy chain 12 | DNAH12 | - | - | - | - | - |
| Dynein axonemal heavy chain 2 | DNAH2 | - | - | - | - | - |
| Dynein axonemal heavy chain 3 | DNAH3 |  | - |  |  |  |
| Dynein axonemal heavy chain 5 | DNAH5 | - |  |  | - | - |
| Dynein axonemal heavy chain 6 | DNAH6 | - | - | - | - | - |
| Dynein axonemal heavy chain 7 | DNAH7 | - | - | - | - | - |
| Dynein axonemal heavy chain 9 | DNAH9 | - | - | - | - | - |
| Dynein axonemal intermediate chain 1 | DNAI1 | - | - | - | - | - |
| Dynein axonemal intermediate chain 2 | DNAI2 | - | - | - | - | - |
| Dynein axonemal intermediate chain 3 | DNAI3 | - | - | - | - | - |
| Dynein axonemal intermediate chain 4 | DNAI4 | - | - | - | - | - |
| Dynein axonemal intermediate chain 7 | DNAI7 | - | - | - | - | - |

We systematically evaluated the effect of *DNAH5* mutations using MS. Volcano plots displaying fold-changes and p-values for ciliary components from patient PD441 compared to healthy controls demonstrated that more than 40 proteins were significantly decreased in the *DNAH5* mutant (Fig. 3A). These affected components included not only outer dyneins (shown in red, Fig. 3A), but also dyneins more broadly, including inner arm and cytoplasmic dyneins (shown in brown, Fig. 3A).

Another outer dynein mutant, PD551 (lacking functional *DNAH11*), displayed a similar trend (Fig. 3B): outer dynein deletion was accompanied by a reduction in IDA proteins (red and brown data points in Fig. 3B, respectively). Cryo-ET and subtomogram averaging of PD441 cilia confirmed reduced density at IDA (indicated by red arrowheads in Fig. 3C; see also Fig. 6 in (De Ceuninck Van Capelle et al, 2025)).

For comparison, we conducted MS studies on the corresponding *Chlamydomonas* mutant, *oda4-s7*, in which the outer arm dynein heavy chain is truncated (Ishikawa et al, 2007; Sakakibara et al, 1993). Proteomic analysis of this mutant demonstrated a specific loss of outer dynein components (red points in Fig. 3D), while other dyneins remained unaffected (brown points in Fig. 3D). This indicates that although humans and *Chlamydomonas* share the basic 9+2 axonemal architecture, the mechanisms governing the assembly, transport, and localization of ciliary components differ between the species, with human ciliogenesis exhibiting higher complexity than that of unicellular organisms.

Interestingly, both human outer dynein mutants also showed decreased levels of intraflagellar transport (IFT) proteins (blue points in Fig. 3A, B). The *Chlamydomonas oda4-s7* mutant did not display this decrease in IFT proteins (blue points in Fig. 3D). This suggests a mechanism by which outer dynein loss impacts other dynein arms in humans: the absence of DNAH5 may impair IFT function, thereby disrupting the transport and assembly of other dyneins.

### Comparison of Ciliary Components Across PCD Patients

To expand on these findings, we investigated additional PCD patient cilia using MS proteomics. Results are summarized in Tables 2, 3, and 4 for ODA/IDA, RS, and IFT components, respectively.

**Table 3.** MS of radial spoke (RS) proteins from 5 patients. In each patient, RS proteins detected significantly lower than in WT are marked with ‘-’. For full datasets of all the patients, refer Suppl. Table 3.

| Protein names | Gene names | DNAH5 PD122 | DNAH5 PD441 | DNAH5 PD480 | DNAH11 PD442 | DNAH11 PD551 |
| --- | --- | --- | --- | --- | --- | --- |
| Radial spoke head 1 homolog | RSPH1 | - | - | - | - | - |
| Radial spoke head 10 homolog B | RSPH10B |  |  |  |  |  |
| Radial spoke head 10 homolog B2 | RSPH10B2 |  | - |  | - |  |
| Radial spoke head 14 homolog | RSPH14 | - | - | - | - | - |
| Radial spoke head protein 3 homolog | RSPH3 | - | - | - | - | - |
| Radial spoke head protein 4 homolog A | RSPH4A | - | - |  | - | - |

Table 2 shows that genetic defects in outer dyneins (*DNAH5* and *DNAH11*), as observed in patients PD122, PD480, and PD442, lead to a severe loss of other dyneins. Exceptionally, patients PD222 and PD272 showed only a limited decrease in dyneins, likely because they carry point mutations restricted to *cis* alleles.

Intriguingly, DNAH3 was the only dynein not markedly reduced by outer dynein mutations (Table 2, Suppl. Table 2). DNAH3 corresponds to *Chlamydomonas* dynein c (Walton et al, 2023) and appears weak in 3D structural reconstructions of WT human cilia (Fig. 2), though it is detected in MS (Suppl. Table 1B). Dynein assembly factors 1, 3, and 5 were also unaffected by these mutations, confirming that they are not responsible for the reduction in axonemal dyneins.

Patient PD691, carrying defects in *DNAH1* (corresponding to dynein d) and *DNAH11*, showed the expected loss of DNAH1 by MS, as well as a loss of density at the dynein d locus in cryo-ET maps (Fig. 4C). However, there was a discrepancy between MS and cryo-ET structural data: while PD691 exhibited significantly reduced inner and outer dynein levels by MS (Table 2; Suppl. Table 2), these structures remained visible in cryo-ET reconstructions (Fig. 4B).

**Fig. 4:**
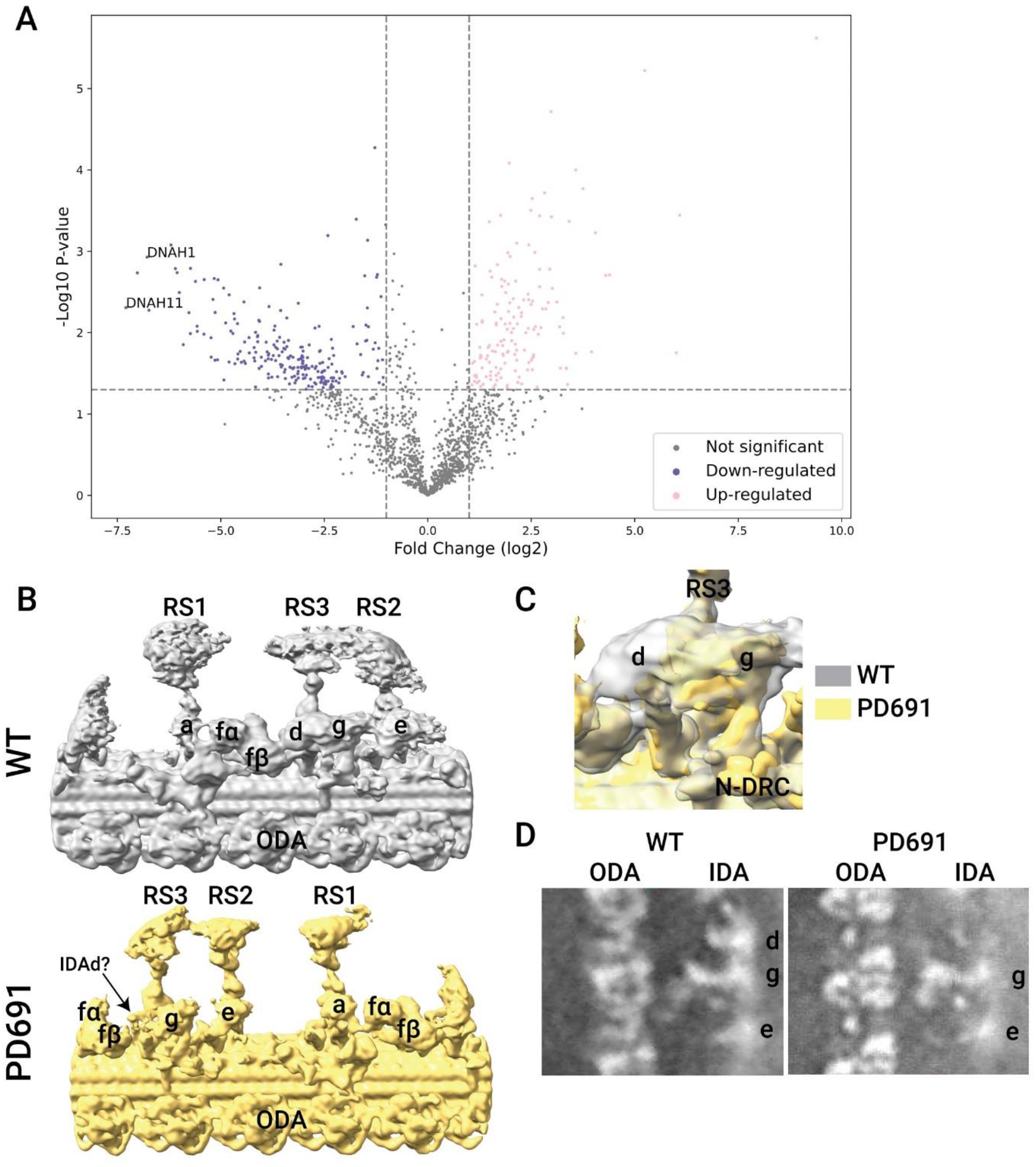
PD691 with genetic defect in DNAH1 and DNAH11. (A) The volcano plot highlighting difference from WT. (B, C, D) Comparison of cryo-ET structure, presenting the overview of the 96nm repeat (B), enlarged view of dynein d (corresponding to DNAH1) (C) and cross section (D), all showing the loss of DNAH1 protein.

### Radial Spokes and Central Pair Apparatus

We systematically evaluated the effect of various mutations on radial spoke (RS) proteins. Two major RS stalk proteins, RSPH3 and RSPH14, decreased in several patients lacking or harboring mutations in dyneins or central pair proteins (Table 3, Suppl. Table 3). Two RS head proteins, RSPH1 and RSPH9, were reduced in most patients, whereas RSH10B was unaffected (Table 3, Suppl. Table 3). Interestingly, patient PD311 (*HYDIN* mutation) exhibited a broader reduction in dyneins than in CP proteins (Suppl. Table 3). None of the three patients with CP protein mutations (PD311, PD343, PD358) showed reductions in RS proteins.

### Intraflagellar Transport (IFT)

While dynein mutations led to reduced IFT components in several patients (Fig. 3A, B), this effect was not uniform across all PCD cases. Among the three patients with *DNAH5* defects (De Ceuninck Van Capelle et al, 2025), PD441 showed a significant decrease in IFT proteins, whereas PD122 and PD480 maintained IFT protein levels comparable to WT (Table 4, Suppl. Table 4). If the loss of dyneins and the decrease in IFT proteins in PD441 are functionally linked, PD122 and PD480 must employ alternative mechanisms that prevent dynein transport or assembly within cilia. Furthermore, among three “unknown” patients (patients presenting PCD-like symptoms but lacking mutations in standard gene panels; Suppl. Table 1A), two (PD403, PD461) showed significant decreases in IFT proteins, whereas one (PD428) did not (Table 4, Suppl. Table 4). This divergence correlated with the loss of dyneins (Table 2, Suppl. Table 2) and RS (Table 3, Suppl. Table 3): mutants with reduced IFT also lacked inner and outer dyneins and RS proteins, further supporting an IFT-mediated loss mechanism.

**Table 4.** Decrease of IFT proteins in 5 patients detected by MS. In each patient, IFT proteins detected significantly lower than in WT are marked with ‘-’. For full datasets of all the patients, refer Suppl. Table 4.

| • Protein names | Gene names | DNAH5 PD122 | DNAH5 PD441 | DNAH5 PD480 | DNAH11 PD442 | DNAH11 PD551 |
| --- | --- | --- | --- | --- | --- | --- |
| Intraflagellar transport protein 122 homolog | IFT122 |  | - |  | - |  |
| Intraflagellar transport protein 140 homolog | IFT140 |  | - |  | - |  |
| Intraflagellar transport protein 172 homolog | IFT172 |  | - |  | - | - |
| Intraflagellar transport protein 20 homolog | IFT20 |  | - |  | - | - |
| Intraflagellar transport protein 22 homolog | IFT22 |  | - |  | - | - |
| Intraflagellar transport protein 25 homolog | IFT25 |  | - |  | - | - |
| Intraflagellar transport protein 27 homolog | IFT27 |  | - |  | - | - |
| Intraflagellar transport protein 43 homolog | IFT43 |  | - |  | - | - |
| Intraflagellar transport protein 46 homolog | IFT46 |  | - |  | - | - |
| Intraflagellar transport protein 52 homolog | IFT52 |  | - |  | - | - |
| Intraflagellar transport protein 56 | IFT56 |  | - |  | - | - |
| Intraflagellar transport protein 57 homolog | IFT57 |  | - |  | - | - |
| Intraflagellar transport protein 70A | IFT70A |  | - |  | - | - |
| Intraflagellar transport protein 70B | IFT70B |  | - |  | - | - |
| Intraflagellar transport protein 74 homolog | IFT74 |  | - |  | - | - |
| Intraflagellar transport protein 80 homolog | IFT80 |  | - |  | - | - |
| Intraflagellar transport protein 81 homolog | IFT81 |  | - |  | - | - |

### ODA Pre- and Post-Powerstroke Conformational States

Reference-free 3D classification using an ODA mask yielded three dominant structural classes (Fig. 5A, B). Between these classes, pre- and post-powerstroke (pre- and post-PS) conformations characteristic of dynein motors could be reliably distinguished (Fig. 5) (Movassagh et al, 2010). Classes were aligned relative to the ODA docking complex, which is fixed onto the doublet microtubule (DC, indicated by black arrows in Fig. 5A).

**Fig. 5:**
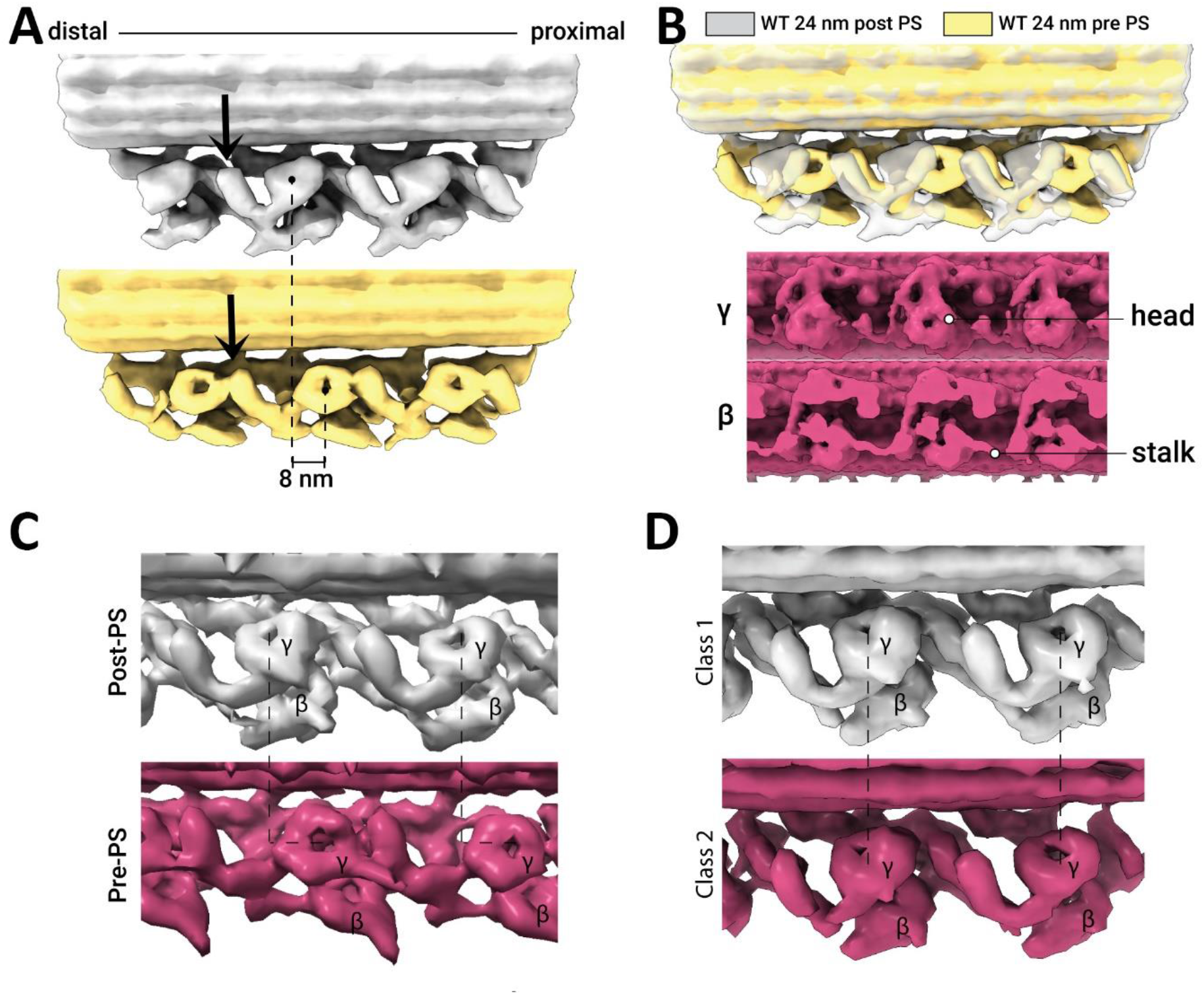
Heterogeneity of outer dynein arm structures corresponding to pre- and post-powerstroke states. **(A)** WT classification map demonstrating ∼8 nm displacement of dynein heads relative to the docking complex (DC, black arrow). **(B)** Superimposition and orthogonal views of pre- and post-powerstroke states. **(C)** Classification of PD442 (*DNAH11* defect) displaying both conformational states. **(D)** Classification of PD435 (*DNAH11* mutation) showing loss of conformational heterogeneity.

Conformational states were assigned based on the position of the AAA+ ring head domain on the heavy chains, the position of the linker, and the reorientation of the stalk. In the pre-PS conformation, the head rotates and shifts ∼8 nm toward the proximal end of the axoneme (Fig. 5A, B), as observed in *Chlamydomonas* ODA (Zimmermann et al, 2023). The linker, which interacts with the AAA4 domain in the post-PS state, predominantly interacts with the AAA2 domain in the pre-PS conformation. Finally, the stalk projects at roughly an angle toward the adjacent doublet microtubule in post-PS, whereas in pre-PS it lies more parallel to the doublets (Fig. 5AB).

Within our WT dataset, 3,685 subtomograms were classified as post-PS (27.6% of total particles) and 2,049 subtomograms as pre-PS (15.4%) (Supplementary Fig. 1). The overall ratio between these states was 1.8:1 (post:pre), indicating a directional bias favoring the post-PS state.

Subtomogram reference-free classification of outer arm dyneins was performed for WT, PD435, PD422, PD222, and PD428. WT and PD442 displayed similar pre/post ratios (Fig. 5A–C). In contrast, PD435 was heavily dominated by the post-PS state (Fig. 5D). While PD442 retained a higher proportion of pre-PS states, PD428 had a higher proportion of post-PS states. These structural distribution differences correlate interestingly with ciliary beating phenotypes: PD428 exhibited partial immotility, PD222 was normal, and PD442 and PD435 exhibited vibrating motions (Table 1).

### DNAH9 Mutation and Sibling Comparison

Patients PD222 and PD272 harbor a *DNAH9* defect in the *cis* allele (Table 1). They are siblings who inherited both mutations from the same mother. Decreases in dyneins were observed by proteomics (Suppl. Fig. 2), though not uniformly across all axonemal dyneins as seen in other patients.

DNAH9 itself was detected in the proteomic data from these patients, and density corresponding to DNAH9 remained visible in cryo-ET maps (Suppl. Fig. 2). This observation can be explained by two possibilities: (1) because the mutation is restricted to the *cis* allele, the density represents wild-type DNAH9 expressed from the *trans* allele; or (2) the analyzed subtomograms selectively originated from the proximal region of cilia (where DNAH11 resides), whereas DNAH9 normally localizes to the distal region.

While both patients showed significant protein reductions compared to WT controls (Suppl. Fig. 2D, E), direct comparison between the two siblings—who share a genetic background—revealed minimal differences (Suppl. Fig. 2F), demonstrating the reproducibility of our experimental strategy.

## Discussion

In this study, we investigated cilia from PCD patients to establish relationships between genetic mutations, quantitative changes in protein composition, and 3D structural architecture. Each patient underwent genetic testing confirming a single primary gene defect, with all other causal genes in the panel remaining intact. While unannotated non-exonic mutations cannot be entirely excluded, our findings indicate that no additional exonic mutations exist within the established PCD gene panel.

### Cascades of Protein Reduction in Human Cilia

Defects in a single gene can cause a broad loss or reduction of multiple secondary proteins in human motile cilia (Fig. 3A, B). In contrast, alteration of ODA components in the *Chlamydomonas oda4-s7* mutant results in far less secondary protein loss (Fig. 3D). This discrepancy likely arises because *oda4-s7* harbors only an N-terminal truncation of dynein, retaining the N-terminal tail that allows assembly of remaining ODA components.

Several mechanisms could explain this secondary cascade loss. The absence or misfolding of a single protein can disrupt complex assembly or prevent loading onto IFT trains at the transition zone, as visualized in *Chlamydomonas* (van den Hoek et al, 2022).

### Differential Phenotypes Among DNAH5 Truncation Mutants

Patients PD122, PD441, and PD480 carry distinct truncation mutations in *DNAH5*, resulting in reduced DNAH5 levels and immotile cilia (De Ceuninck Van Capelle et al, 2025). Proteomics data from all three patients show reductions in inner and outer dyneins (Table 2, Suppl. Table 2) and most RS proteins (Table 3, Suppl. Table 3). However, IFT protein responses varied markedly: PD441 showed a significant decrease in IFT proteins, whereas PD122 and PD480 maintained normal IFT protein levels (Table 4, Suppl. Table 4).

The truncation sites differ significantly: PD441 is truncated at R3000, whereas PD122 and PD480 terminate at Q1450 and D4398, respectively (De Ceuninck Van Capelle et al, 2025). Why should intermediate truncation disrupt IFT components when shorter and longer truncations do not? A plausible explanation is that the remaining fragment in PD441 (residues 1451–3000, spanning the linker to near the C-terminus of AAA5) misfolds, generating steric hindrance or abnormal binding interactions (analogous to domain misfolding effects in p53 (He et al, 2019)). In contrast, the shorter fragment in PD122 and the nearly full-length protein in PD480 may avoid these detrimental interactions.

### Discrepancies Between Proteomics and Cryo-ET Analysis

Certain proteins were detected at normal levels by MS but were absent in cryo-ET reconstructions. In PD222 (Suppl. Fig. 2B), the DNAH3 loci is empty, although proteomics does not show decrease of this protein in PD222 (Table 2, Suppl. Table 2). These proteins may be imported into cilia but fail to dock correctly onto microtubule doublets, rendering them invisible after subtomogram averaging.

Conversely, some proteins appeared structural intact by cryo-ET but were decreased by MS. In PD691, DNAH7 density (dynein e, Fig. 4D) was visible by cryo-ET, despite a decrease in MS intensity relative to WT (Table 2, Suppl. Table 2). In many dynein mutants, RSPH3 decreased in MS (Table 3, Suppl. Table 3), yet radial spokes appeared structurally normal in cryo-ET (Fig. 2C). This discrepancy likely stems from selection bias during particle picking: 96 nm units lacking radial spokes may be systematically omitted during automated or manual picking. Similarly, in PD442, DNAH11 was absent in MS (Table 1), yet full-sized dimeric ODAs were observed in cryo-ET maps (Fig. 5C). This could reflect picking bias or functional compensation by DNAH9, which may expand into the proximal region to occupy vacant DNAH11 sites.

### ODA Assembly in the Absence of Powerstroke States

Patient PD435 (*DNAH11* mutation) displayed morphologically intact ODAs (Fig. 5D) but lacked evidence of powerstroke dynamics, with all subclasses adopting the post-powerstroke conformation. While a minor fraction of pre-powerstroke ODAs below the detection threshold cannot be ruled out, this represents a significant reduction compared to WT cilia (De Ceuninck Van Capelle et al, 2025; Zimmermann et al, 2023).

Because *DNAH11* distribution is restricted to the proximal region of axonemes (Dougherty et al, 2016), DNAH11 represents a minority of total ODA heavy chains. Since our cryo-ET resolution does not distinguish DNAH9 from DNAH11 structurally, averaged maps contain DNAH9/DNAH5 complexes. The finding that unmutated DNAH5 is also trapped in a post-powerstroke state suggests that powerstroke initiation in human ODAs requires coordinated signaling across an interconnected protein network.

### DNAH3 Retention

DNAH3 did not decrease significantly in most patients (with the exception of PD441). DNAH3 corresponds to dynein c in *Chlamydomonas* (Walton et al, 2023). Although cryo-ET density at this locus is weak even in WT human cilia (Fig. 2) (De Ceuninck Van Capelle et al, 2025; Walton et al, 2023), its relative stability across mutants suggests that DNAH3 may be integrated into the axoneme via mechanisms distinct from other dynein arms.

### Radial Spoke Subcomplexes

In several patients, radial spoke stalk proteins (RSPH3 and RSPH14) were reduced while head components like RSH10B remained unaffected (Table 3, Supplementary Table 3). While loss of the spoke stalk typically destabilizes the spoke head (as observed for RSPH1 and RSPH9), RSH10B may be imported into cilia via IFT independently without incorporating into functional axonemal radial spokes. Furthermore, patients with central pair defects (PD311, PD343, PD358) showed no reduction in RS proteins (Table 3, Suppl. Table 3), indicating that while the CP and RS interact functionally (Oda et al, 2014), their assembly and transport pathways are independent.

### Regulation of IFT Entry by Dynein Defects

In multiple dynein mutants (e.g., PD441, PD442, PD551), IFT protein levels were decreased in cilia (Fig. 3A, B; Table 4, Suppl. Table 4). This unexpected finding suggests that cargo availability directly influences IFT train assembly or import. Misfolded or incomplete ciliary cargo complexes may prevent proper assembly with IFT trains (Mali et al, 2021) or block passage through the ciliary transition zone gate.

## Methods

### Cell Culture and Cilia Isolation

WT and patient-derived nasal epithelial cells were cultured, and cilia were isolated as described previously (De Ceuninck Van Capelle et al, 2025). Ethical approval was granted by the Ethics Committees of the University Children’s Hospital, Inselspital Bern, and the Canton of Bern, Switzerland (reference number 2018-02155).

### Cryo-Electron Tomography (Cryo-ET)

Cryo-ET grids were prepared and data acquired on a Titan Krios TEM as described previously (De Ceuninck Van Capelle et al, 2025; Bui & Ishikawa, 2013). Datasets for PD441 were processed using RELION-2 (De Ceuninck Van Capelle et al, 2025; Zimmermann et al, 2023), whereas WT, PD343, and PD691 datasets were processed in RELION-5 (Burt et al, 2024).

Frames and tilt series were pre-processed, reconstructed, and particles were picked manually using Napari (Sofroniew et al, 2026) at 24 nm intervals using filament picking. Filaments were picked directionally from base to tip for each axoneme. Particles were extracted at binning 3 (13.51 Å/pixel box size). Initial 3D references were generated by 3D classification and refined with helical parameters. Subsequent 3D classification resolved azimuthal rotations around the Z-axis. Refinement was performed at progressively lower binning down to binning 1 (4.51 Å/pixel), followed by 200 cycles of tomography refinement, CTF refinement, and Bayesian polishing (Burt et al, 2024).

### Mass Spectrometry Proteomics

Proteomic analysis was conducted as described by Leitner et al. (2014). Peptide spectrum matching was performed in MaxQuant (Cox & Mann, 2008). Statistical significance was assessed using Perseus (Tyanova et al, 2016) via two-sided -tests with 250 permutations.

- Supplementary Table 1A. One typical gene table for PCD diagnosis.
- Supplementary Table 1B. MS proteomics of motile cilia from WT human bronchial epithelial cells. See the attached Excel sheet for the entire list.
- Supplementary Table 2. MS data of 14 patients to show proteins which decreased significantly in the patients’ cilia compared to WT (marked by -).
- Supplementary Table 3 MS of radial spoke (RS) proteins from 14 patients. In each patient, RS proteins detected significantly lower than in WT are marked with ‘-’.
- Supplementary Table 4 Decrease of IFT proteins in 14 patients detected by MS. In each patient, IFT proteins detected significantly lower than in WT are marked with ‘-’.

---

**Supplementary Figure 1.**
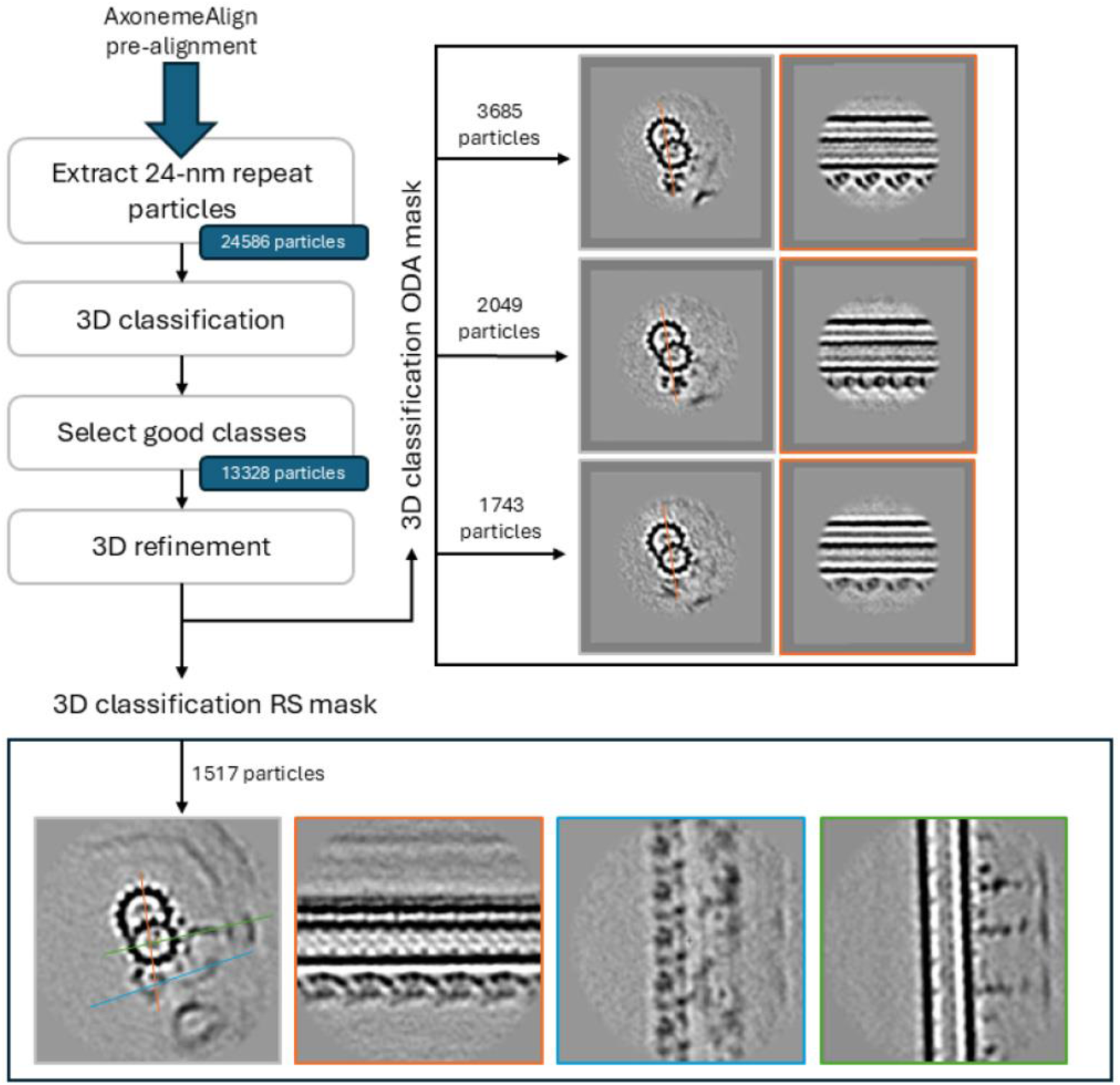
Cryo-ET/STA procedure in this study. Number of particles are for WT.

**Supplementary Figure 2.**
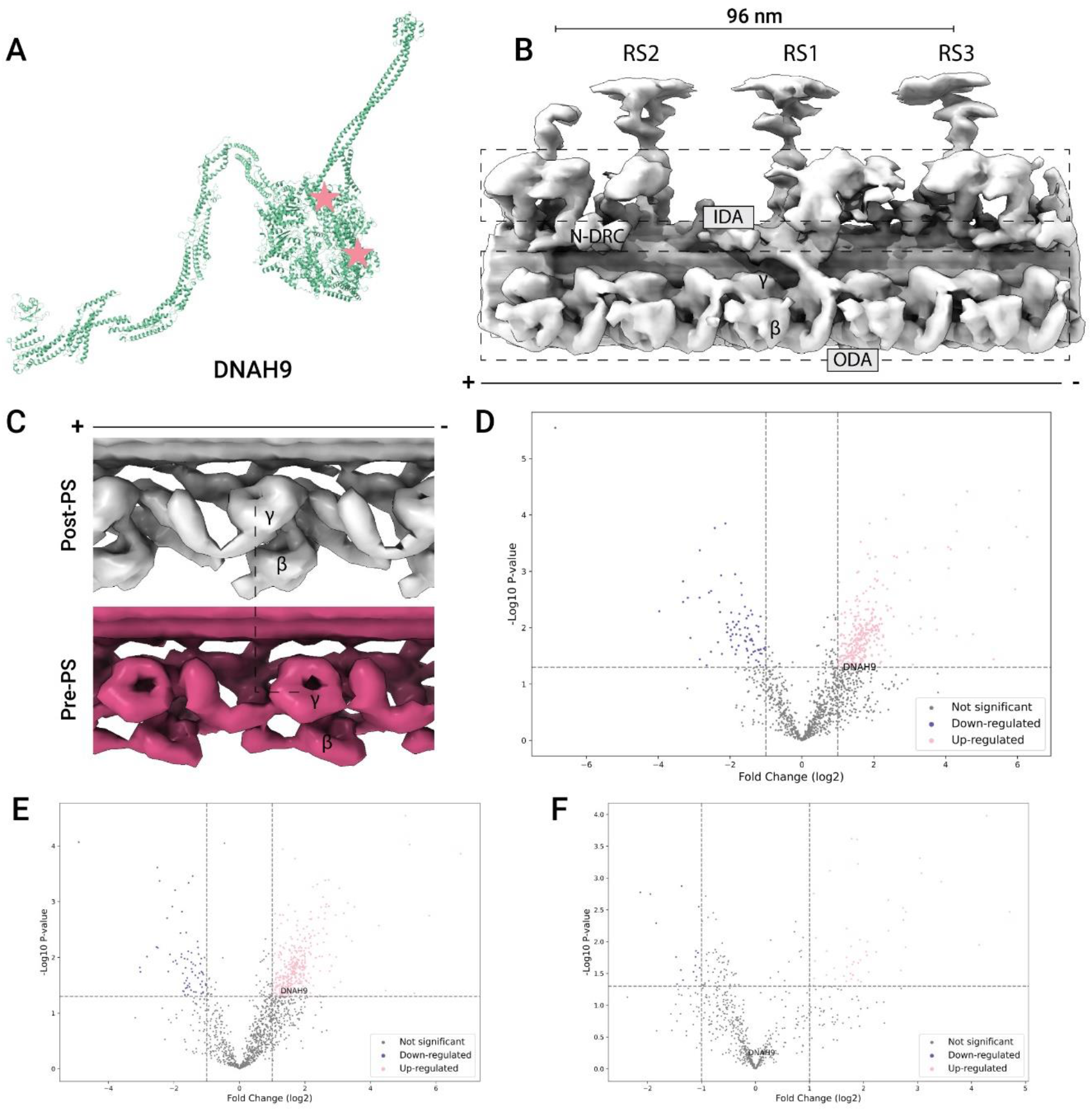
MS and cryo-ET analysis of PD222 and PD272, siblings with cis DNAH9 defect from the same mother. (A) indicates location of mutation in DNAH9 on the 3D atomic model PDB-8J07 (Walton et al, 2023). (B) 3D cryo-ET structure of PD222. (C) Pre- and Post-power stroke structures, detected in ODA from these mutants by cryo-ET and subtomogram classification. (D, E) Comparison of proteomics with WT, shown as volcano plots. (F) Comparison between PD222 and PD272, demonstrating the difference between siblings is small.

