## Supplementary material for "Novel protein networks in motile cilia from human epithelial cells revealed by cryo-ET and proteomics analysis of cilia from PCD patients": SupplTable1A

| Dynein | Assembly/IFT | RS | CP | Ruler | DRC | DC | BB | Ciliogenesis | MIP | Other |
|---|---|---|---|---|---|---|---|---|---|---|
| DNAH1 | DNAAF1 | RSPH1 | HYDIN | CCDC39 | DRC1 | ODAD1 | RPGR | CCNO | CFAP53 | LRRC56 |
| DNAH11 | DNAAF11 | RSHP3 | CFAP74 | CCDC40 | GAS8 | ODAD2 | GAS2L2 | FOXJ1 | ENKUR | LZTFL1 |
| DNAH5 | DNAAF2 | RSPH4A | STK36 | | CCDC65 | ODAD3 | OFD1 | MCIDAS | MNS1 | |
| DNAH7 | DNAAF3 | RSPH9 | CFAP221 | | | ODAD4 | | NEK10 | | |
| DNAH8 | DNAAF4 | DNAJB13 | SPAG17 | | | ODAD5 | | TP73 | | |
| DNAH9 | DNAAF5 | NME5 | SPEF2 | | | | | | | |
| DNAI1 | DNAAF6 | | | | | | | | | |
| DNAI2 | CCDC103 | | | | | | | | | |
| DNAL1 | CFAP300 | | | | | | | | | |
| NME8 | CFAP298 | | | | | | | | | |
| | SPAG1 | | | | | | | | | |
| | TTC12 | | | | | | | | | |
| | ZMYND10 | | | | | | | | | |
| | DAW1 | | | | | | | | | |
| | LRRC6 | | | | | | | | | |

Supplementary Table 1A. One typical gene table for PCD diagnosis. PCD causal genes are classified. BB: basal body. MIP: Microtubule inner protein. DRC: dynein regulatory complex. DC: docking complex.
