## Supplementary material for "Novel protein networks in motile cilia from human epithelial cells revealed by cryo-ET and proteomics analysis of cilia from PCD patients": SupplTable1B

| Protein IDs | Majority protein IDs | Protein names |
| --- | --- | --- |
| A0A024RBG1;Q9NZJ9;( | A0A024RBG1;Q9NZJ! | Diphosphoinositol polyphosphate phosphohydrolase NUD' |
| A0A1B0GU71 | A0A1B0GU71 | Uncharacterized protein CFAP97D2 |
| A0A1B0GV96 | A0A1B0GV96 | CHD9 neighbor protein |
| A0A1B0GVB3 | A0A1B0GVB3 | |
| A0AVF1 | A0AVF1 | Intraflagellar transport protein 56 |
| A0AVT1 | A0AVT1 | Ubiquitin-like modifier-activating enzyme 6 |
| A0MZ66 | A0MZ66 | Shootin-1 |
| A1A4V9 | A1A4V9 | Cilia- and flagella-associated protein 119 |
| A1L0T0 | A1L0T0 | 2-hydroxyacyl-CoA lyase 2 |
| A2IDD5 | A2IDD5 | Coiled-coil domain-containing protein 78 |
| A2VCK2 | A2VCK2 | Doublecortin domain-containing protein 2B |
| A4QMS7 | A4QMS7 | Cilia- and flagella-associated protein 90 |
| A5D8V7 | A5D8V7 | Outer dynein arm-docking complex subunit 3 |
| A5D8W1 | A5D8W1 | Cilia- and flagella-associated protein 69 |
| A6H8M9 | A6H8M9 | Cadherin-related family member 4 |
| A6NCI4 | A6NCI4 | von Willebrand factor A domain-containing protein 3A |
| A6NDG6 | A6NDG6 | Glycerol-3-phosphate phosphatase |
| A6NEE1 | A6NEE1 | Pleckstrin homology domain-containing family D member : |
| A6NFE3 | A6NFE3 | EF-hand calcium-binding domain-containing protein 10 |
| A6NFN9 | A6NFN9 | Protein ANKUB1 |
| A6NFT4 | A6NFT4 | Cilia- and flagella-associated protein 73 |
| A6NHL2 | A6NHL2 | Tubulin alpha chain-like 3 |
| A6NIH7;Q13432 | A6NIH7 | Protein unc-119 homolog B |
| A6NJV1 | A6NJV1 | Ciliary microtubule inner protein 2C |
| A6NL82;Q6ZVS7 | A6NL82 | Cilia- and flagella-associated protein 144 |
| A6NMZ2 | A6NMZ2 | Sentan |
| A6NN90 | A6NN90 | Uncharacterized protein C2orf81 |
| A6NNW6 | A6NNW6 | Enolase 4 |
| A6NNX1 | A6NNX1 | RIIa domain-containing protein 1 |
| A7E2U8 | A7E2U8 | Cilia-and flagella-associated protein 96 |
| A8K855 | A8K855 | EF-hand calcium-binding domain-containing protein 7 |
| A8MQ03 | A8MQ03 | Cysteine-rich tail protein 1 |
| A8MT70 | A8MT70 | Zinc finger B-box domain-containing protein 1 |
| A8MTA8 | A8MTA8 | Ciliary microtubule inner protein 2B |
| A8MYP8 | A8MYP8 | Ciliary microtubule associated protein 1B |
| E5RIL1;B0FP48 | E5RIL1;B0FP48 | Uroplakin-3b-like protein 2;Uroplakin-3b-like protein 1 |
| B1AJZ9 | B1AJZ9 | Forkhead-associated domain-containing protein 1 |
| B1ANS9 | B1ANS9 | WD repeat-containing protein 64 |
| B2RC85 | B2RC85 | Radial spoke head 10 homolog B2 |
| Q99613;B5ME19 | Q99613;B5ME19 | Eukaryotic translation initiation factor 3 subunit C;Eukaryot |
| D6REC4 | D6REC4 | Cilia- and flagella-associated protein 99 |
| E9PAV3;Q13765;Q9BZI | E9PAV3;Q13765 | Nascent polypeptide-associated complex subunit alpha, m |
| H3BMG3 | H3BMG3 | Small lysine-rich protein 1 |
| H3BQB6 | H3BQB6 | Stathmin domain-containing protein 1 |
| H3BRN8 | H3BRN8 | Piercer of microtubule wall 2 protein |
| H7BZ55 | H7BZ55 | Ciliary rootlet coiled-coil protein 2 |

| | | |
|---|---|---|
| K7EJ46 | K7EJ46 | Small integral membrane protein 22 |
| M0R2J8 | M0R2J8 | Doublecortin domain-containing protein 1 |
| O00139 | O00139 | Kinesin-like protein KIF2A |
| O00151 | O00151 | PDZ and LIM domain protein 1 |
| O00154 | O00154 | Cytosolic acyl coenzyme A thioester hydrolase |
| O00159 | O00159 | Unconventional myosin-Ic |
| O00161 | O00161 | Synaptosomal-associated protein 23 |
| O00182;Q6DKI2;Q3B8I | O00182 | Galectin-9 |
| O00194;P51159 | O00194 | Ras-related protein Rab-27B |
| O00203;Q13367 | O00203 | AP-3 complex subunit beta-1 |
| O00204 | O00204 | Sulfotransferase 2B1 |
| O00217 | O00217 | NADH dehydrogenase [ubiquinone] iron-sulfur protein 8, m |
| O00231 | O00231 | 26S proteasome non-ATPase regulatory subunit 11 |
| O00232 | O00232 | 26S proteasome non-ATPase regulatory subunit 12 |
| O00233 | O00233 | 26S proteasome non-ATPase regulatory subunit 9 |
| O00244 | O00244 | Copper transport protein ATOX1 |
| O00264 | O00264 | Membrane-associated progesterone receptor component [illegible] |
| O00273 | O00273 | DNA fragmentation factor subunit alpha |
| O00299 | O00299 | Chloride intracellular channel protein 1 |
| O00391 | O00391 | Sulfhydryl oxidase 1 |
| O00410;O60518 | O00410 | Importin-5 |
| O00425 | O00425 | Insulin-like growth factor 2 mRNA-binding protein 3 |
| O00429 | O00429 | Dynamin-1-like protein |
| O00442 | O00442 | RNA 3-terminal phosphate cyclase |
| O00468 | O00468 | Agrin;Agrin N-terminal 110 kDa subunit;Agrin C-terminal 11 |
| O00483 | O00483 | Cytochrome c oxidase subunit NDUFA4 |
| O00487 | O00487 | 26S proteasome non-ATPase regulatory subunit 14 |
| O00505 | O00505 | Importin subunit alpha-4 |
| O00534 | O00534 | von Willebrand factor A domain-containing protein 5A |
| O00560 | O00560 | Syntenin-1 |
| O00567 | O00567 | Nucleolar protein 56 |
| O00571;O15523;Q9N[illegible] | O00571;O15523 | ATP-dependent RNA helicase DDX3X;ATP-dependent RNA [illegible] |
| O00584 | O00584 | Ribonuclease T2 |
| O00625 | O00625 | Pirin |
| O00743 | O00743 | Serine/threonine-protein phosphatase 6 catalytic subunit;[illegible] |
| O00748;Q5XG92;P223 | O00748 | Cocaine esterase |
| O00757 | O00757 | Fructose-1,6-bisphosphatase isozyme 2 |
| O00764 | O00764 | Pyridoxal kinase |
| O14545 | O14545 | TRAF-type zinc finger domain-containing protein 1 |
| O14561 | O14561 | Acyl carrier protein, mitochondrial |
| O14579 | O14579 | Coatomer subunit epsilon |
| O14645 | O14645 | Axonemal dynein light intermediate polypeptide 1 |
| O14737 | O14737 | Programmed cell death protein 5 |
| O14744 | O14744 | Protein arginine N-methyltransferase 5;Protein arginine N-[illegible] |
| O14745 | O14745 | Na(+)/H(+) exchange regulatory cofactor NHE-RF1 |
| O14757 | O14757 | Serine/threonine-protein kinase Chk1 |
| O14772 | O14772 | Fucose-1-phosphate guanylyltransferase |

| | | |
|---|---|---|
| O14773 | O14773 | Tripeptidyl-peptidase 1 |
| O14818;Q8TAA3 | O14818 | Proteasome subunit alpha type-7 |
| O14879 | O14879 | Interferon-induced protein with tetratricopeptide repeats 3 |
| O14880 | O14880 | Glutathione S-transferase 3, mitochondrial |
| O14896 | O14896 | Interferon regulatory factor 6 |
| O14907 | O14907 | Tax1-binding protein 3 |
| O14908 | O14908 | PDZ domain-containing protein GIPC1 |
| O14933 | O14933 | Ubiquitin/ISG15-conjugating enzyme E2 L6 |
| O14949 | O14949 | Cytochrome b-c1 complex subunit 8 |
| P19105;O14950;P2484 | P19105;O14950;P24 | Myosin regulatory light chain 12A;Myosin regulatory light ch |
| O14964 | O14964 | Hepatocyte growth factor-regulated tyrosine kinase substra |
| O14974 | O14974 | Protein phosphatase 1 regulatory subunit 12A |
| O14975 | O14975 | Long-chain fatty acid transport protein 2 |
| O14980 | O14980 | Exportin-1 |
| O15020 | O15020 | Spectrin beta chain, non-erythrocytic 2 |
| O15066;O14782;Q9BV | O15066 | Kinesin-like protein KIF3B;Kinesin-like protein KIF3B, N-ter |
| O15067 | O15067 | Phosphoribosylformylglycinamidine synthase |
| O15143 | O15143 | Actin-related protein 2/3 complex subunit 1B |
| O15144 | O15144 | Actin-related protein 2/3 complex subunit 2 |
| O15145 | O15145 | Actin-related protein 2/3 complex subunit 3 |
| O15162 | O15162 | Phospholipid scramblase 1 |
| O15195 | O15195 | Villin-like protein |
| O15212 | O15212 | Prefoldin subunit 6 |
| O15231 | O15231 | Zinc finger protein 185 |
| O15254 | O15254 | Peroxisomal acyl-coenzyme A oxidase 3 |
| O15260 | O15260 | Surfeit locus protein 4 |
| O15305 | O15305 | Phosphomannomutase 2 |
| O15393 | O15393 | Transmembrane protease serine 2;Transmembrane protea |
| O15427 | O15427 | Monocarboxylate transporter 4 |
| O15431 | O15431 | High affinity copper uptake protein 1;Truncated CTR1 form |
| O15484 | O15484 | Calpain-5 |
| O15511 | O15511 | Actin-related protein 2/3 complex subunit 5 |
| Q6A1A2;O15530 | Q6A1A2;O15530 | Putative 3-phosphoinositide-dependent protein kinase 2;3- |
| O15533 | O15533 | Tapasin |
| O43143;O60231;Q145 | O43143 | ATP-dependent RNA helicase DHX15 |
| O43175 | O43175 | D-3-phosphoglycerate dehydrogenase |
| O43237 | O43237 | Cytoplasmic dynein 1 light intermediate chain 2 |
| O43240 | O43240 | Kallikrein-10 |
| O43242 | O43242 | 26S proteasome non-ATPase regulatory subunit 3 |
| O43252;O95340 | O43252;O95340 | Bifunctional 3-phosphoadenosine 5-phosphosulfate synth |
| O43324 | O43324 | Eukaryotic translation elongation factor 1 epsilon-1 |
| O43390 | O43390 | Heterogeneous nuclear ribonucleoprotein R |
| O43396 | O43396 | Thioredoxin-like protein 1 |
| O43399 | O43399 | Tumor protein D54 |
| O43447 | O43447 | Peptidyl-prolyl cis-trans isomerase H |
| O43488;Q8NHP1;O951 | O43488 | Aflatoxin B1 aldehyde reductase member 2 |
| O43490 | O43490 | Prominin-1 |

| | | |
|---|---|---|
| O43504 | O43504 | Ragulator complex protein LAMTOR5 |
| O43583 | O43583 | Density-regulated protein |
| O43598 | O43598 | 5-hydroxymethyl-dUMP N-hydrolase |
| O43617 | O43617 | Trafficking protein particle complex subunit 3 |
| O43633 | O43633 | Charged multivesicular body protein 2a |
| O43653 | O43653 | Prostate stem cell antigen |
| O43681 | O43681 | ATPase GET3 |
| O43704 | O43704 | Sulfotransferase 1B1 |
| O43707;Q08043 | O43707 | Alpha-actinin-4 |
| O43708 | O43708 | Maleylacetoacetate isomerase |
| O43747 | O43747 | AP-1 complex subunit gamma-1 |
| O43760 | O43760 | Synaptogyrin-2 |
| O43765 | O43765 | Small glutamine-rich tetratricopeptide repeat-containing p |
| O43768 | O43768 | Alpha-endosulfine |
| O43776 | O43776 | Asparagine--tRNA ligase, cytoplasmic |
| O43795 | O43795 | Unconventional myosin-Ib |
| O43813 | O43813 | Glutathione S-transferase LANCL1 |
| O43837 | O43837 | Isocitrate dehydrogenase [NAD] subunit beta, mitochondri |
| O43847 | O43847 | Nardilysin |
| O43852 | O43852 | Calumenin |
| O43865;Q96HN2 | O43865;Q96HN2 | S-adenosylhomocysteine hydrolase-like protein 1;Adenosy |
| O60218 | O60218 | Aldo-keto reductase family 1 member B10 |
| O60229 | O60229 | Kalirin |
| O60308 | O60308 | Centrosomal protein of 104 kDa |
| O60437 | O60437 | Periplakin |
| O60493 | O60493 | Sorting nexin-3 |
| O60547 | O60547 | GDP-mannose 4,6 dehydratase |
| O60610 | O60610 | Protein diaphanous homolog 1 |
| O60635 | O60635 | Tetraspanin-1 |
| O60664 | O60664 | Perilipin-3 |
| O60701 | O60701 | UDP-glucose 6-dehydrogenase |
| O60716 | O60716 | Catenin delta-1 |
| O60749 | O60749 | Sorting nexin-2 |
| O60763;REV__Q5VZ89 | O60763 | General vesicular transport factor p115 |
| O60784 | O60784 | Target of Myb1 membrane trafficking protein |
| Q99880;Q99879;Q998 | Q99880;Q99879;Q9 | Histone H2B type 1-L;Histone H2B type 1-M;Histone H2B t |
| O60869 | O60869 | Endothelial differentiation-related factor 1 |
| O60884 | O60884 | DnaJ homolog subfamily A member 2 |
| O60888 | O60888 | Protein CutA |
| O60906 | O60906 | Sphingomyelin phosphodiesterase 2 |
| O75083 | O75083 | WD repeat-containing protein 1 |
| O75131;Q96FN4;Q8IYJ | O75131 | Copine-3 |
| O75190;Q8WWF6;Q8N | O75190 | DnaJ homolog subfamily B member 6 |
| O75223 | O75223 | Gamma-glutamylcyclotransferase |
| O75306 | O75306 | NADH dehydrogenase [ubiquinone] iron-sulfur protein 2, m |
| O75323 | O75323 | Protein NipSnap homolog 2 |
| O75340 | O75340 | Programmed cell death protein 6 |

| | | |
|---|---|---|
| O75347 | O75347 | Tubulin-specific chaperone A |
| O75348;O95670 | O75348 | V-type proton ATPase subunit G 1 |
| O75351 | O75351 | Vacuolar protein sorting-associated protein 4B |
| O75352 | O75352 | Mannose-P-dolichol utilization defect 1 protein |
| O75363 | O75363 | Breast carcinoma-amplified sequence 1 |
| O75367;Q9P0M6 | O75367 | Core histone macro-H2A.1 |
| O75368 | O75368 | Adapter SH3BGRL |
| O75369 | O75369 | Filamin-B |
| O75386 | O75386 | Tubby-related protein 3 |
| O75390 | O75390 | Citrate synthase, mitochondrial |
| O75396 | O75396 | Vesicle-trafficking protein SEC22b |
| O75436 | O75436 | Vacuolar protein sorting-associated protein 26A |
| O75438 | O75438 | NADH dehydrogenase [ubiquinone] 1 beta subcomplex sub |
| O75487;Q9Y625 | O75487 | Glypican-4;Secreted glypican-4 |
| O75494 | O75494 | Serine/arginine-rich splicing factor 10 |
| O75506 | O75506 | Heat shock factor-binding protein 1 |
| O75531 | O75531 | Barrier-to-autointegration factor;Barrier-to-autointegration |
| O75602 | O75602 | Sperm-associated antigen 6 |
| O75604 | O75604 | Ubiquitin carboxyl-terminal hydrolase 2 |
| O75608 | O75608 | Acyl-protein thioesterase 1 |
| O75663 | O75663 | TIP41-like protein |
| O75695 | O75695 | Protein XRP2 |
| O75787 | O75787 | Renin receptor;Renin receptor N-terminal fragment;Renin r |
| O75800 | O75800 | Zinc finger MYND domain-containing protein 10 |
| O75828 | O75828 | Carbonyl reductase [NADPH] 3 |
| O75832 | O75832 | 26S proteasome non-ATPase regulatory subunit 10 |
| O75841 | O75841 | Uroplakin-1b |
| O75874 | O75874 | Isocitrate dehydrogenase [NADP] cytoplasmic |
| O75884 | O75884 | Serine hydrolase RBBP9 |
| O75891;Q3SY69 | O75891 | Cytosolic 10-formyltetrahydrofolate dehydrogenase |
| O75911 | O75911 | Short-chain dehydrogenase/reductase 3 |
| O75915 | O75915 | PRA1 family protein 3 |
| O75947 | O75947 | ATP synthase subunit d, mitochondrial |
| O75955 | O75955 | Flotillin-1 |
| O75964 | O75964 | ATP synthase subunit g, mitochondrial |
| O76003 | O76003 | Glutaredoxin-3 |
| O76054 | O76054 | SEC14-like protein 2 |
| O94760 | O94760 | N(G),N(G)-dimethylarginine dimethylaminohydrolase 1 |
| O94811 | O94811 | Tubulin polymerization-promoting protein |
| O94826 | O94826 | Mitochondrial import receptor subunit TOM70 |
| O94832 | O94832 | Unconventional myosin-Id |
| O94903 | O94903 | Pyridoxal phosphate homeostasis protein |
| O94905 | O94905 | Erlin-2 |
| O94967 | O94967 | WD repeat-containing protein 47 |
| O94979 | O94979 | Protein transport protein Sec31A |
| O95084 | O95084 | Serine protease 23 |
| O95292 | O95292 | Vesicle-associated membrane protein-associated protein I |

| | | |
|---|---|---|
| O95299 | O95299 | NADH dehydrogenase [ubiquinone] 1 alpha subcomplex su |
| O95336 | O95336 | 6-phosphogluconolactonase |
| O95352 | O95352 | Ubiquitin-like modifier-activating enzyme ATG7 |
| O95361;Q309B1 | O95361 | Tripartite motif-containing protein 16 |
| O95372 | O95372 | Acyl-protein thioesterase 2 |
| O95373;O15397 | O95373 | Importin-7 |
| O95379 | O95379 | Tumor necrosis factor alpha-induced protein 8 |
| O95394 | O95394 | Phosphoacetylglucosamine mutase |
| O95433 | O95433 | Activator of 90 kDa heat shock protein ATPase homolog 1 |
| O95447 | O95447 | Lebercilin-like protein |
| O95571 | O95571 | Persulfide dioxygenase ETHE1, mitochondrial |
| O95573;O60488 | O95573 | Fatty acid CoA ligase Acsl3 |
| O95716 | O95716 | Ras-related protein Rab-3D |
| O95747 | O95747 | Serine/threonine-protein kinase OSR1 |
| O95757 | O95757 | Heat shock 70 kDa protein 4L |
| O95782 | O95782 | AP-2 complex subunit alpha-1 |
| O95786 | O95786 | Antiviral innate immune response receptor RIG-I |
| O95833 | O95833 | Chloride intracellular channel protein 3 |
| O95834 | O95834 | Echinoderm microtubule-associated protein-like 2 |
| O95837;P19086;Q031 | O95837 | Guanine nucleotide-binding protein subunit alpha-14 |
| O95848 | O95848 | Uridine diphosphate glucose pyrophosphatase NUDT14 |
| O95861 | O95861 | 3(2),5-bisphosphate nucleotidase 1 |
| O95865 | O95865 | Putative hydrolase DDAH2 |
| O95881 | O95881 | Thioredoxin domain-containing protein 12 |
| O95989 | O95989 | Diphosphoinositol polyphosphate phosphohydrolase 1 |
| O95994 | O95994 | Anterior gradient protein 2 homolog |
| O95995 | O95995 | Dynein regulatory complex subunit 4 |
| O96000 | O96000 | NADH dehydrogenase [ubiquinone] 1 beta subcomplex sub |
| O96008 | O96008 | Mitochondrial import receptor subunit TOM40 homolog |
| O96015 | O96015 | Dynein axonemal light chain 4 |
| P00167 | P00167 | Cytochrome b5 |
| P00326;P07327;P0032 | P00326 | Alcohol dehydrogenase 1C |
| P00338;Q6ZMR3 | P00338 | L-lactate dehydrogenase A chain |
| P00352 | P00352 | Aldehyde dehydrogenase 1A1 |
| P00367;P49448 | P00367;P49448 | Glutamate dehydrogenase 1, mitochondrial;Glutamate del |
| P00387 | P00387 | NADH-cytochrome b5 reductase 3 |
| P00390 | P00390 | Glutathione reductase, mitochondrial |
| P00403 | P00403 | Cytochrome c oxidase subunit 2 |
| P00441 | P00441 | Superoxide dismutase [Cu-Zn] |
| P00450 | P00450 | Ceruloplasmin |
| P00491 | P00491 | Purine nucleoside phosphorylase |
| P00492;Q9NRG1 | P00492 | Hypoxanthine-guanine phosphoribosyltransferase |
| P00505 | P00505 | Aspartate aminotransferase, mitochondrial |
| P00558;P07205 | P00558 | Phosphoglycerate kinase 1 |
| P00568;Q9Y6K8 | P00568 | Adenylate kinase isoenzyme 1 |
| P00749 | P00749 | Urokinase-type plasminogen activator;Urokinase-type plas |
| P00751 | P00751 | Complement factor B;Complement factor B Ba fragment;C |

| | | |
|---|---|---|
| P00846 | P00846 | ATP synthase subunit a |
| P00966 | P00966 | Argininosuccinate synthase |
| P01009 | P01009 | Alpha-1-antitrypsin;Short peptide from AAT |
| P01024;CON__Q2UVX4 | P01024 | Complement C3;Complement C3 beta chain;C3-beta-c;Co |
| P01034 | P01034 | Cystatin-C |
| P01040 | P01040 | Cystatin-A;Cystatin-A, N-terminally processed |
| P01112 | P01112 | GTPase HRas;GTPase HRas, N-terminally processed |
| P01116 | P01116 | GTPase KRas;GTPase KRas, N-terminally processed |
| P01308;F8WCM5 | P01308;F8WCM5 | Insulin;Insulin B chain;Insulin A chain;Insulin, isoform 2 |
| P01833 | P01833 | Polymeric immunoglobulin receptor;Secretory component |
| P01903;P01906 | P01903;P01906 | HLA class II histocompatibility antigen, DR alpha chain;HLA |
| P01911 | P01911 | HLA class II histocompatibility antigen, DRB1 beta chain |
| P02144 | P02144 | Myoglobin |
| P02545 | P02545 | Prelamin-A/C;Lamin-A/C |
| P02750 | P02750 | Leucine-rich alpha-2-glycoprotein |
| P02751 | P02751 | Fibronectin;Anastellin;Ugl-Y1;Ugl-Y2;Ugl-Y3 |
| P02787 | P02787 | Serotransferrin |
| P02790 | P02790 | Hemopexin |
| P02794 | P02794 | Ferritin heavy chain;Ferritin heavy chain, N-terminally proc |
| P03973 | P03973 | Antileukoproteinase |
| P04040 | P04040 | Catalase |
| P04066 | P04066 | Tissue alpha-L-fucosidase |
| P04075;P05062 | P04075 | Fructose-bisphosphate aldolase A |
| P04080 | P04080 | Cystatin-B |
| P04083 | P04083 | Annexin A1;Annexin Ac2-26 |
| P04179 | P04179 | Superoxide dismutase [Mn], mitochondrial |
| P04181 | P04181 | Ornithine aminotransferase, mitochondrial;Ornithine amin |
| P04406;O14556 | P04406 | Glyceraldehyde-3-phosphate dehydrogenase |
| P04424 | P04424 | Argininosuccinate lyase |
| P04439;P01893;P3051 | P04439;P01893 | HLA class I histocompatibility antigen, A alpha chain;Putati |
| P04632 | P04632 | Calpain small subunit 1 |
| P04792 | P04792 | Heat shock protein beta-1 |
| P04843 | P04843 | Dolichyl-diphosphooligosaccharide--protein glycosyltransf |
| P04844 | P04844 | Dolichyl-diphosphooligosaccharide--protein glycosyltransf |
| P04899 | P04899 | Guanine nucleotide-binding protein G(i) subunit alpha-2 |
| P05023;P13637;P5099 | P05023 | Sodium/potassium-transporting ATPase subunit alpha-1 |
| P05026 | P05026 | Sodium/potassium-transporting ATPase subunit beta-1 |
| P05090 | P05090 | Apolipoprotein D |
| P05091 | P05091 | Aldehyde dehydrogenase, mitochondrial |
| P05109 | P05109 | Protein S100-A8 |
| P05120 | P05120 | Plasminogen activator inhibitor 2 |
| P05141;Q9H0C2 | P05141 | ADP/ATP translocase 2;ADP/ATP translocase 2, N-terminall |
| P05161 | P05161 | Ubiquitin-like protein ISG15 |
| P05186 | P05186 | Alkaline phosphatase, tissue-nonspecific isozyme |
| P05386 | P05386 | Large ribosomal subunit protein P1 |
| P05387 | P05387 | Large ribosomal subunit protein P2 |
| P05388;Q8NHW5 | P05388;Q8NHW5 | Large ribosomal subunit protein uL10;Putative ribosomal p |

| | | |
|---|---|---|
| P05455 | P05455 | Lupus La protein |
| P05783;CON__H-INV:H | P05783 | Keratin, type I cytoskeletal 18 |
| P05976;P08590;P1282 | P05976;P08590 | Myosin light chain 1/3, skeletal muscle isoform;Myosin ligh |
| P06396;CON__Q3SX14 | P06396 | Gelsolin |
| P06454 | P06454 | Prothymosin alpha;Prothymosin alpha, N-terminally proces |
| P06576 | P06576 | ATP synthase subunit beta, mitochondrial |
| P06702 | P06702 | Protein S100-A9 |
| P06703 | P06703 | Protein S100-A6 |
| P06730;A6NMX2 | P06730 | Eukaryotic translation initiation factor 4E |
| P06731;P40198;P3199 | P06731 | Carcinoembryonic antigen-related cell adhesion molecule |
| P06733 | P06733 | Alpha-enolase |
| P06737 | P06737 | Glycogen phosphorylase, liver form |
| P06744;Q8N196 | P06744 | Glucose-6-phosphate isomerase |
| P06748 | P06748 | Nucleophosmin |
| P06753;P09493 | P06753 | Tropomyosin alpha-3 chain |
| P06756 | P06756 | Integrin alpha-V;Integrin alpha-V heavy chain;Integrin alpha |
| P06865 | P06865 | Beta-hexosaminidase subunit alpha |
| Q16778;P33778;P2352 | Q16778;P33778;P23 | Histone H2B type 2-E;Histone H2B type 1-B;Histone H2B ty |
| P07099 | P07099 | Epoxide hydrolase 1 |
| P07108 | P07108 | Acyl-CoA-binding protein |
| P07195;P07864 | P07195 | L-lactate dehydrogenase B chain |
| P07203 | P07203 | Glutathione peroxidase 1 |
| P07237 | P07237 | Protein disulfide-isomerase |
| P07305 | P07305 | Histone H1.0;Histone H1.0, N-terminally processed |
| P07311 | P07311 | Acylphosphatase-1 |
| P07339 | P07339 | Cathepsin D;Cathepsin D light chain;Cathepsin D heavy ch |
| P07355;A6NMY6 | P07355;A6NMY6 | Annexin A2;Putative annexin A2-like protein |
| P07384 | P07384 | Calpain-1 catalytic subunit |
| P07437;A6NNZ2;Q9H4 | P07437 | Tubulin beta chain |
| P07476 | P07476 | Involucrin |
| P07686 | P07686 | Beta-hexosaminidase subunit beta;Beta-hexosaminidase s |
| P07737;CON__P02584 | P07737 | Profilin-1 |
| P07738 | P07738 | Bisphosphoglycerate mutase |
| P07741 | P07741 | Adenine phosphoribosyltransferase |
| P07814 | P07814 | Bifunctional glutamate/proline--tRNA ligase;Glutamate--tR |
| P07858 | P07858 | Cathepsin B;Cathepsin B light chain;Cathepsin B heavy cha |
| P07900;Q58FG0;Q145 | P07900 | Heat shock protein HSP 90-alpha |
| P07910;P0DMR1;O608 | P07910;P0DMR1;O60 | Heterogeneous nuclear ribonucleoproteins C1/C2;Heterog |
| P07919;A0A096LP55 | P07919 | Cytochrome b-c1 complex subunit 6, mitochondrial |
| P07947;P06241 | P07947 | Tyrosine-protein kinase Yes |
| P07948;P08631 | P07948 | Tyrosine-protein kinase Lyn |
| P07954 | P07954 | Fumarate hydratase, mitochondrial |
| P08118 | P08118 | Beta-microseminoprotein |
| P08134;P62745 | P08134 | Rho-related GTP-binding protein RhoC |
| P08174 | P08174 | Complement decay-accelerating factor |
| P08195 | P08195 | Amino acid transporter heavy chain SLC3A2 |
| P08238;Q58FF7 | P08238 | Heat shock protein HSP 90-beta |

| | | |
|---|---|---|
| P08263;Q7RTV2 | P08263 | Glutathione S-transferase A1;Glutathione S-transferase A1 |
| P08559;P29803 | P08559 | Pyruvate dehydrogenase E1 component subunit alpha, son |
| P08574 | P08574 | Cytochrome c1, heme protein, mitochondrial |
| P08603;Q03591;Q029 | P08603 | Complement factor H |
| P08670 | P08670 | Vimentin |
| P08708 | P08708 | Small ribosomal subunit protein eS17 |
| P08754;P11488;P1908 | P08754 | Guanine nucleotide-binding protein G(i) subunit alpha-3 |
| P08758 | P08758 | Annexin A5 |
| P08865 | P08865 | Small ribosomal subunit protein uS2 |
| P09104;P13929 | P09104 | Gamma-enolase |
| P09210 | P09210 | Glutathione S-transferase A2 |
| P09211 | P09211 | Glutathione S-transferase P |
| P09238;P08254;P2289 | P09238 | Stromelysin-2 |
| P09341;P19876;P1987 | P09341 | Growth-regulated alpha protein;GRO-alpha(4-73);GRO-alp |
| P09382 | P09382 | Galectin-1 |
| P09417 | P09417 | Dihydropteridine reductase |
| P09429;B2RPK0;P2658 | P09429;B2RPK0 | High mobility group protein B1;Putative high mobility group |
| P09455 | P09455 | Retinol-binding protein 1 |
| P09467 | P09467 | Fructose-1,6-bisphosphatase 1 |
| P09488 | P09488 | Glutathione S-transferase Mu 1 |
| P09496 | P09496 | Clathrin light chain A |
| P09525 | P09525 | Annexin A4 |
| P09543 | P09543 | 2,3-cyclic-nucleotide 3-phosphodiesterase |
| P09622 | P09622 | Dihydrolipoyl dehydrogenase, mitochondrial |
| P09651;A0A2R8Y4L2;C | P09651;A0A2R8Y4L2 | Heterogeneous nuclear ribonucleoprotein A1;Heterogenec |
| P09668 | P09668 | Pro-cathepsin H;Cathepsin H mini chain;Cathepsin H;Cath |
| P09669 | P09669 | Cytochrome c oxidase subunit 6C |
| P09758 | P09758 | Tumor-associated calcium signal transducer 2 |
| P09914;Q5T764 | P09914 | Interferon-induced protein with tetratricopeptide repeats 1 |
| P09919 | P09919 | Granulocyte colony-stimulating factor |
| P09960 | P09960 | Leukotriene A-4 hydrolase |
| P09972 | P09972 | Fructose-bisphosphate aldolase C |
| Q71UI9;P0C0S5 | Q71UI9;P0C0S5 | Histone H2A.V;Histone H2A.Z |
| Q99878;Q96KK5;Q9BT | Q99878;Q96KK5;Q9I | Histone H2A type 1-J;Histone H2A type 1-H;Histone H2A.J; |
| P0C7W0 | P0C7W0 | Proline-rich protein 29 |
| P0C870 | P0C870 | Bifunctional peptidase and (3S)-lysyl hydroxylase JMJD7 |
| P0C875 | P0C875 | Protein FAM228B |
| P0CG21 | P0CG21 | NHL-repeat-containing protein 4 |
| P0DJI8;P0DJI9 | P0DJI8;P0DJI9 | Serum amyloid A-1 protein;Amyloid protein A;Serum amylo |
| P0DME0;Q01105 | P0DME0;Q01105 | Protein SETSIP;Protein SET |
| P0DMN0;P0DMM9 | P0DMN0;P0DMM9 | Sulfotransferase 1A4;Sulfotransferase 1A3 |
| P0DMV9;P0DMV8 | P0DMV9;P0DMV8 | Heat shock 70 kDa protein 1B;Heat shock 70 kDa protein 1 |
| P0DP25;P0DP24;P0DP | P0DP25;P0DP24;P0D | Calmodulin-3;Calmodulin-2;Calmodulin-1 |
| P0DP57 | P0DP57 | Secreted Ly-6/uPAR domain-containing protein 2 |
| P10155 | P10155 | RNA-binding protein RO60 |
| P10253 | P10253 | Lysosomal alpha-glucosidase;76 kDa lysosomal alpha-gluc |
| P10321 | P10321 | HLA class I histocompatibility antigen, C alpha chain |

| | | |
|---|---|---|
| P10412;P16402;P2249 | P10412;P16402 | Histone H1.4;Histone H1.3 |
| P10515 | P10515 | Dihydrolipoyllysine-residue acetyltransferase component c |
| P10599 | P10599 | Thioredoxin |
| P10606 | P10606 | Cytochrome c oxidase subunit 5B, mitochondrial |
| P10619 | P10619 | Lysosomal protective protein;Lysosomal protective protein |
| P10620 | P10620 | Microsomal glutathione S-transferase 1 |
| P10644;P31321 | P10644 | cAMP-dependent protein kinase type I-alpha regulatory sub |
| P10768 | P10768 | S-formylglutathione hydrolase |
| P10809 | P10809 | 60 kDa heat shock protein, mitochondrial |
| P10909 | P10909 | Clusterin;Clusterin beta chain;Clusterin alpha chain |
| P11021 | P11021 | Endoplasmic reticulum chaperone BiP |
| P11117 | P11117 | Lysosomal acid phosphatase |
| P11142 | P11142 | Heat shock cognate 71 kDa protein |
| P11172 | P11172 | Uridine 5-monophosphate synthase;Orotate phosphoribos |
| P11177 | P11177 | Pyruvate dehydrogenase E1 component subunit beta, mito |
| P11182 | P11182 | Lipoamide acyltransferase component of branched-chain a |
| P11216 | P11216 | Glycogen phosphorylase, brain form |
| P11233 | P11233 | Ras-related protein Ral-A |
| P11234 | P11234 | Ras-related protein Ral-B |
| P11279 | P11279 | Lysosome-associated membrane glycoprotein 1 |
| P11310 | P11310 | Medium-chain specific acyl-CoA dehydrogenase, mitochor |
| P11413 | P11413 | Glucose-6-phosphate 1-dehydrogenase |
| P11586;Q6UB35 | P11586 | C-1-tetrahydrofolate synthase, cytoplasmic;Methylenetetra |
| P11684 | P11684 | Uteroglobin |
| P11766 | P11766 | Alcohol dehydrogenase class-3 |
| P11908 | P11908 | Ribose-phosphate pyrophosphokinase 2 |
| P11940;Q4VXU2;Q9H3 | P11940 | Polyadenylate-binding protein 1 |
| P12081;P49590 | P12081 | Histidine--tRNA ligase, cytoplasmic |
| P12235 | P12235 | ADP/ATP translocase 1 |
| P12236 | P12236 | ADP/ATP translocase 3;ADP/ATP translocase 3, N-terminall |
| P12268;P20839 | P12268 | Inosine-5-monophosphate dehydrogenase 2 |
| P12277;P06732 | P12277 | Creatine kinase B-type |
| P12429 | P12429 | Annexin A3 |
| P12532;P17540 | P12532 | Creatine kinase U-type, mitochondrial |
| P12814;P35609 | P12814 | Alpha-actinin-1 |
| P12830 | P12830 | Cadherin-1;E-Cad/CTF1;E-Cad/CTF2;E-Cad/CTF3 |
| P12955 | P12955 | Xaa-Pro dipeptidase |
| P12956 | P12956 | X-ray repair cross-complementing protein 6 |
| P13010 | P13010 | X-ray repair cross-complementing protein 5 |
| P13073 | P13073 | Cytochrome c oxidase subunit 4 isoform 1, mitochondrial |
| P13164;Q01629;Q016 | P13164;Q01629;Q01 | Interferon-induced transmembrane protein 1;Interferon-in |
| P13284 | P13284 | Gamma-interferon-inducible lysosomal thiol reductase |
| P13489 | P13489 | Ribonuclease inhibitor |
| P13533;Q9H6N6 | P13533 | Myosin-6 |
| P13639 | P13639 | Elongation factor 2 |
| P13667 | P13667 | Protein disulfide-isomerase A4 |
| P13688 | P13688 | Carcinoembryonic antigen-related cell adhesion molecule |

| | | |
|---|---|---|
| P13693;Q9HAU6;Q56L | P13693 | Translationally-controlled tumor protein |
| P13716 | P13716 | Delta-aminolevulinic acid dehydratase |
| P13726 | P13726 | Tissue factor |
| P13797 | P13797 | Plastin-3 |
| P13798 | P13798 | Acylamino-acid-releasing enzyme |
| P13804 | P13804 | Electron transfer flavoprotein subunit alpha, mitochondrial |
| P13861 | P13861 | cAMP-dependent protein kinase type II-alpha regulatory su |
| Q5VT79;P13928 | Q5VT79;P13928 | Annexin A8-like protein 1;Annexin A8 |
| P13987 | P13987 | CD59 glycoprotein |
| P14174 | P14174 | Macrophage migration inhibitory factor |
| P14314 | P14314 | Glucosidase 2 subunit beta |
| P14317 | P14317 | Hematopoietic lineage cell-specific protein |
| P14324 | P14324 | Farnesyl pyrophosphate synthase |
| P14406 | P14406 | Cytochrome c oxidase subunit 7A2, mitochondrial |
| P14550 | P14550 | Aldo-keto reductase family 1 member A1 |
| P14618;P30613 | P14618 | Pyruvate kinase PKM |
| P14621 | P14621 | Acylphosphatase-2 |
| P14625;Q58FF3 | P14625 | Endoplasmin |
| P63162;P14678 | P63162;P14678 | Small nuclear ribonucleoprotein-associated protein N;Sma |
| P14735 | P14735 | Insulin-degrading enzyme |
| P14780 | P14780 | Matrix metalloproteinase-9;67 kDa matrix metalloproteinas |
| P14854 | P14854 | Cytochrome c oxidase subunit 6B1 |
| P14866 | P14866 | Heterogeneous nuclear ribonucleoprotein L |
| P14868 | P14868 | Aspartate--tRNA ligase, cytoplasmic |
| P14902 | P14902 | Indoleamine 2,3-dioxygenase 1 |
| P14923 | P14923 | Junction plakoglobin |
| P14927 | P14927 | Cytochrome b-c1 complex subunit 7 |
| P15104 | P15104 | Glutamine synthetase |
| P15121 | P15121 | Aldo-keto reductase family 1 member B1 |
| P15144 | P15144 | Aminopeptidase N |
| P15170 | P15170 | Eukaryotic peptide chain release factor GTP-binding subun |
| P15289 | P15289 | Arylsulfatase A;Arylsulfatase A component B;Arylsulfatase |
| P15291 | P15291 | Beta-1,4-galactosyltransferase 1;Processed beta-1,4-gala |
| P15311 | P15311 | Ezrin |
| P15328 | P15328 | Folate receptor alpha |
| P15374 | P15374 | Ubiquitin carboxyl-terminal hydrolase isozyme L3 |
| P15428 | P15428 | 15-hydroxyprostaglandin dehydrogenase [NAD(+)] |
| P15531 | P15531 | Nucleoside diphosphate kinase A |
| P15559 | P15559 | NAD(P)H dehydrogenase [quinone] 1 |
| P15586 | P15586 | N-acetylglucosamine-6-sulfatase |
| P15880 | P15880 | Small ribosomal subunit protein uS5 |
| P15924 | P15924 | Desmoplakin |
| P15941 | P15941 | Mucin-1;Mucin-1 subunit alpha;Mucin-1 subunit beta |
| P16050 | P16050 | Polyunsaturated fatty acid lipoxygenase ALOX15 |
| P16083 | P16083 | Ribosyldihydronicotinamide dehydrogenase [quinone] |
| P16152 | P16152 | Carbonyl reductase [NADPH] 1 |
| P16401 | P16401 | Histone H1.5 |

| | | |
|---|---|---|
| P16403 | P16403 | Histone H1.2 |
| P16435 | P16435 | NADPH--cytochrome P450 reductase |
| P16455 | P16455 | Methylated-DNA--protein-cysteine methyltransferase |
| P16591;P07332 | P16591 | Tyrosine-protein kinase Fer |
| P16615;O14983;Q930 | P16615 | Sarcoplasmic/endoplasmic reticulum calcium ATPase 2 |
| P16885 | P16885 | 1-phosphatidylinositol 4,5-bisphosphate phosphodiestera |
| P16930 | P16930 | Fumarylacetoacetase |
| P17050 | P17050 | Alpha-N-acetylgalactosaminidase |
| P17066;P48741 | P17066 | Heat shock 70 kDa protein 6 |
| P17174 | P17174 | Aspartate aminotransferase, cytoplasmic |
| P17612;P22694 | P17612;P22694 | cAMP-dependent protein kinase catalytic subunit alpha;cA |
| P17655 | P17655 | Calpain-2 catalytic subunit |
| P17844 | P17844 | Probable ATP-dependent RNA helicase DDX5 |
| P17858 | P17858 | ATP-dependent 6-phosphofructokinase, liver type |
| P17931 | P17931 | Galectin-3 |
| P17980 | P17980 | 26S proteasome regulatory subunit 6A |
| P17987 | P17987 | T-complex protein 1 subunit alpha |
| P18031 | P18031 | Tyrosine-protein phosphatase non-receptor type 1 |
| P18065 | P18065 | Insulin-like growth factor-binding protein 2 |
| P18077 | P18077 | Large ribosomal subunit protein eL33 |
| P18085 | P18085 | ADP-ribosylation factor 4 |
| P18124 | P18124 | Large ribosomal subunit protein uL30 |
| P18206 | P18206 | Vinculin |
| P18283 | P18283 | Glutathione peroxidase 2 |
| P18440 | P18440 | Arylamine N-acetyltransferase 1 |
| P18510 | P18510 | Interleukin-1 receptor antagonist protein |
| P18564 | P18564 | Integrin beta-6 |
| P18621 | P18621 | Large ribosomal subunit protein uL22 |
| P18669;Q8N0Y7;P152 | P18669 | Phosphoglycerate mutase 1 |
| P19338 | P19338 | Nucleolin |
| P19367;Q2TB90;P527 | P19367 | Hexokinase-1 |
| P19525 | P19525 | Interferon-induced, double-stranded RNA-activated protei |
| P19838 | P19838 | Nuclear factor NF-kappa-B p105 subunit;Nuclear factor NF |
| P19957 | P19957 | Elafin |
| P19971 | P19971 | Thymidine phosphorylase |
| P20073 | P20073 | Annexin A7 |
| P20290 | P20290 | Transcription factor BTF3 |
| P20339 | P20339 | Ras-related protein Rab-5A |
| P20340;Q9NRW1 | P20340;Q9NRW1 | Ras-related protein Rab-6A;Ras-related protein Rab-6B |
| P20591 | P20591 | Interferon-induced GTP-binding protein Mx1;Interferon-ind |
| P20618 | P20618 | Proteasome subunit beta type-1 |
| P20674 | P20674 | Cytochrome c oxidase subunit 5A, mitochondrial |
| P20700 | P20700 | Lamin-B1 |
| P20794 | P20794 | Serine/threonine-protein kinase MAK |
| P20810 | P20810 | Calpastatin |
| P21217 | P21217 | 3-galactosyl-N-acetylglucosaminide 4-alpha-L-fucosyltran |
| P21281 | P21281 | V-type proton ATPase subunit B, brain isoform |

| | | |
|---|---|---|
| P21291 | P21291 | Cysteine and glycine-rich protein 1 |
| P21333 | P21333 | Filamin-A |
| P21397 | P21397 | Amine oxidase [flavin-containing] A |
| P21399 | P21399 | Cytoplasmic aconitate hydratase |
| P21741 | P21741 | Midkine |
| P21796 | P21796 | Voltage-dependent anion-selective channel protein 1 |
| P21817 | P21817 | Ryanodine receptor 1 |
| P21912 | P21912 | Succinate dehydrogenase [ubiquinone] iron-sulfur subunit, |
| P21926 | P21926 | CD9 antigen |
| P21964 | P21964 | Catechol O-methyltransferase |
| P21980;P49221;O4354 | P21980 | Protein-glutamine gamma-glutamyltransferase 2 |
| P22059;Q969R2 | P22059;Q969R2 | Oxysterol-binding protein 1;Oxysterol-binding protein 2 |
| P22061 | P22061 | Protein-L-isoaspartate(D-aspartate) O-methyltransferase |
| P22087;A6NHQ2 | P22087 | rRNA 2-O-methyltransferase fibrillarin |
| P22102 | P22102 | Trifunctional purine biosynthetic protein adenosine-3;Phos |
| P22234 | P22234 | Bifunctional phosphoribosylaminoimidazole carboxylase/p |
| P22307 | P22307 | Sterol carrier protein 2 |
| P22314 | P22314 | Ubiquitin-like modifier-activating enzyme 1 |
| P22392;O60361 | P22392;O60361 | Nucleoside diphosphate kinase B;Putative nucleoside diph |
| P22528 | P22528 | Cornifin-B |
| P22532 | P22532 | Small proline-rich protein 2D |
| P22570 | P22570 | NADPH:adrenodoxin oxidoreductase, mitochondrial |
| P22612 | P22612 | cAMP-dependent protein kinase catalytic subunit gamma |
| P22626 | P22626 | Heterogeneous nuclear ribonucleoproteins A2/B1 |
| P22695 | P22695 | Cytochrome b-c1 complex subunit 2, mitochondrial |
| P22735 | P22735 | Protein-glutamine gamma-glutamyltransferase K |
| P23141;Q9UKY3 | P23141 | Liver carboxylesterase 1 |
| P23246 | P23246 | Splicing factor, proline- and glutamine-rich |
| P23284 | P23284 | Peptidyl-prolyl cis-trans isomerase B |
| P23381 | P23381 | Tryptophan--tRNA ligase, cytoplasmic;T1-TrpRS;T2-TrpRS |
| P23396 | P23396 | Small ribosomal subunit protein uS3 |
| P23526 | P23526 | Adenosylhomocysteinase |
| P23528 | P23528 | Cofilin-1 |
| P23743 | P23743 | Diacylglycerol kinase alpha |
| P23786 | P23786 | Carnitine O-palmitoyltransferase 2, mitochondrial |
| P24534 | P24534 | Elongation factor 1-beta |
| P24539 | P24539 | ATP synthase F(0) complex subunit B1, mitochondrial |
| P24666 | P24666 | Low molecular weight phosphotyrosine protein phosphatas |
| P24752 | P24752 | Acetyl-CoA acetyltransferase, mitochondrial |
| P24821 | P24821 | Tenascin |
| P25325 | P25325 | 3-mercaptopyruvate sulfurtransferase |
| P25398 | P25398 | Small ribosomal subunit protein eS12 |
| P25685 | P25685 | DnaJ homolog subfamily B member 1 |
| P25686 | P25686 | DnaJ homolog subfamily B member 2 |
| P25705 | P25705 | ATP synthase subunit alpha, mitochondrial |
| P25713 | P25713 | Metallothionein-3 |
| P25774 | P25774 | Cathepsin S |

| | | |
|---|---|---|
| P25786 | P25786 | Proteasome subunit alpha type-1 |
| P25787 | P25787 | Proteasome subunit alpha type-2 |
| P25788 | P25788 | Proteasome subunit alpha type-3 |
| P25789 | P25789 | Proteasome subunit alpha type-4 |
| P25815 | P25815 | Protein S100-P |
| P26038 | P26038 | Moesin |
| P26368 | P26368 | Splicing factor U2AF 65 kDa subunit |
| P26373 | P26373 | Large ribosomal subunit protein eL13 |
| P26440 | P26440 | Isovaleryl-CoA dehydrogenase, mitochondrial |
| P26447 | P26447 | Protein S100-A4 |
| P26572 | P26572 | Alpha-1,3-mannosyl-glycoprotein 2-beta-N-acetylglucosar |
| P26599 | P26599 | Polypyrimidine tract-binding protein 1 |
| P26639;A2RTX5;Q9BW | P26639 | Threonine--tRNA ligase 1, cytoplasmic |
| P26640 | P26640 | Valine--tRNA ligase |
| P26641 | P26641 | Elongation factor 1-gamma |
| P26885 | P26885 | Peptidyl-prolyl cis-trans isomerase FKBP2 |
| P27105 | P27105 | Stomatin |
| P27348 | P27348 | 14-3-3 protein theta |
| P27361;P31152;Q166 | P27361 | Mitogen-activated protein kinase 3 |
| P27635;Q96L21 | P27635;Q96L21 | Large ribosomal subunit protein uL16;Ribosomal protein ul |
| P27695 | P27695 | DNA repair nuclease/redox regulator APEX1;DNA repair nu |
| P27701 | P27701 | CD82 antigen |
| P27797 | P27797 | Calreticulin |
| P27816 | P27816 | Microtubule-associated protein 4 |
| P27824 | P27824 | Calnexin |
| P28062 | P28062 | Proteasome subunit beta type-8 |
| P28065 | P28065 | Proteasome subunit beta type-9 |
| P28066 | P28066 | Proteasome subunit alpha type-5 |
| P28070 | P28070 | Proteasome subunit beta type-4 |
| P28072 | P28072 | Proteasome subunit beta type-6 |
| P28074 | P28074 | Proteasome subunit beta type-5 |
| P28161;P21266 | P28161 | Glutathione S-transferase Mu 2 |
| P28288 | P28288 | ATP-binding cassette sub-family D member 3 |
| P28331 | P28331 | NADH-ubiquinone oxidoreductase 75 kDa subunit, mitoch |
| P28482 | P28482 | Mitogen-activated protein kinase 1 |
| P28838 | P28838 | Cytosol aminopeptidase |
| P29034 | P29034 | Protein S100-A2 |
| P29144 | P29144 | Tripeptidyl-peptidase 2 |
| P29218 | P29218 | Inositol monophosphatase 1 |
| P29350 | P29350 | Tyrosine-protein phosphatase non-receptor type 6 |
| P29373 | P29373 | Cellular retinoic acid-binding protein 2 |
| P29401 | P29401 | Transketolase |
| P29466;Q5EG05 | P29466 | Caspase-1;Caspase-1 subunit p20;Caspase-1 subunit p10 |
| P29508 | P29508 | Serpin B3 |
| P29590 | P29590 | Protein PML |
| P29692 | P29692 | Elongation factor 1-delta |
| P29728 | P29728 | 2-5-oligoadenylate synthase 2 |

| | | |
|---|---|---|
| P29966 | P29966 | Myristoylated alanine-rich C-kinase substrate |
| P29992 | P29992 | Guanine nucleotide-binding protein subunit alpha-11 |
| P30039 | P30039 | Phenazine biosynthesis-like domain-containing protein |
| P30040 | P30040 | Endoplasmic reticulum resident protein 29 |
| P30041 | P30041 | Peroxiredoxin-6 |
| P30043 | P30043 | Flavin reductase (NADPH) |
| P30044 | P30044 | Peroxiredoxin-5, mitochondrial |
| P30046;A6NHG4 | P30046;A6NHG4 | D-dopachrome decarboxylase;D-dopachrome decarboxyla |
| P30047 | P30047 | GTP cyclohydrolase 1 feedback regulatory protein |
| P30048 | P30048 | Thioredoxin-dependent peroxide reductase, mitochondrial |
| P30050 | P30050 | Large ribosomal subunit protein uL11 |
| P30084 | P30084 | Enoyl-CoA hydratase, mitochondrial |
| P30085 | P30085 | UMP-CMP kinase |
| P30086 | P30086 | Phosphatidylethanolamine-binding protein 1;Hippocampal |
| P30101 | P30101 | Protein disulfide-isomerase A3 |
| P30153;P30154 | P30153 | Serine/threonine-protein phosphatase 2A 65 kDa regulator |
| P30520 | P30520 | Adenylosuccinate synthetase isozyme 2 |
| P30536 | P30536 | Translocator protein |
| P30566 | P30566 | Adenylosuccinate lyase |
| P30626 | P30626 | Sorcin |
| P30711;P0CG30;P0CG | P30711 | Glutathione S-transferase theta-1 |
| P30740;O75830 | P30740 | Leukocyte elastase inhibitor |
| P30838 | P30838 | Aldehyde dehydrogenase, dimeric NADP-preferring |
| P31040 | P31040 | Succinate dehydrogenase [ubiquinone] flavoprotein subun |
| P31150 | P31150 | Rab GDP dissociation inhibitor alpha |
| P31151;Q86SG5 | P31151 | Protein S100-A7 |
| P31153;Q00266 | P31153 | S-adenosylmethionine synthase isoform type-2 |
| P31323 | P31323 | cAMP-dependent protein kinase type II-beta regulatory sub |
| P31513 | P31513 | Flavin-containing monooxygenase 3 |
| P31689 | P31689 | DnaJ homolog subfamily A member 1 |
| P31749 | P31749 | RAC-alpha serine/threonine-protein kinase |
| P31930;O75439 | P31930 | Cytochrome b-c1 complex subunit 1, mitochondrial |
| P31939 | P31939 | Bifunctional purine biosynthesis protein ATIC;Phosphoribo: |
| P31941;Q9UH17 | P31941;Q9UH17 | DNA dC->dU-editing enzyme APOBEC-3A;DNA dC->dU-edi |
| P31942 | P31942 | Heterogeneous nuclear ribonucleoprotein H3 |
| P31943 | P31943 | Heterogeneous nuclear ribonucleoprotein H;Heterogeneou |
| P31946 | P31946 | 14-3-3 protein beta/alpha;14-3-3 protein beta/alpha, N-ter |
| P31947 | P31947 | 14-3-3 protein sigma |
| P31948 | P31948 | Stress-induced-phosphoprotein 1 |
| P31949 | P31949 | Protein S100-A11;Protein S100-A11, N-terminally processe |
| P32119 | P32119 | Peroxiredoxin-2 |
| P32189;Q14409 | P32189;Q14409 | Glycerol kinase;Glycerol kinase 3 |
| P32455;P32456;Q9H0I | P32455 | Guanylate-binding protein 1 |
| P32856 | P32856 | Syntaxin-2 |
| P32969 | P32969 | Large ribosomal subunit protein uL6 |
| P33121 | P33121 | Long-chain-fatty-acid--CoA ligase 1 |
| P33176;Q12840;O602 | P33176 | Kinesin-1 heavy chain |

| | | |
|---|---|---|
| P34897 | P34897 | Serine hydroxymethyltransferase, mitochondrial |
| P34913 | P34913 | Bifunctional epoxide hydrolase 2;Cytosolic epoxide hydrola |
| P34932 | P34932 | Heat shock 70 kDa protein 4 |
| P34949 | P34949 | Mannose-6-phosphate isomerase |
| P35080 | P35080 | Profilin-2 |
| P35221 | P35221 | Catenin alpha-1 |
| P35222 | P35222 | Catenin beta-1 |
| P35228;P29474 | P35228 | Nitric oxide synthase, inducible |
| P35232 | P35232 | Prohibitin 1 |
| P35237 | P35237 | Serpin B6 |
| P35241 | P35241 | Radixin |
| P35244 | P35244 | Replication protein A 14 kDa subunit |
| P35268 | P35268 | Large ribosomal subunit protein eL22 |
| P35270;REV__Q16649 | P35270 | Sepiapterin reductase |
| P35321 | P35321 | Cornifin-A |
| P35326 | P35326 | Small proline-rich protein 2A |
| P35579;REV__Q9UKV3 | P35579 | Myosin-9 |
| P35606 | P35606 | Coatomer subunit beta |
| P35626;P25098 | P35626 | G protein-coupled receptor kinase 3 |
| P35749;A7E2Y1 | P35749 | Myosin-11 |
| P35754 | P35754 | Glutaredoxin-1 |
| P35813 | P35813 | Protein phosphatase 1A |
| P35914 | P35914 | Hydroxymethylglutaryl-CoA lyase, mitochondrial |
| P35998 | P35998 | 26S proteasome regulatory subunit 7 |
| P36405 | P36405 | ADP-ribosylation factor-like protein 3 |
| P36507 | P36507 | Dual specificity mitogen-activated protein kinase kinase 2 |
| P36542 | P36542 | ATP synthase subunit gamma, mitochondrial |
| P36543;Q96A05 | P36543 | V-type proton ATPase subunit E 1 |
| P36578 | P36578 | Large ribosomal subunit protein uL4 |
| P36776 | P36776 | Lon protease homolog, mitochondrial |
| P36871 | P36871 | Phosphoglucomutase-1 |
| P36873 | P36873 | Serine/threonine-protein phosphatase PP1-gamma catalyt |
| P36952 | P36952 | Serpin B5 |
| P36955 | P36955 | Pigment epithelium-derived factor |
| P36957 | P36957 | Dihydrolipoyllysine-residue succinyltransferase componen |
| P36969 | P36969 | Phospholipid hydroperoxide glutathione peroxidase GPX4 |
| P37108 | P37108 | Signal recognition particle 14 kDa protein |
| P37802 | P37802 | Transgelin-2 |
| P37837 | P37837 | Transaldolase |
| P37840 | P37840 | Alpha-synuclein |
| P38117 | P38117 | Electron transfer flavoprotein subunit beta |
| P38159;Q96E39;O755: | P38159 | RNA-binding motif protein, X chromosome;RNA-binding mc |
| P38606 | P38606 | V-type proton ATPase catalytic subunit A |
| P38646 | P38646 | Stress-70 protein, mitochondrial |
| P38919 | P38919 | Eukaryotic initiation factor 4A-III;Eukaryotic initiation factor |
| P39023;Q92901 | P39023 | Large ribosomal subunit protein uL3 |
| P39656 | P39656 | Dolichyl-diphosphooligosaccharide--protein glycosyltransf |

| | | |
|---|---|---|
| P39687;O43423 | P39687 | Acidic leucine-rich nuclear phosphoprotein 32 family mem |
| P40121 | P40121 | Macrophage-capping protein |
| P40227;Q92526 | P40227 | T-complex protein 1 subunit zeta |
| P40306 | P40306 | Proteasome subunit beta type-10 |
| P40394 | P40394 | All-trans-retinol dehydrogenase [NAD(+)] ADH7 |
| P40429;Q6NVV1 | P40429 | Large ribosomal subunit protein uL13 |
| P40616 | P40616 | ADP-ribosylation factor-like protein 1 |
| P40763 | P40763 | Signal transducer and activator of transcription 3 |
| P40925 | P40925 | Malate dehydrogenase, cytoplasmic |
| P40926 | P40926 | Malate dehydrogenase, mitochondrial |
| P40939 | P40939 | Trifunctional enzyme subunit alpha, mitochondrial;Long-ch |
| P41091;Q2VIR3 | P41091;Q2VIR3 | Eukaryotic translation initiation factor 2 subunit 3;Eukaryot |
| P41208;Q12798 | P41208 | Centrin-2 |
| P41226 | P41226 | Ubiquitin-like modifier-activating enzyme 7 |
| P41236;Q6NXS1 | P41236;Q6NXS1 | Protein phosphatase inhibitor 2;Protein phosphatase inhibi |
| P41240 | P41240 | Tyrosine-protein kinase CSK |
| P41250 | P41250 | Glycine--tRNA ligase |
| P41252 | P41252 | Isoleucine--tRNA ligase, cytoplasmic |
| P41567;O60739 | P41567 | Eukaryotic translation initiation factor 1 |
| P42224 | P42224 | Signal transducer and activator of transcription 1-alpha/be |
| P42330 | P42330 | Aldo-keto reductase family 1 member C3 |
| P42574 | P42574 | Caspase-3;Caspase-3 subunit p17;Caspase-3 subunit p12 |
| P42677;Q71UM5 | P42677;Q71UM5 | Small ribosomal subunit protein eS27;Ribosomal protein e |
| P42685 | P42685 | Tyrosine-protein kinase FRK |
| P42704 | P42704 | Leucine-rich PPR motif-containing protein, mitochondrial |
| P42766 | P42766 | Large ribosomal subunit protein uL29 |
| P42771;P42772 | P42771;P42772 | Cyclin-dependent kinase inhibitor 2A;Cyclin-dependent kir |
| P42830 | P42830 | C-X-C motif chemokine 5;ENA-78(8-78);ENA-78(9-78) |
| P43034 | P43034 | Platelet-activating factor acetylhydrolase IB subunit beta |
| P43243 | P43243 | Matrin-3 |
| P43304 | P43304 | Glycerol-3-phosphate dehydrogenase, mitochondrial |
| P43307 | P43307 | Translocon-associated protein subunit alpha |
| P43353;P48448 | P43353 | Aldehyde dehydrogenase family 3 member B1 |
| P43359 | P43359 | Putative melanoma-associated antigen 5P |
| P43487 | P43487 | Ran-specific GTPase-activating protein |
| P43490 | P43490 | Nicotinamide phosphoribosyltransferase |
| P43686 | P43686 | 26S proteasome regulatory subunit 6B |
| P45877 | P45877 | Peptidyl-prolyl cis-trans isomerase C |
| P45880 | P45880 | Voltage-dependent anion-selective channel protein 2 |
| P45974;Q92995 | P45974 | Ubiquitin carboxyl-terminal hydrolase 5 |
| P45985 | P45985 | Dual specificity mitogen-activated protein kinase kinase 4 |
| P46060 | P46060 | Ran GTPase-activating protein 1 |
| P46108 | P46108 | Adapter molecule crk |
| P46109 | P46109 | Crk-like protein |
| P46459 | P46459 | Vesicle-fusing ATPase |
| P46776 | P46776 | Large ribosomal subunit protein uL15 |
| P46777 | P46777 | Large ribosomal subunit protein uL18 |

| | | |
|---|---|---|
| P46778 | P46778 | Large ribosomal subunit protein eL21 |
| P46779 | P46779 | Large ribosomal subunit protein eL28 |
| P46781 | P46781 | Small ribosomal subunit protein uS4 |
| P46782 | P46782 | Small ribosomal subunit protein uS7;Small ribosomal subu |
| P46783 | P46783 | Small ribosomal subunit protein eS10 |
| P46940;Q86VI3 | P46940 | Ras GTPase-activating-like protein IQGAP1 |
| P46977;Q8TCJ2 | P46977 | Dolichyl-diphosphooligosaccharide--protein glycosyltransf |
| P47224 | P47224 | Guanine nucleotide exchange factor MSS4 |
| P47755 | P47755 | F-actin-capping protein subunit alpha-2 |
| P47756 | P47756 | F-actin-capping protein subunit beta |
| P47895;P30837 | P47895 | Retinaldehyde dehydrogenase 3 |
| P47897 | P47897 | Glutamine--tRNA ligase |
| P47929 | P47929 | Galectin-7 |
| P47985;P0C7P4 | P47985;P0C7P4 | Cytochrome b-c1 complex subunit Rieske, mitochondrial;C |
| P48047 | P48047 | ATP synthase subunit O, mitochondrial |
| P48147 | P48147 | Prolyl endopeptidase |
| P48163 | P48163 | NADP-dependent malic enzyme |
| P48444 | P48444 | Coatomer subunit delta |
| P48506 | P48506 | Glutamate--cysteine ligase catalytic subunit |
| P48507 | P48507 | Glutamate--cysteine ligase regulatory subunit |
| P48594 | P48594 | Serpin B4 |
| P48595 | P48595 | Serpin B10 |
| P48637 | P48637 | Glutathione synthetase |
| P48643 | P48643 | T-complex protein 1 subunit epsilon |
| P48729;Q8N752 | P48729;Q8N752 | Casein kinase I isoform alpha;Casein kinase I isoform alpha |
| P48735 | P48735 | Isocitrate dehydrogenase [NADP], mitochondrial |
| P48739 | P48739 | Phosphatidylinositol transfer protein beta isoform |
| P49006 | P49006 | MARCKS-related protein |
| P49184 | P49184 | Deoxyribonuclease-1-like 1 |
| P49189 | P49189 | 4-trimethylaminobutyraldehyde dehydrogenase;4-trimethy |
| P49247 | P49247 | Ribose-5-phosphate isomerase |
| P49257 | P49257 | Protein ERGIC-53 |
| P49281 | P49281 | Natural resistance-associated macrophage protein 2 |
| P49327 | P49327 | Fatty acid synthase;[Acyl-carrier-protein] S-acetyltransfera |
| P49354 | P49354 | Protein farnesyltransferase/geranylgeranyltransferase type |
| P49368 | P49368 | T-complex protein 1 subunit gamma |
| P49407;P36575 | P49407 | Beta-arrestin-1 |
| P49411 | P49411 | Elongation factor Tu, mitochondrial |
| P49419 | P49419 | Alpha-aminoadipic semialdehyde dehydrogenase |
| P49441 | P49441 | Inositol polyphosphate 1-phosphatase |
| P49458 | P49458 | Signal recognition particle 9 kDa protein |
| P49588 | P49588 | Alanine--tRNA ligase, cytoplasmic |
| P49589 | P49589 | Cysteine--tRNA ligase, cytoplasmic |
| P49591 | P49591 | Serine--tRNA ligase, cytoplasmic |
| P49662;P51878 | P49662;P51878 | Caspase-4;Caspase-4 subunit p10;Caspase-4 subunit p20 |
| P49720 | P49720 | Proteasome subunit beta type-3 |
| P49721 | P49721 | Proteasome subunit beta type-2 |

| | | |
|---|---|---|
| P49748 | P49748 | Very long-chain specific acyl-CoA dehydrogenase, mitocho |
| P49755 | P49755 | Transmembrane emp24 domain-containing protein 10 |
| P49773 | P49773 | Adenosine 5-monophosphoramidase HINT1 |
| P49788 | P49788 | Retinoic acid receptor responder protein 1 |
| P49789 | P49789 | Bis(5-adenosyl)-triphosphatase |
| P49841;P49840 | P49841;P49840 | Glycogen synthase kinase-3 beta;Glycogen synthase kinas |
| P49902 | P49902 | Cytosolic purine 5-nucleotidase |
| P49914 | P49914 | 5-formyltetrahydrofolate cyclo-ligase |
| P50135 | P50135 | Histamine N-methyltransferase |
| P50148 | P50148 | Guanine nucleotide-binding protein G(q) subunit alpha |
| P50213 | P50213 | Isocitrate dehydrogenase [NAD] subunit alpha, mitochondr |
| P50225;P50226 | P50225;P50226 | Sulfotransferase 1A1;Sulfotransferase 1A2 |
| P50238 | P50238 | Cysteine-rich protein 1 |
| P50336 | P50336 | Protoporphyrinogen oxidase |
| P50395 | P50395 | Rab GDP dissociation inhibitor beta |
| P50452;P50453 | P50452;P50453 | Serpin B8;Serpin B9 |
| P50454 | P50454 | Serpin H1 |
| P50479 | P50479 | PDZ and LIM domain protein 4 |
| P50502;Q8IZP2 | P50502 | Hsc70-interacting protein |
| P50570;Q05193;Q9UQ | P50570 | Dynamin-2 |
| P50583 | P50583 | Bis(5-nucleosyl)-tetraphosphatase [asymmetrical] |
| P50851 | P50851 | Lipopolysaccharide-responsive and beige-like anchor prote |
| P50914 | P50914 | Large ribosomal subunit protein eL14 |
| P50990 | P50990 | T-complex protein 1 subunit theta |
| P50991 | P50991 | T-complex protein 1 subunit delta |
| P50995;P27216 | P50995 | Annexin A11 |
| P51148 | P51148 | Ras-related protein Rab-5C |
| P51149 | P51149 | Ras-related protein Rab-7a |
| P51153;P59190;P2033 | P51153 | Ras-related protein Rab-13 |
| P51452 | P51452 | Dual specificity protein phosphatase 3 |
| P51553 | P51553 | Isocitrate dehydrogenase [NAD] subunit gamma, mitochon |
| P51570 | P51570 | Galactokinase |
| P51571 | P51571 | Translocon-associated protein subunit delta |
| P51572 | P51572 | B-cell receptor-associated protein 31 |
| P51580 | P51580 | Thiopurine S-methyltransferase |
| P51648 | P51648 | Aldehyde dehydrogenase family 3 member A2 |
| P51654;REV__Q96HE7 | P51654 | Glypican-3;Glypican-3 alpha subunit;Glypican-3 beta subu |
| P51659 | P51659 | Peroxisomal multifunctional enzyme type 2;(3R)-hydroxyac |
| P51665 | P51665 | 26S proteasome non-ATPase regulatory subunit 7 |
| P51688 | P51688 | N-sulphoglucosamine sulphohydrolase |
| P51812;Q15349;Q9UK | P51812 | Ribosomal protein S6 kinase alpha-3 |
| P51857 | P51857 | Aldo-keto reductase family 1 member D1 |
| P51858 | P51858 | Hepatoma-derived growth factor |
| P51991 | P51991 | Heterogeneous nuclear ribonucleoprotein A3 |
| P51993;Q11128 | P51993;Q11128 | 4-galactosyl-N-acetylglucosaminide 3-alpha-L-fucosyltran |
| P52209 | P52209 | 6-phosphogluconate dehydrogenase, decarboxylating |
| P52272 | P52272 | Heterogeneous nuclear ribonucleoprotein M |

| | | |
|---|---|---|
| P52294;O60684;O151 | P52294;O60684;O15 | Importin subunit alpha-5;Importin subunit alpha-5, N-term |
| P52306 | P52306 | Rap1 GTPase-GDP dissociation stimulator 1 |
| P52565 | P52565 | Rho GDP-dissociation inhibitor 1 |
| P52566 | P52566 | Rho GDP-dissociation inhibitor 2 |
| P52597 | P52597 | Heterogeneous nuclear ribonucleoprotein F;Heterogeneou |
| P52788 | P52788 | Spermine synthase |
| P52789 | P52789 | Hexokinase-2 |
| P52888 | P52888 | Thimet oligopeptidase |
| P52895 | P52895 | Aldo-keto reductase family 1 member C2 |
| P52907 | P52907 | F-actin-capping protein subunit alpha-1 |
| P52943 | P52943 | Cysteine-rich protein 2 |
| P53004 | P53004 | Biliverdin reductase A |
| P53007 | P53007 | Tricarboxylate transport protein, mitochondrial |
| P53041 | P53041 | Serine/threonine-protein phosphatase 5 |
| P53367 | P53367 | Arfaptin-1 |
| P53384 | P53384 | Cytosolic Fe-S cluster assembly factor NUBP1 |
| P53396 | P53396 | ATP-citrate synthase |
| P53597 | P53597 | Succinate--CoA ligase [ADP/GDP-forming] subunit alpha, n |
| P53602 | P53602 | Diphosphomevalonate decarboxylase |
| P53609 | P53609 | Geranylgeranyl transferase type-1 subunit beta |
| P53611 | P53611 | Geranylgeranyl transferase type-2 subunit beta |
| P53618 | P53618 | Coatomer subunit beta |
| P53621 | P53621 | Coatomer subunit alpha;Xenin;Proxenin |
| P53634 | P53634 | Dipeptidyl peptidase 1;Dipeptidyl peptidase 1 exclusion do |
| P53680 | P53680 | AP-2 complex subunit sigma |
| P53990 | P53990 | IST1 homolog |
| P53999 | P53999 | Activated RNA polymerase II transcriptional coactivator p1 |
| P54105 | P54105 | Methylosome subunit pICln |
| P54136 | P54136 | Arginine--tRNA ligase, cytoplasmic |
| P54577 | P54577 | Tyrosine--tRNA ligase, cytoplasmic;Tyrosine--tRNA ligase, |
| P54578 | P54578 | Ubiquitin carboxyl-terminal hydrolase 14 |
| P54652 | P54652 | Heat shock-related 70 kDa protein 2 |
| P54727 | P54727 | UV excision repair protein RAD23 homolog B |
| P54819 | P54819 | Adenylate kinase 2, mitochondrial;Adenylate kinase 2, mit |
| P54886 | P54886 | Delta-1-pyrroline-5-carboxylate synthase;Glutamate 5-kin |
| P54920;Q9H115 | P54920 | Alpha-soluble NSF attachment protein |
| P55036 | P55036 | 26S proteasome non-ATPase regulatory subunit 4 |
| P55042;P55040;O756 | P55042 | GTP-binding protein RAD |
| P55058 | P55058 | Phospholipid transfer protein |
| P55060 | P55060 | Exportin-2 |
| P55064 | P55064 | Aquaporin-5 |
| P55072 | P55072 | Transitional endoplasmic reticulum ATPase |
| P55084 | P55084 | Trifunctional enzyme subunit beta, mitochondrial;3-ketoac |
| P55145 | P55145 | Mesencephalic astrocyte-derived neurotrophic factor |
| P55209 | P55209 | Nucleosome assembly protein 1-like 1 |
| P55210 | P55210 | Caspase-7;Caspase-7 subunit p20;Caspase-7 subunit p11 |
| P55263 | P55263 | Adenosine kinase |

| | | |
|---|---|---|
| P55327 | P55327 | Tumor protein D52 |
| P55735 | P55735 | Protein SEC13 homolog |
| P55769 | P55769 | NHP2-like protein 1;NHP2-like protein 1, N-terminally proc |
| P55786;A6NEC2 | P55786 | Puromycin-sensitive aminopeptidase |
| P55795 | P55795 | Heterogeneous nuclear ribonucleoprotein H2;Heterogene |
| P55884 | P55884 | Eukaryotic translation initiation factor 3 subunit B |
| P55957 | P55957 | BH3-interacting domain death agonist;BH3-interacting dor |
| P56134 | P56134 | ATP synthase subunit f, mitochondrial |
| P56192 | P56192 | Methionine--tRNA ligase, cytoplasmic |
| P56385 | P56385 | ATP synthase subunit e, mitochondrial;ATP synthase subur |
| P56537 | P56537 | Eukaryotic translation initiation factor 6 |
| P56597 | P56597 | Nucleoside diphosphate kinase homolog 5 |
| P56715 | P56715 | Oxygen-regulated protein 1 |
| P57076 | P57076 | Cilia- and flagella-associated protein 298 |
| P57088 | P57088 | Transmembrane protein 33 |
| P57721 | P57721 | Poly(rC)-binding protein 3 |
| P57735 | P57735 | Ras-related protein Rab-25 |
| P57737 | P57737 | Coronin-7 |
| P57764 | P57764 | Gasdermin-D;Gasdermin-D, N-terminal;Gasdermin-D, C-te |
| P58107 | P58107 | Epiplakin |
| P58340 | P58340 | Myeloid leukemia factor 1 |
| P58505 | P58505 | Uncharacterized protein C21orf58 |
| P58546 | P58546 | Myotrophin |
| P59910 | P59910 | DnaJ homolog subfamily B member 13 |
| P59998 | P59998 | Actin-related protein 2/3 complex subunit 4 |
| P60174 | P60174 | Triosephosphate isomerase |
| P60660;P14649 | P60660 | Myosin light polypeptide 6 |
| P60709;Q9BYX7 | P60709 | Actin, cytoplasmic 1;Actin, cytoplasmic 1, N-terminally pro |
| P60842 | P60842 | Eukaryotic initiation factor 4A-I |
| P60866 | P60866 | Small ribosomal subunit protein uS10 |
| P60891 | P60891 | Ribose-phosphate pyrophosphokinase 1 |
| P60900 | P60900 | Proteasome subunit alpha type-6 |
| P60903 | P60903 | Protein S100-A10 |
| P60953 | P60953 | Cell division control protein 42 homolog |
| P60981 | P60981 | Destrin |
| P61006 | P61006 | Ras-related protein Rab-8A |
| P61019;Q8WUD1 | P61019;Q8WUD1 | Ras-related protein Rab-2A;Ras-related protein Rab-2B |
| P61020 | P61020 | Ras-related protein Rab-5B |
| P61026 | P61026 | Ras-related protein Rab-10 |
| P61077;P62837;Q9Y2X | P61077;P62837 | Ubiquitin-conjugating enzyme E2 D3;Ubiquitin-conjugating |
| P61081 | P61081 | NEDD8-conjugating enzyme Ubc12 |
| P61086 | P61086 | Ubiquitin-conjugating enzyme E2 K |
| P61088;Q5JXB2 | P61088 | Ubiquitin-conjugating enzyme E2 N |
| P61106 | P61106 | Ras-related protein Rab-14 |
| P61158;Q9P1U1;Q9C0 | P61158 | Actin-related protein 3 |
| P61160 | P61160 | Actin-related protein 2 |
| P61163 | P61163 | Alpha-centractin |

| | | |
|---|---|---|
| P61204 | P61204 | ADP-ribosylation factor 3 |
| P61221 | P61221 | ATP-binding cassette sub-family E member 1 |
| P61224;A6NIZ1 | P61224;A6NIZ1 | Ras-related protein Rap-1b;Ras-related protein Rap-1b-like |
| P61247 | P61247 | Small ribosomal subunit protein eS1 |
| P61254;Q9UNX3 | P61254;Q9UNX3 | Large ribosomal subunit protein uL24;Ribosomal protein ul |
| P61313 | P61313 | Large ribosomal subunit protein eL15 |
| P61326;Q96A72 | P61326;Q96A72 | Protein mago nashi homolog;Protein mago nashi homolog |
| P61353 | P61353 | Large ribosomal subunit protein eL27 |
| P61421 | P61421 | V-type proton ATPase subunit d 1 |
| P61457 | P61457 | Pterin-4-alpha-carbinolamine dehydratase |
| P61513;A6NKH3 | P61513 | Large ribosomal subunit protein eL43 |
| P61586 | P61586 | Transforming protein RhoA |
| P61604 | P61604 | 10 kDa heat shock protein, mitochondrial |
| P61619;Q9H9S3 | P61619 | Protein transport protein Sec61 subunit alpha isoform 1 |
| P61626 | P61626 | Lysozyme C |
| P61764 | P61764 | Syntaxin-binding protein 1 |
| P61769 | P61769 | Beta-2-microglobulin;Beta-2-microglobulin form pI 5.3 |
| P61803 | P61803 | Dolichyl-diphosphooligosaccharide--protein glycosyltransf |
| P61916 | P61916 | NPC intracellular cholesterol transporter 2 |
| P61956;P55854 | P61956;P55854 | Small ubiquitin-related modifier 2;Small ubiquitin-related r |
| P61960 | P61960 | Ubiquitin-fold modifier 1 |
| P61966;P56377 | P61966 | AP-1 complex subunit sigma-1A |
| P61970 | P61970 | Nuclear transport factor 2 |
| P61978 | P61978 | Heterogeneous nuclear ribonucleoprotein K |
| P61981 | P61981 | 14-3-3 protein gamma;14-3-3 protein gamma, N-terminally |
| P62070 | P62070 | Ras-related protein R-Ras2 |
| P62081 | P62081 | Small ribosomal subunit protein eS7 |
| P62136 | P62136 | Serine/threonine-protein phosphatase PP1-alpha catalytic |
| P62140 | P62140 | Serine/threonine-protein phosphatase PP1-beta catalytic s |
| P62191 | P62191 | 26S proteasome regulatory subunit 4 |
| P62195;Q8NB90 | P62195 | 26S proteasome regulatory subunit 8 |
| P62241 | P62241 | Small ribosomal subunit protein eS8 |
| P62244 | P62244 | Small ribosomal subunit protein uS8 |
| P62249 | P62249 | Small ribosomal subunit protein uS9 |
| P62253 | P62253 | Ubiquitin-conjugating enzyme E2 G1;Ubiquitin-conjugating |
| P62256 | P62256 | Ubiquitin-conjugating enzyme E2 H |
| P62258 | P62258 | 14-3-3 protein epsilon |
| P62263 | P62263 | Small ribosomal subunit protein uS11 |
| P62266 | P62266 | Small ribosomal subunit protein uS12 |
| P62269 | P62269 | Small ribosomal subunit protein uS13 |
| P62273 | P62273 | Small ribosomal subunit protein uS14 |
| P62277 | P62277 | Small ribosomal subunit protein uS15 |
| P62280 | P62280 | Small ribosomal subunit protein uS17 |
| P62304 | P62304 | Small nuclear ribonucleoprotein E |
| P62316 | P62316 | Small nuclear ribonucleoprotein Sm D2 |
| P62318 | P62318 | Small nuclear ribonucleoprotein Sm D3 |
| P62328 | P62328 | Thymosin beta-4;Hemoregulatory peptide AcSDKP |

| | | |
|---|---|---|
| P62330 | P62330 | ADP-ribosylation factor 6 |
| P62333 | P62333 | 26S proteasome regulatory subunit 10B |
| P62424 | P62424 | Large ribosomal subunit protein eL8 |
| P62495 | P62495 | Eukaryotic peptide chain release factor subunit 1 |
| P62633 | P62633 | CCHC-type zinc finger nucleic acid binding protein |
| P62701;Q8TD47;P2209 | P62701 | Small ribosomal subunit protein eS4, X isoform |
| P62714 | P62714 | Serine/threonine-protein phosphatase 2A catalytic subunit |
| P62750 | P62750 | Large ribosomal subunit protein uL23 |
| P62753 | P62753 | Small ribosomal subunit protein eS6 |
| P62805 | P62805 | Histone H4 |
| P62820 | P62820 | Ras-related protein Rab-1A |
| P62826 | P62826 | GTP-binding nuclear protein Ran |
| P62829 | P62829 | Large ribosomal subunit protein uL14 |
| P62834 | P62834 | Ras-related protein Rap-1A |
| P62847 | P62847 | Small ribosomal subunit protein eS24 |
| P62851 | P62851 | Small ribosomal subunit protein eS25 |
| P62854;Q5JNZ5 | P62854;Q5JNZ5 | Small ribosomal subunit protein eS26;Putative ribosomal p |
| P62857 | P62857 | Small ribosomal subunit protein eS28 |
| P62873;P16520 | P62873 | Guanine nucleotide-binding protein G(I)/G(S)/G(T) subunit |
| P62879 | P62879 | Guanine nucleotide-binding protein G(I)/G(S)/G(T) subunit |
| P62888 | P62888 | Large ribosomal subunit protein eL30 |
| P62899 | P62899 | Large ribosomal subunit protein eL31 |
| P62906 | P62906 | Large ribosomal subunit protein uL1 |
| P62910 | P62910 | Large ribosomal subunit protein eL32 |
| P62913 | P62913 | Large ribosomal subunit protein uL5 |
| P62917 | P62917 | Large ribosomal subunit protein uL2 |
| P62937;A0A0B4J2A2;A | P62937 | Peptidyl-prolyl cis-trans isomerase A;Peptidyl-prolyl cis-tra |
| P62942;Q5VVH2 | P62942 | Peptidyl-prolyl cis-trans isomerase FKBP1A |
| P62979;P62987;P0CG | P62979;P62987;P0C | Ubiquitin-ribosomal protein eS31 fusion protein;Ubiquitin; |
| P63000;P60763 | P63000;P60763 | Ras-related C3 botulinum toxin substrate 1;Ras-related C3 |
| P63010 | P63010 | AP-2 complex subunit beta |
| Q5JWF2;P63092;P384 | Q5JWF2;P63092 | Guanine nucleotide-binding protein G(s) subunit alpha isof |
| P63096 | P63096 | Guanine nucleotide-binding protein G(i) subunit alpha-1 |
| P63104 | P63104 | 14-3-3 protein zeta/delta |
| P63151;Q00005;Q9Y2 | P63151 | Serine/threonine-protein phosphatase 2A 55 kDa regulator |
| P63167 | P63167 | Dynein light chain 1, cytoplasmic |
| P63172 | P63172 | Dynein light chain Tctex-type 1 |
| P63173 | P63173 | Large ribosomal subunit protein eL38 |
| P63208 | P63208 | S-phase kinase-associated protein 1 |
| P63218;A0A804HLA8 | P63218 | Guanine nucleotide-binding protein G(I)/G(S)/G(O) subuni |
| P63220 | P63220 | Small ribosomal subunit protein eS21 |
| P63241;Q6IS14;Q9GZ | P63241;Q6IS14;Q9G | Eukaryotic translation initiation factor 5A-1;Eukaryotic tran |
| P63244 | P63244 | Small ribosomal subunit protein RACK1;Small ribosomal su |
| P63261 | P63261 | Actin, cytoplasmic 2;Actin, cytoplasmic 2, N-terminally pro |
| P63279 | P63279 | SUMO-conjugating enzyme UBC9 |
| P67775 | P67775 | Serine/threonine-protein phosphatase 2A catalytic subunit |
| P67809;Q9Y2T7;P1698 | P67809 | Y-box-binding protein 1 |

| | | |
|---|---|---|
| P67812 | P67812 | Signal peptidase complex catalytic subunit SEC11A |
| P67870 | P67870 | Casein kinase II subunit beta |
| P67936;Q2TAC2 | P67936 | Tropomyosin alpha-4 chain |
| P68036;A0A1B0GUS4 | P68036;A0A1B0GUS4 | Ubiquitin-conjugating enzyme E2 L3;Ubiquitin-conjugating |
| P68104;Q5VTE0;Q0563 | P68104;Q5VTE0 | Elongation factor 1-alpha 1;Putative elongation factor 1-alp |
| P68133 | P68133 | Actin, alpha skeletal muscle;Actin, alpha skeletal muscle, i |
| P68363;Q9H853 | P68363 | Tubulin alpha-1B chain;Detyrosinated tubulin alpha-1B cha |
| P68366 | P68366 | Tubulin alpha-4A chain |
| P68371 | P68371 | Tubulin beta-4B chain |
| P68400;Q8NEV1 | P68400;Q8NEV1 | Casein kinase II subunit alpha;Casein kinase II subunit alph |
| P68402 | P68402 | Platelet-activating factor acetylhydrolase IB subunit alpha2 |
| P68431;Q16695;Q6NX | P68431;Q16695;Q6N | Histone H3.1;Histone H3.1t;Histone H3.3C |
| P69849;Q15155;Q5JPE | P69849;Q15155;Q5J | BOS complex subunit NOMO3;BOS complex subunit NOM( |
| P78330 | P78330 | Phosphoserine phosphatase |
| P78371 | P78371 | T-complex protein 1 subunit beta |
| P78417 | P78417 | Glutathione S-transferase omega-1 |
| P78527 | P78527 | DNA-dependent protein kinase catalytic subunit |
| P78559 | P78559 | Microtubule-associated protein 1A;MAP1A heavy chain;MA |
| P80162 | P80162 | C-X-C motif chemokine 6;Small-inducible cytokine B6, N-p |
| P80188 | P80188 | Neutrophil gelatinase-associated lipocalin |
| P80303 | P80303 | Nucleobindin-2;Nesfatin-1 |
| P80723 | P80723 | Brain acid soluble protein 1 |
| P81605 | P81605 | Dermcidin;Survival-promoting peptide;DCD-1 |
| P83731 | P83731 | Large ribosomal subunit protein eL24 |
| Q969Q0;P83881 | Q969Q0;P83881 | Ribosomal protein eL42-like;Large ribosomal subunit prote |
| P84077 | P84077 | ADP-ribosylation factor 1 |
| P84085 | P84085 | ADP-ribosylation factor 5 |
| P84090 | P84090 | Enhancer of rudimentary homolog |
| P84098 | P84098 | Large ribosomal subunit protein eL19 |
| P84103 | P84103 | Serine/arginine-rich splicing factor 3 |
| P84243 | P84243 | Histone H3.3 |
| P98088;Q02817 | P98088 | Mucin-5AC |
| P98160 | P98160 | Basement membrane-specific heparan sulfate proteoglyca |
| P99999;CON__P62894 | P99999 | Cytochrome c |
| Q00169 | Q00169 | Phosphatidylinositol transfer protein alpha isoform |
| Q00325 | Q00325 | Solute carrier family 25 member 3 |
| Q00341 | Q00341 | Vigilin |
| Q00532;Q92772;Q8IW | Q00532 | Cyclin-dependent kinase-like 1 |
| Q00535 | Q00535 | Cyclin-dependent kinase 5 |
| Q00610 | Q00610 | Clathrin heavy chain 1 |
| Q00688 | Q00688 | Peptidyl-prolyl cis-trans isomerase FKBP3 |
| Q00765 | Q00765 | Receptor expression-enhancing protein 5 |
| Q00796 | Q00796 | Sorbitol dehydrogenase |
| Q00839 | Q00839 | Heterogeneous nuclear ribonucleoprotein U |
| Q01081;Q8WU68 | Q01081;Q8WU68 | Splicing factor U2AF 35 kDa subunit;Splicing factor U2AF 2 |
| Q01082;P11277 | Q01082 | Spectrin beta chain, non-erythrocytic 1 |
| Q01469;A8MUU1 | Q01469 | Fatty acid-binding protein 5 |

| | | |
|---|---|---|
| Q01518 | Q01518 | Adenylyl cyclase-associated protein 1 |
| Q01581 | Q01581 | Hydroxymethylglutaryl-CoA synthase, cytoplasmic |
| Q01813 | Q01813 | ATP-dependent 6-phosphofructokinase, platelet type |
| Q02413 | Q02413 | Desmoglein-1 |
| Q02543 | Q02543 | Large ribosomal subunit protein eL20 |
| Q02750 | Q02750 | Dual specificity mitogen-activated protein kinase kinase 1 |
| Q02790 | Q02790 | Peptidyl-prolyl cis-trans isomerase FKBP4;Peptidyl-prolyl c |
| Q02878 | Q02878 | Large ribosomal subunit protein eL6 |
| Q02978 | Q02978 | Mitochondrial 2-oxoglutarate/malate carrier protein |
| Q03013 | Q03013 | Glutathione S-transferase Mu 4 |
| Q03154 | Q03154 | Aminoacylase-1 |
| Q03169 | Q03169 | Tumor necrosis factor alpha-induced protein 2 |
| Q03252 | Q03252 | Lamin-B2 |
| Q03518 | Q03518 | Antigen peptide transporter 1 |
| Q04323 | Q04323 | UBX domain-containing protein 1 |
| Q04446 | Q04446 | 1,4-alpha-glucan-branching enzyme |
| Q04637 | Q04637 | Eukaryotic translation initiation factor 4 gamma 1 |
| Q04760 | Q04760 | Lactoylglutathione lyase |
| Q04828;P17516 | Q04828 | Aldo-keto reductase family 1 member C1 |
| Q04837 | Q04837 | Single-stranded DNA-binding protein, mitochondrial |
| Q04917 | Q04917 | 14-3-3 protein eta |
| Q05655;Q04759 | Q05655 | Protein kinase C delta type;Protein kinase C delta type regu |
| Q06124 | Q06124 | Tyrosine-protein phosphatase non-receptor type 11 |
| Q06210;O94808 | Q06210 | Glutamine--fructose-6-phosphate aminotransferase [isom |
| Q06323 | Q06323 | Proteasome activator complex subunit 1 |
| Q06830 | Q06830 | Peroxiredoxin-1 |
| Q07020 | Q07020 | Large ribosomal subunit protein eL18 |
| Q07021 | Q07021 | Complement component 1 Q subcomponent-binding prote |
| Q07065 | Q07065 | Cytoskeleton-associated protein 4 |
| Q07617 | Q07617 | Sperm-associated antigen 1 |
| Q07812 | Q07812 | Apoptosis regulator BAX |
| Q07955 | Q07955 | Serine/arginine-rich splicing factor 1 |
| Q07960 | Q07960 | Rho GTPase-activating protein 1 |
| Q08117 | Q08117 | TLE family member 5 |
| Q08211 | Q08211 | ATP-dependent RNA helicase A |
| Q08257 | Q08257 | Quinone oxidoreductase |
| Q08380 | Q08380 | Galectin-3-binding protein |
| Q08623 | Q08623 | Pseudouridine-5-phosphatase |
| Q08752 | Q08752 | Peptidyl-prolyl cis-trans isomerase D |
| Q09666 | Q09666 | Neuroblast differentiation-associated protein AHNAK |
| Q0VF49 | Q0VF49 | Uncharacterized protein KIAA2012 |
| Q0VFZ6 | Q0VFZ6 | Cilia- and flagella- associated protein 210 |
| Q10471 | Q10471 | Polypeptide N-acetylgalactosaminyltransferase 2;Polypept |
| Q10567 | Q10567 | AP-1 complex subunit beta-1 |
| Q10588 | Q10588 | ADP-ribosyl cyclase/cyclic ADP-ribose hydrolase 2 |
| Q12765 | Q12765 | Secernin-1 |
| Q12792 | Q12792 | Twinfilin-1 |

| | | |
|---|---|---|
| Q12805 | Q12805 | EGF-containing fibulin-like extracellular matrix protein 1 |
| Q12904 | Q12904 | Aminoacyl tRNA synthase complex-interacting multifunctic |
| Q12905 | Q12905 | Interleukin enhancer-binding factor 2 |
| Q12906 | Q12906 | Interleukin enhancer-binding factor 3 |
| Q12907 | Q12907 | Vesicular integral-membrane protein VIP36 |
| Q12931 | Q12931 | Heat shock protein 75 kDa, mitochondrial |
| Q12959 | Q12959 | Disks large homolog 1 |
| Q12965 | Q12965 | Unconventional myosin-Ie |
| Q13045 | Q13045 | Protein flightless-1 homolog |
| Q13057 | Q13057 | Bifunctional coenzyme A synthase;Phosphopantetheine ac |
| Q13098 | Q13098 | COP9 signalosome complex subunit 1 |
| Q13099 | Q13099 | Intraflagellar transport protein 88 homolog |
| Q13126 | Q13126 | S-methyl-5-thioadenosine phosphorylase |
| Q13131;P54646 | Q13131 | 5-AMP-activated protein kinase catalytic subunit alpha-1 |
| Q13148 | Q13148 | TAR DNA-binding protein 43 |
| Q13153 | Q13153 | Serine/threonine-protein kinase PAK 1 |
| Q13155 | Q13155 | Aminoacyl tRNA synthase complex-interacting multifunctic |
| Q13162 | Q13162 | Peroxiredoxin-4 |
| Q13177;O75914 | Q13177 | Serine/threonine-protein kinase PAK 2;PAK-2p27;PAK-2p3 |
| Q13183 | Q13183 | Solute carrier family 13 member 2 |
| Q13200 | Q13200 | 26S proteasome non-ATPase regulatory subunit 2 |
| Q13228 | Q13228 | Methanethiol oxidase |
| Q13247;Q08170;Q132 | Q13247;Q08170 | Serine/arginine-rich splicing factor 6;Serine/arginine-rich s |
| Q13263 | Q13263 | Transcription intermediary factor 1-beta |
| Q13277 | Q13277 | Syntaxin-3 |
| Q13283;Q9UN86 | Q13283 | Ras GTPase-activating protein-binding protein 1 |
| Q13393 | Q13393 | Phospholipase D1 |
| Q13404 | Q13404 | Ubiquitin-conjugating enzyme E2 variant 1 |
| Q13409 | Q13409 | Cytoplasmic dynein 1 intermediate chain 2 |
| Q13421 | Q13421 | Mesothelin;Megakaryocyte-potentiating factor;Mesothelin, |
| Q13442 | Q13442 | 28 kDa heat- and acid-stable phosphoprotein |
| Q13509 | Q13509 | Tubulin beta-3 chain |
| Q13510 | Q13510 | Acid ceramidase;Acid ceramidase subunit alpha;Acid cerai |
| Q13526;O15428 | Q13526 | Peptidyl-prolyl cis-trans isomerase NIMA-interacting 1 |
| Q13561 | Q13561 | Dynactin subunit 2 |
| Q13564 | Q13564 | NEDD8-activating enzyme E1 regulatory subunit |
| Q13596 | Q13596 | Sorting nexin-1 |
| Q13618 | Q13618 | Cullin-3 |
| Q13630 | Q13630 | GDP-L-fucose synthase |
| Q13724 | Q13724 | Mannosyl-oligosaccharide glucosidase |
| Q13740 | Q13740 | CD166 antigen |
| Q13751 | Q13751 | Laminin subunit beta-3 |
| Q13753 | Q13753 | Laminin subunit gamma-2 |
| Q13813 | Q13813 | Spectrin alpha chain, non-erythrocytic 1 |
| Q13838;O00148 | Q13838;O00148 | Spliceosome RNA helicase DDX39B;ATP-dependent RNA h |
| Q13867 | Q13867 | Bleomycin hydrolase |
| Q13885 | Q13885 | Tubulin beta-2A chain |

| | | |
|---|---|---|
| Q13907;Q9BXS1 | Q13907 | Isopentenyl-diphosphate Delta-isomerase 1 |
| Q13938 | Q13938 | Calcyphosin |
| Q14019 | Q14019 | Coactosin-like protein |
| Q14103 | Q14103 | Heterogeneous nuclear ribonucleoprotein D0 |
| Q14108 | Q14108 | Lysosome membrane protein 2 |
| Q14116 | Q14116 | Interleukin-18 |
| Q14134 | Q14134 | Tripartite motif-containing protein 29 |
| Q14145 | Q14145 | Kelch-like ECH-associated protein 1 |
| Q14166 | Q14166 | Tubulin--tyrosine ligase-like protein 12 |
| Q14204 | Q14204 | Cytoplasmic dynein 1 heavy chain 1 |
| Q14210 | Q14210 | Lymphocyte antigen 6D |
| Q14240 | Q14240 | Eukaryotic initiation factor 4A-II |
| Q14244 | Q14244 | Ensconsin |
| Q14247 | Q14247 | Src substrate cortactin |
| Q14254 | Q14254 | Flotillin-2 |
| Q14258 | Q14258 | E3 ubiquitin/ISG15 ligase TRIM25 |
| Q14344 | Q14344 | Guanine nucleotide-binding protein subunit alpha-13 |
| Q14353 | Q14353 | Guanidinoacetate N-methyltransferase |
| Q14376 | Q14376 | UDP-glucose 4-epimerase |
| Q14508 | Q14508 | WAP four-disulfide core domain protein 2 |
| Q14558 | Q14558 | Phosphoribosyl pyrophosphate synthase-associated protei |
| Q14573 | Q14573 | Inositol 1,4,5-trisphosphate receptor type 3 |
| Q14651 | Q14651 | Plastin-1 |
| Q14677 | Q14677 | Clathrin interactor 1 |
| Q14697 | Q14697 | Neutral alpha-glucosidase AB |
| Q14728 | Q14728 | Major facilitator superfamily domain-containing protein 10 |
| Q14738;Q13362 | Q14738 | Serine/threonine-protein phosphatase 2A 56 kDa regulator |
| Q14764 | Q14764 | Major vault protein |
| Q14790 | Q14790 | Caspase-8;Caspase-8 subunit p18;Caspase-8 subunit p10 |
| Q14847 | Q14847 | LIM and SH3 domain protein 1 |
| Q14894 | Q14894 | Ketimine reductase mu-crystallin |
| Q14914 | Q14914 | Prostaglandin reductase 1 |
| Q14956 | Q14956 | Transmembrane glycoprotein NMB |
| Q14974 | Q14974 | Importin subunit beta-1 |
| Q14C86 | Q14C86 | GTPase-activating protein and VPS9 domain-containing pro |
| Q14CN2 | Q14CN2 | Calcium-activated chloride channel regulator 4;Calcium-a |
| Q15008 | Q15008 | 26S proteasome non-ATPase regulatory subunit 6 |
| Q15019 | Q15019 | Septin-2 |
| Q15046 | Q15046 | Lysine--tRNA ligase |
| Q15056 | Q15056 | Eukaryotic translation initiation factor 4H |
| Q15084 | Q15084 | Protein disulfide-isomerase A6 |
| Q15102 | Q15102 | Platelet-activating factor acetylhydrolase IB subunit alpha1 |
| Q15121 | Q15121 | Astrocytic phosphoprotein PEA-15 |
| Q15125 | Q15125 | 3-beta-hydroxysteroid-Delta(8),Delta(7)-isomerase |
| Q15126 | Q15126 | Phosphomevalonate kinase |
| Q15149 | Q15149 | Plectin |
| Q15154 | Q15154 | Pericentriolar material 1 protein |

| | | |
|---|---|---|
| Q15172 | Q15172 | Serine/threonine-protein phosphatase 2A 56 kDa regulator |
| Q15181 | Q15181 | Inorganic pyrophosphatase |
| Q15185 | Q15185 | Prostaglandin E synthase 3 |
| Q15208;Q9Y2H1 | Q15208 | Serine/threonine-protein kinase 38 |
| Q15233 | Q15233 | Non-POU domain-containing octamer-binding protein |
| Q15257 | Q15257 | Serine/threonine-protein phosphatase 2A activator |
| Q15286 | Q15286 | Ras-related protein Rab-35 |
| Q15293 | Q15293 | Reticulocalbin-1 |
| Q15363 | Q15363 | Transmembrane emp24 domain-containing protein 2 |
| Q15365;P57723 | Q15365 | Poly(rC)-binding protein 1 |
| Q15366 | Q15366 | Poly(rC)-binding protein 2 |
| Q15369 | Q15369 | Elongin-C |
| Q15370 | Q15370 | Elongin-B |
| Q15392 | Q15392 | Delta(24)-sterol reductase |
| Q15393 | Q15393 | Splicing factor 3B subunit 3 |
| Q15417 | Q15417 | Calponin-3 |
| Q15418 | Q15418 | Ribosomal protein S6 kinase alpha-1 |
| Q15435 | Q15435 | Protein phosphatase 1 regulatory subunit 7 |
| Q15437 | Q15437 | Protein transport protein Sec23B |
| Q15506 | Q15506 | Sperm surface protein Sp17 |
| Q15599 | Q15599 | Na(+)/H(+) exchange regulatory cofactor NHE-RF2 |
| Q15628 | Q15628 | Tumor necrosis factor receptor type 1-associated DEATH d |
| Q15631 | Q15631 | Translin |
| Q15691 | Q15691 | Microtubule-associated protein RP/EB family member 1 |
| Q15714 | Q15714 | TSC22 domain family protein 1 |
| Q15746 | Q15746 | Myosin light chain kinase, smooth muscle;Myosin light cha |
| Q15773 | Q15773 | Myeloid leukemia factor 2 |
| Q15785 | Q15785 | Mitochondrial import receptor subunit TOM34 |
| Q15796;P84022;Q997 | Q15796;P84022;Q99 | Mothers against decapentaplegic homolog 2;Mothers agai |
| Q15819 | Q15819 | Ubiquitin-conjugating enzyme E2 variant 2 |
| Q15833 | Q15833 | Syntaxin-binding protein 2 |
| Q15843 | Q15843 | NEDD8 |
| Q15847 | Q15847 | Adipogenesis regulatory factor |
| Q15907;P62491 | Q15907;P62491 | Ras-related protein Rab-11B;Ras-related protein Rab-11A |
| Q16181;Q6ZU15 | Q16181 | Septin-7 |
| Q16186 | Q16186 | Proteasomal ubiquitin receptor ADRM1 |
| Q16222;Q3KQV9 | Q16222 | UDP-N-acetylhexosamine pyrophosphorylase |
| Q16270 | Q16270 | Insulin-like growth factor-binding protein 7 |
| Q16348 | Q16348 | Solute carrier family 15 member 2 |
| Q16401 | Q16401 | 26S proteasome non-ATPase regulatory subunit 5 |
| Q16531 | Q16531 | DNA damage-binding protein 1 |
| Q16537;Q15173 | Q16537 | Serine/threonine-protein phosphatase 2A 56 kDa regulator |
| Q16543 | Q16543 | Hsp90 co-chaperone Cdc37;Hsp90 co-chaperone Cdc37, |
| Q16563 | Q16563 | Synaptophysin-like protein 1 |
| Q16610 | Q16610 | Extracellular matrix protein 1 |
| Q16651 | Q16651 | Prostasin;Prostasin light chain;Prostasin heavy chain |
| Q16658 | Q16658 | Fascin |

| | | |
|---|---|---|
| Q16666 | Q16666 | Gamma-interferon-inducible protein 16 |
| Q16674 | Q16674 | Melanoma-derived growth regulatory protein |
| Q16698 | Q16698 | 2,4-dienoyl-CoA reductase [(3E)-enoyl-CoA-producing], mi |
| Q16718 | Q16718 | NADH dehydrogenase [ubiquinone] 1 alpha subcomplex su |
| Q16719 | Q16719 | Kynureninase |
| Q16762 | Q16762 | Thiosulfate sulfurtransferase |
| Q16772 | Q16772 | Glutathione S-transferase A3 |
| Q16774 | Q16774 | Guanylate kinase |
| Q16775 | Q16775 | Hydroxyacylglutathione hydrolase, mitochondrial |
| Q16795 | Q16795 | NADH dehydrogenase [ubiquinone] 1 alpha subcomplex su |
| Q16822;P35558 | Q16822 | Phosphoenolpyruvate carboxykinase [GTP], mitochondrial |
| Q16836 | Q16836 | Hydroxyacyl-coenzyme A dehydrogenase, mitochondrial |
| Q16851 | Q16851 | UTP--glucose-1-phosphate uridylyltransferase |
| Q16881;Q9NNW7 | Q16881 | Thioredoxin reductase 1, cytoplasmic |
| Q16890 | Q16890 | Tumor protein D53 |
| Q16891 | Q16891 | MICOS complex subunit MIC60 |
| Q1KMD3 | Q1KMD3 | Heterogeneous nuclear ribonucleoprotein U-like protein 2 |
| Q1MSJ5 | Q1MSJ5 | Centrosome and spindle pole-associated protein 1 |
| Q2M329 | Q2M329 | Coiled-coil domain-containing protein 96 |
| Q2TAA2 | Q2TAA2 | Isoamyl acetate-hydrolyzing esterase 1 homolog |
| Q2TAC6 | Q2TAC6 | Kinesin-like protein KIF19 |
| Q32MZ4 | Q32MZ4 | Leucine-rich repeat flightless-interacting protein 1 |
| Q3B820 | Q3B820 | Protein FAM161A |
| Q3LXA3 | Q3LXA3 | Triokinase/FMN cyclase;ATP-dependent dihydroxyacetone |
| Q3SXY8 | Q3SXY8 | ADP-ribosylation factor-like protein 13B |
| Q3SYG4 | Q3SYG4 | Protein PTHB1 |
| Q3ZCM7 | Q3ZCM7 | Tubulin beta-8 chain |
| Q494V2 | Q494V2 | Cilia- and flagella-associated protein 100 |
| Q49MG5 | Q49MG5 | Microtubule-associated protein 9 |
| Q4G0A6 | Q4G0A6 | Probable ubiquitin carboxyl-terminal hydrolase MINDY-4 |
| Q4G0P3 | Q4G0P3 | Hydrocephalus-inducing protein homolog |
| Q4G0U5 | Q4G0U5 | Cilia- and flagella-associated protein 221 |
| Q4G0X9 | Q4G0X9 | Coiled-coil domain-containing protein 40 |
| Q4LDG9 | Q4LDG9 | Dynein axonemal light chain 1 |
| Q502W6 | Q502W6 | von Willebrand factor A domain-containing protein 3B |
| Q502X0 | Q502X0 | MORN repeat-containing protein 2 |
| Q53EL6 | Q53EL6 | Programmed cell death protein 4 |
| Q53EV4 | Q53EV4 | Leucine-rich repeat-containing protein 23 |
| Q53FA7 | Q53FA7 | Quinone oxidoreductase PIG3 |
| Q53FT3 | Q53FT3 | Protein Hikeshi |
| Q53GD3 | Q53GD3 | Choline transporter-like protein 4 |
| Q53GQ0 | Q53GQ0 | Very-long-chain 3-oxoacyl-CoA reductase |
| Q53H82 | Q53H82 | Endoribonuclease LACTB2 |
| Q53H96 | Q53H96 | Pyrroline-5-carboxylate reductase 3 |
| Q53S08 | Q53S08 | Ras-related protein Rab-6D |
| Q569K6 | Q569K6 | Coiled-coil domain-containing protein 157 |
| Q56VL3 | Q56VL3 | OCIA domain-containing protein 2 |

| | | |
|---|---|---|
| Q58FF6 | Q58FF6 | Putative heat shock protein HSP 90-beta 4 |
| Q58FF8 | Q58FF8 | Putative heat shock protein HSP 90-beta 2 |
| Q5BJF6 | Q5BJF6 | Outer dense fiber protein 2 |
| Q5BN46 | Q5BN46 | Piercer of microtubule wall 1 protein |
| Q5I0G3 | Q5I0G3 | Putative malate dehydrogenase 1B |
| Q5JR98 | Q5JR98 | Dynein light chain Tctex-type 4 |
| Q5JSH3 | Q5JSH3 | WD repeat-containing protein 44 |
| Q5JST6 | Q5JST6 | EF-hand domain-containing family member C2 |
| Q5JTN6 | Q5JTN6 | WD repeat-containing protein 38 |
| Q5JU00 | Q5JU00 | Dynein regulatory complex subunit 5 |
| Q5JU67 | Q5JU67 | Cilia- and flagella-associated protein 157 |
| Q5JVL4 | Q5JVL4 | EF-hand domain-containing protein 1 |
| Q5R3I4 | Q5R3I4 | Tetratricopeptide repeat protein 38 |
| Q5RHP9 | Q5RHP9 | Glutamate-rich protein 3 |
| Q5SQI0 | Q5SQI0 | Alpha-tubulin N-acetyltransferase 1 |
| Q5SRE7 | Q5SRE7 | Phytanoyl-CoA dioxygenase domain-containing protein 1 |
| Q5SXH7 | Q5SXH7 | Pleckstrin homology domain-containing family S member 1 |
| Q5T089 | Q5T089 | MORN repeat-containing protein 1 |
| Q5T0N1 | Q5T0N1 | Cilia- and flagella-associated protein 70 |
| Q5T0Z8 | Q5T0Z8 | Uncharacterized protein C6orf132 |
| Q5T124 | Q5T124 | UBX domain-containing protein 11 |
| Q5T2S8 | Q5T2S8 | Outer dynein arm-docking complex subunit 2 |
| Q5T5A4 | Q5T5A4 | Cilia- and flagella-associated protein 276 |
| Q5T601 | Q5T601 | Adhesion G-protein coupled receptor F1 |
| Q5T655;O95822 | Q5T655 | Cilia- and flagella-associated protein 58 |
| Q5T6J7 | Q5T6J7 | Probable gluconokinase |
| Q5T6V5 | Q5T6V5 | Queuosine 5-phosphate N-glycosylase/hydrolase |
| Q5T700 | Q5T700 | Low-density lipoprotein receptor class A domain-containin |
| Q5T749 | Q5T749 | Keratinocyte proline-rich protein |
| Q5TBC7 | Q5TBC7 | Bcl-2-like protein 15 |
| Q5TCS8 | Q5TCS8 | Adenylate kinase 9 |
| Q5TD94 | Q5TD94 | Radial spoke head protein 4 homolog A |
| Q5TDH0;Q8WTU0 | Q5TDH0 | Protein DDI1 homolog 2 |
| Q5TEC6 | Q5TEC6 | Histone H3-7 |
| Q5TFE4 | Q5TFE4 | 5-nucleotidase domain-containing protein 1 |
| Q5TH74 | Q5TH74 | O(6)-methylguanine-induced apoptosis 2 |
| Q5THR3 | Q5THR3 | EF-hand calcium-binding domain-containing protein 6 |
| Q5TID7 | Q5TID7 | Coiled-coil domain-containing protein 181 |
| Q5TZA2;Q86T23;Q8IVE | Q5TZA2 | Rootletin |
| Q5TZF3 | Q5TZF3 | Ankyrin repeat domain-containing protein 45 |
| Q5VTH2 | Q5VTH2 | Protein Flattop |
| Q5VTH9 | Q5VTH9 | Dynein axonemal intermediate chain 4 |
| Q5VTT2 | Q5VTT2 | Cilia- and flagella-associated protein 95 |
| Q5VU69 | Q5VU69 | Cilia- and flagella-associated protein 141 |
| Q5VUJ9 | Q5VUJ9 | Dynein regulatory complex protein 8 |
| Q5VVH5 | Q5VVH5 | Interleukin-1 receptor-associated kinase 1-binding protein |
| Q5VW32 | Q5VW32 | BRO1 domain-containing protein BROX |

| | | |
|---|---|---|
| Q5VWZ2 | Q5VWZ2 | Lysophospholipase-like protein 1 |
| Q5VZ52 | Q5VZ52 | MORN repeat-containing protein 5 |
| Q5W041 | Q5W041 | Armadillo repeat-containing protein 3 |
| Q5XUX1 | Q5XUX1 | F-box/WD repeat-containing protein 9 |
| Q658L1 | Q658L1 | Stabilizer of axonemal microtubules 2 |
| Q687X5 | Q687X5 | Metalloreductase STEAP4 |
| Q6AI08 | Q6AI08 | HEAT repeat-containing protein 6 |
| Q6BCY4 | Q6BCY4 | NADH-cytochrome b5 reductase 2 |
| Q6DD88 | Q6DD88 | Atlastin-3 |
| Q6DKJ4 | Q6DKJ4 | Nucleoredoxin |
| Q6FI81 | Q6FI81 | Anamorsin |
| Q6GMV3 | Q6GMV3 | Putative peptidyl-tRNA hydrolase PTRHD1 |
| Q6IBS0 | Q6IBS0 | Twinfilin-2 |
| Q6IPM2 | Q6IPM2 | IQ domain-containing protein E |
| Q6MZZ7 | Q6MZZ7 | Calpain-13 |
| Q6N063 | Q6N063 | 2-oxoglutarate and iron-dependent oxygenase domain-con |
| Q6NUK1 | Q6NUK1 | Mitochondrial adenyl nucleotide antiporter SLC25A24 |
| Q6NUM9 | Q6NUM9 | All-trans-retinol 13,14-reductase |
| Q6NUN7 | Q6NUN7 | Jhy protein homolog |
| Q6NVY1 | Q6NVY1 | 3-hydroxyisobutyryl-CoA hydrolase, mitochondrial |
| Q6P2C0 | Q6P2C0 | WD repeat-containing protein 93 |
| Q6P2Q9 | Q6P2Q9 | Pre-mRNA-processing-splicing factor 8 |
| Q6P3R8 | Q6P3R8 | Serine/threonine-protein kinase Nek5 |
| Q6P656 | Q6P656 | Cilia- and flagella-associated protein 161 |
| Q6P6B1 | Q6P6B1 | Glutamate-rich protein 5 |
| Q6P996;Q6P474 | Q6P996 | Pyridoxal-dependent decarboxylase domain-containing pro |
| Q6P9G0 | Q6P9G0 | Cytochrome b5 domain-containing protein 1 |
| Q6PCE3 | Q6PCE3 | Glucose 1,6-bisphosphate synthase |
| Q6PF18 | Q6PF18 | MORN repeat-containing protein 3 |
| Q6PI97 | Q6PI97 | Cilia- and flagella-associated protein HOATZ |
| Q6PIY5 | Q6PIY5 | Armadillo-like helical domain containing protein 1 |
| Q6Q759 | Q6Q759 | Sperm-associated antigen 17 |
| Q6S9Z5 | Q6S9Z5 | Zinc finger protein 474 |
| Q6TDU7;Q9H4E7 | Q6TDU7 | Dynein axonemal intermediate chain 7 |
| Q6UW68 | Q6UW68 | Transmembrane protein 205 |
| Q6UX06 | Q6UX06 | Olfactomedin-4 |
| Q6UXB2 | Q6UXB2 | C-X-C motif chemokine 17;4-Cys CXCL17 |
| Q6UXB3 | Q6UXB3 | Ly6/PLAUR domain-containing protein 2 |
| Q6UXY8 | Q6UXY8 | Transmembrane channel-like protein 5 |
| Q6V702;Q14185 | Q6V702 | Cilia- and flagella-associated protein 299 |
| Q6XQN6 | Q6XQN6 | Nicotinate phosphoribosyltransferase |
| Q6YN16 | Q6YN16 | Hydroxysteroid dehydrogenase-like protein 2 |
| Q6YP21 | Q6YP21 | Kynurenine--oxoglutarate transaminase 3 |
| Q6ZMT9 | Q6ZMT9 | Death domain-containing protein 1 |
| Q6ZN30 | Q6ZN30 | Zinc finger protein basonuclin-2 |
| Q6ZN66 | Q6ZN66 | Guanylate-binding protein 6 |
| Q6ZQR2 | Q6ZQR2 | Cilia- and flagella-associated protein 77 |

| | | |
|---|---|---|
| Q6ZQY2 | Q6ZQY2 | Leucine-rich repeat-containing protein 74B |
| Q6ZR08 | Q6ZR08 | Dynein axonemal heavy chain 12 |
| Q6ZTQ4 | Q6ZTQ4 | Cadherin-related family member 3 |
| Q6ZTR5 | Q6ZTR5 | Cilia- and flagella-associated protein 47 |
| Q6ZTR7 | Q6ZTR7 | CBY1-interacting BAR domain-containing protein 2 |
| Q6ZU64 | Q6ZU64 | Cilia- and flagella-associated protein 65 |
| Q6ZUG5 | Q6ZUG5 | Uncharacterized protein FLJ43738 |
| Q6ZUX3 | Q6ZUX3 | TOG array regulator of axonemal microtubules protein 2 |
| Q6ZV65 | Q6ZV65 | Protein FAM47E |
| Q6ZVX7 | Q6ZVX7 | F-box only protein 50 |
| Q709C8 | Q709C8 | Intermembrane lipid transfer protein VPS13C |
| Q71DI3 | Q71DI3 | Histone H3.2 |
| Q71U36;P0DPH8;P0DF | Q71U36;P0DPH8;P0I | Tubulin alpha-1A chain;Detyrosinated tubulin alpha-1A cha |
| Q7KZF4 | Q7KZF4 | Staphylococcal nuclease domain-containing protein 1 |
| Q7L1Q6 | Q7L1Q6 | eIF5-mimic protein 2 |
| Q7L266 | Q7L266 | Isoaspartyl peptidase/L-asparaginase;Isoaspartyl peptidas |
| Q7L576;Q96F07 | Q7L576 | Cytoplasmic FMR1-interacting protein 1 |
| Q7L5N1 | Q7L5N1 | COP9 signalosome complex subunit 6 |
| Q7L5Y1 | Q7L5Y1 | Mitochondrial enolase superfamily member 1 |
| Q9H8S9;Q7L9L4 | Q9H8S9;Q7L9L4 | MOB kinase activator 1A;MOB kinase activator 1B |
| Q7LBR1 | Q7LBR1 | Charged multivesicular body protein 1b |
| Q7LG56 | Q7LG56 | Ribonucleoside-diphosphate reductase subunit M2 B |
| Q7Z2K6 | Q7Z2K6 | Endoplasmic reticulum metallopeptidase 1 |
| Q7Z2Y8 | Q7Z2Y8 | Interferon-induced very large GTPase 1 |
| Q7Z3E5 | Q7Z3E5 | LisH domain-containing protein ARMC9 |
| Q7Z404 | Q7Z404 | Transmembrane channel-like protein 4 |
| Q7Z406 | Q7Z406 | Myosin-14 |
| Q7Z4G1 | Q7Z4G1 | COMM domain-containing protein 6 |
| Q7Z4H3 | Q7Z4H3 | 5-deoxynucleotidase HDDC2 |
| Q7Z4L5 | Q7Z4L5 | Tetratricopeptide repeat protein 21B |
| Q7Z4S6;O75037 | Q7Z4S6 | Kinesin-like protein KIF21A |
| Q7Z4T9;A1A5B4 | Q7Z4T9 | Cilia- and flagella-associated protein 91 |
| Q7Z4W1 | Q7Z4W1 | L-xylulose reductase |
| Q7Z5L0 | Q7Z5L0 | Vitelline membrane outer layer protein 1 homolog |
| Q7Z5V6 | Q7Z5V6 | Stabilizer of axonemal microtubules 4 |
| Q7Z7H3 | Q7Z7H3 | Ciliogenesis-associated TTC17-interacting protein |
| Q7Z7H5 | Q7Z7H5 | Transmembrane emp24 domain-containing protein 4 |
| Q7Z7M9 | Q7Z7M9 | Polypeptide N-acetylgalactosaminyltransferase 5 |
| Q86TC9;Q8WX93 | Q86TC9;Q8WX93 | Myopalladin;Palladin |
| Q86TX2;P49753;Q3I5F | Q86TX2;P49753 | Acyl-coenzyme A thioesterase 1;Acyl-coenzyme A thioeste |
| Q86UC2 | Q86UC2 | Radial spoke head protein 3 homolog |
| Q86UN6 | Q86UN6 | A-kinase anchor protein 14 |
| Q86UP2 | Q86UP2 | Kinectin |
| Q86UP3 | Q86UP3 | Zinc finger homeobox protein 4 |
| Q86UQ4 | Q86UQ4 | ATP-binding cassette sub-family A member 13 |
| Q86V21 | Q86V21 | Acetoacetyl-CoA synthetase |
| Q86VP6;O75155 | Q86VP6 | Cullin-associated NEDD8-dissociated protein 1 |

| | | |
|---|---|---|
| Q86VQ0 | Q86VQ0 | Lebercilin |
| Q86WT1 | Q86WT1 | Intraflagellar transport protein 70A |
| Q86X29 | Q86X29 | Lipolysis-stimulated lipoprotein receptor |
| Q86X76 | Q86X76 | Deaminated glutathione amidase |
| Q86XH1;A6NCM1 | Q86XH1 | Dynein regulatory complex protein 11 |
| Q86XW9 | Q86XW9 | Thioredoxin domain-containing protein 6 |
| Q86Y56 | Q86Y56 | Dynein axonemal assembly factor 5 |
| Q8IU85;Q14012 | Q8IU85;Q14012 | Calcium/calmodulin-dependent protein kinase type 1D;Ca |
| Q8IV35 | Q8IV35 | WD repeat-containing protein 49 |
| Q8IV56 | Q8IV56 | Proline-rich protein 15 |
| Q8IVF4;REV__Q9Y5K6 | Q8IVF4 | Dynein axonemal heavy chain 10 |
| Q8IVU9 | Q8IVU9 | Ciliary-associated calcium-binding coiled-coil protein 1 |
| Q8IWA5 | Q8IWA5 | Choline transporter-like protein 2 |
| Q8IWE2 | Q8IWE2 | Protein NOXP20 |
| Q8IWG1 | Q8IWG1 | Dynein axonemal intermediate chain 3 |
| Q8IWZ6 | Q8IWZ6 | Bardet-Biedl syndrome 7 protein |
| Q8IXQ6 | Q8IXQ6 | Protein mono-ADP-ribosyltransferase PARP9 |
| Q8IXS2 | Q8IXS2 | Dynein regulatory complex subunit 2 |
| Q8IXY8 | Q8IXY8 | Probable inactive peptidyl-prolyl cis-trans isomerase-like 6 |
| Q8IXZ2 | Q8IXZ2 | Zinc finger CCCH domain-containing protein 3 |
| Q8IY31 | Q8IY31 | Intraflagellar transport protein 20 homolog |
| Q8IY82 | Q8IY82 | Dynein regulatory complex subunit 7 |
| Q8IYE0 | Q8IYE0 | Coiled-coil domain-containing protein 146 |
| Q8IYE1 | Q8IYE1 | Coiled-coil domain-containing protein 13 |
| Q8IYR0 | Q8IYR0 | Cilia- and flagella-associated protein 206 |
| Q8IYT3 | Q8IYT3 | Coiled-coil domain-containing protein 170 |
| Q8IYT4 | Q8IYT4 | Katanin p60 ATPase-containing subunit A-like 2 |
| Q8IYW2 | Q8IYW2 | Cilia- and flagella-associated protein 46 |
| Q8IZ02 | Q8IZ02 | Leucine-rich repeat-containing protein 34 |
| Q8IZ07;Q6ZTN6 | Q8IZ07 | Ankyrin repeat domain-containing protein 13A |
| Q8IZ83 | Q8IZ83 | Aldehyde dehydrogenase family 16 member A1 |
| Q8IZL9 | Q8IZL9 | Cyclin-dependent kinase 20 |
| Q8IZV5 | Q8IZV5 | Retinol dehydrogenase 10 |
| Q8N0U7 | Q8N0U7 | Uncharacterized protein C1orf87 |
| Q8N0W5 | Q8N0W5 | IQ domain-containing protein K |
| Q8N0X2 | Q8N0X2 | Sperm-associated antigen 16 protein |
| Q8N136 | Q8N136 | Dynein assembly factor with WD repeat domains 1 |
| Q8N163 | Q8N163 | Cell cycle and apoptosis regulator protein 2 |
| Q8N1A6 | Q8N1A6 | UPF0462 protein C4orf33 |
| Q8N1D5 | Q8N1D5 | Cilia- and flagella-associated protein 107 |
| Q8N1Q1 | Q8N1Q1 | Carbonic anhydrase 13 |
| Q8N1V2 | Q8N1V2 | Cilia- and flagella-associated protein 52 |
| Q8N309 | Q8N309 | Leucine-rich repeat-containing protein 43 |
| Q8N335;P21695 | Q8N335 | Glycerol-3-phosphate dehydrogenase 1-like protein |
| Q8N392 | Q8N392 | Rho GTPase-activating protein 18 |
| Q8N3F0 | Q8N3F0 | Maturin |
| Q8N3Y7 | Q8N3Y7 | Epidermal retinol dehydrogenase 2 |

| | | |
|---|---|---|
| Q8N443 | Q8N443 | RIB43A-like with coiled-coils protein 1 |
| Q8N456 | Q8N456 | Leucine-rich repeat-containing protein 18 |
| Q8N4P2 | Q8N4P2 | Intraflagellar transport protein 70B |
| Q8N4P6 | Q8N4P6 | Leucine-rich repeat-containing protein 71 |
| Q8N5I2 | Q8N5I2 | Arrestin domain-containing protein 1 |
| Q8N5I3 | Q8N5I3 | Potassium channel regulatory protein |
| Q8N5M4 | Q8N5M4 | Tetratricopeptide repeat protein 9C |
| Q8N5R6 | Q8N5R6 | Coiled-coil domain-containing protein 33 |
| Q8N6N7 | Q8N6N7 | Acyl-CoA-binding domain-containing protein 7 |
| Q8N6Q3 | Q8N6Q3 | CD177 antigen |
| Q8N6V9 | Q8N6V9 | Testis-expressed protein 9 |
| Q8N7L0 | Q8N7L0 | Protein FAM216B |
| Q8N7M0 | Q8N7M0 | Dynein light chain Tctex-type 5 |
| Q8N7U6 | Q8N7U6 | EF-hand domain-containing family member B |
| Q8N7X0 | Q8N7X0 | Androglobin |
| Q8N8I0 | Q8N8I0 | Sterile alpha motif domain-containing protein 12 |
| Q8N8N7 | Q8N8N7 | Prostaglandin reductase 2 |
| Q8N9B4 | Q8N9B4 | Ankyrin repeat domain-containing protein 42 |
| Q8N9N7 | Q8N9N7 | Leucine-rich repeat-containing protein 57 |
| Q8N9Z9 | Q8N9Z9 | Lamin tail domain-containing protein 1 |
| Q8NA54 | Q8NA54 | IQ motif and ubiquitin-like domain-containing protein |
| Q8NA56 | Q8NA56 | Tetratricopeptide repeat protein 29 |
| Q8NBF2 | Q8NBF2 | NHL repeat-containing protein 2 |
| Q8NBJ4 | Q8NBJ4 | Golgi membrane protein 1 |
| Q8NBJ7 | Q8NBJ7 | Inactive C-alpha-formylglycine-generating enzyme 2 |
| Q8NBS9 | Q8NBS9 | Thioredoxin domain-containing protein 5 |
| Q8NC51 | Q8NC51 | SERPINE1 mRNA-binding protein 1 |
| Q8NCM8 | Q8NCM8 | Cytoplasmic dynein 2 heavy chain 1 |
| Q8NCW5 | Q8NCW5 | NAD(P)H-hydrate epimerase |
| Q8NDC0 | Q8NDC0 | MAPK-interacting and spindle-stabilizing protein-like |
| Q8NDH3 | Q8NDH3 | Probable aminopeptidase NPEPL1 |
| Q8NDM7 | Q8NDM7 | Cilia- and flagella-associated protein 43 |
| Q8NDW8 | Q8NDW8 | Tetratricopeptide repeat protein 21A |
| Q8NE09 | Q8NE09 | Regulator of G-protein signaling 22 |
| Q8NE18 | Q8NE18 | Putative methyltransferase NSUN7 |
| Q8NE86 | Q8NE86 | Calcium uniporter protein, mitochondrial |
| Q8NEG2 | Q8NEG2 | Uncharacterized protein C7orf57 |
| Q8NEH6 | Q8NEH6 | Meiosis-specific nuclear structural protein 1 |
| Q8NEN0 | Q8NEN0 | Armadillo repeat-containing protein 2 |
| Q8NEU8;Q9UKG1 | Q8NEU8 | DCC-interacting protein 13-beta |
| Q8NEZ3 | Q8NEZ3 | WD repeat-containing protein 19 |
| Q8NFJ5 | Q8NFJ5 | Retinoic acid-induced protein 3 |
| Q8NFJ9 | Q8NFJ9 | Bardet-Biedl syndrome 1 protein |
| Q8NFL0 | Q8NFL0 | UDP-GlcNAc:betaGal beta-1,3-N-acetylglucosaminyltrans |
| Q8NFU3 | Q8NFU3 | Thiosulfate:glutathione sulfurtransferase |
| Q8NFV4 | Q8NFV4 | sn-1-specific diacylglycerol lipase ABHD11 |
| Q8NHU2 | Q8NHU2 | Cilia- and flagella-associated protein 61 |

| | | |
|---|---|---|
| Q8NI77 | Q8NI77 | Kinesin-like protein KIF18A |
| Q8TAD7 | Q8TAD7 | Overexpressed in colon carcinoma 1 protein |
| Q8TAT6 | Q8TAT6 | Nuclear protein localization protein 4 homolog |
| Q8TAV4 | Q8TAV4 | Stomatin-like protein 3 |
| Q8TAX9 | Q8TAX9 | Gasdermin-B;Gasdermin-B, N-terminal;Gasdermin-B, C-te |
| Q8TBC4 | Q8TBC4 | NEDD8-activating enzyme E1 catalytic subunit |
| Q8TBF2 | Q8TBF2 | Prostamide/prostaglandin F synthase |
| Q8TBX8 | Q8TBX8 | Phosphatidylinositol 5-phosphate 4-kinase type-2 gamma |
| Q8TBY9 | Q8TBY9 | Cilia- and flagella-associated protein 251 |
| Q8TBZ2 | Q8TBZ2 | MYCBP-associated protein |
| Q8TC05 | Q8TC05 | Nuclear protein MDM1 |
| Q8TC12;Q96NR8 | Q8TC12 | Retinol dehydrogenase 11 |
| Q8TC29;REV__Q6ZSJ8 | Q8TC29 | Enkurin |
| Q8TC71 | Q8TC71 | Mitochondria-eating protein |
| Q8TC84 | Q8TC84 | Fibronectin type 3 and ankyrin repeat domains protein 1 |
| Q8TCD5 | Q8TCD5 | 5(3)-deoxyribonucleotidase, cytosolic type |
| Q8TCI5 | Q8TCI5 | Ciliary microtubule-associated protein 3 |
| Q8TCX1 | Q8TCX1 | Cytoplasmic dynein 2 light intermediate chain 1 |
| Q8TD06 | Q8TD06 | Anterior gradient protein 3 |
| Q8TD08 | Q8TD08 | Mitogen-activated protein kinase 15 |
| Q8TD19 | Q8TD19 | Serine/threonine-protein kinase Nek9 |
| Q8TD57 | Q8TD57 | Dynein axonemal heavy chain 3 |
| Q8TDL5 | Q8TDL5 | BPI fold-containing family B member 1 |
| Q8TDR0 | Q8TDR0 | TRAF3-interacting protein 1 |
| Q8TE68 | Q8TE68 | Epidermal growth factor receptor kinase substrate 8-like pr |
| Q8TE73;Q9NRZ9;Q0VD | Q8TE73 | Dynein axonemal heavy chain 5 |
| Q8TE77 | Q8TE77 | Protein phosphatase Slingshot homolog 3 |
| Q8TEX9 | Q8TEX9 | Importin-4 |
| Q8TF09 | Q8TF09 | Dynein light chain roadblock-type 2 |
| Q8TF65 | Q8TF65 | PDZ domain-containing protein GIPC2 |
| Q8WU79 | Q8WU79 | Stromal membrane-associated protein 2 |
| Q8WUH6 | Q8WUH6 | Transmembrane protein 263 |
| Q8WUM4 | Q8WUM4 | Programmed cell death 6-interacting protein |
| Q8WUW1 | Q8WUW1 | Protein BRICK1 |
| Q8WVF1 | Q8WVF1 | Protein OSCP1 |
| Q8WVJ2 | Q8WVJ2 | NudC domain-containing protein 2 |
| Q8WVM8 | Q8WVM8 | Sec1 family domain-containing protein 1 |
| Q8WVS4 | Q8WVS4 | Cytoplasmic dynein 2 intermediate chain 1 |
| Q8WVV4 | Q8WVV4 | Protein POF1B |
| Q8WW22 | Q8WW22 | DnaJ homolog subfamily A member 4 |
| Q8WW24 | Q8WW24 | Tektin-4 |
| Q8WW35 | Q8WW35 | Dynein light chain Tctex-type protein 2B |
| Q8WWF8 | Q8WWF8 | Calcyphosin-like protein |
| Q8WWI1 | Q8WWI1 | LIM domain only protein 7 |
| Q8WXF1 | Q8WXF1 | Paraspeckle component 1 |
| Q8WXI7 | Q8WXI7 | Mucin-16 |
| Q8WXX0 | Q8WXX0 | Dynein axonemal heavy chain 7 |

| | | |
|---|---|---|
| Q8WYA0 | Q8WYA0 | Intraflagellar transport protein 81 homolog |
| Q8WYQ4 | Q8WYQ4 | Uncharacterized protein C22orf15 |
| Q8WYR4 | Q8WYR4 | Radial spoke head 1 homolog |
| Q8WZ42 | Q8WZ42 | Titin |
| Q8WZA0 | Q8WZA0 | Protein LZIC |
| Q8WZA9 | Q8WZA9 | Immunity-related GTPase family Q protein |
| Q92499 | Q92499 | ATP-dependent RNA helicase DDX1 |
| Q92520 | Q92520 | Protein FAM3C |
| Q92530 | Q92530 | Proteasome inhibitor PI31 subunit |
| Q92597 | Q92597 | Protein NDRG1 |
| Q92598 | Q92598 | Heat shock protein 105 kDa |
| Q92616 | Q92616 | Stalled ribosome sensor GCN1 |
| Q92623 | Q92623 | Tetratricopeptide repeat protein 9A |
| Q92630 | Q92630 | Dual specificity tyrosine-phosphorylation-regulated kinase |
| Q92696 | Q92696 | Geranylgeranyl transferase type-2 subunit alpha |
| Q92734 | Q92734 | Protein TFG |
| Q92747 | Q92747 | Actin-related protein 2/3 complex subunit 1A |
| Q92817;A8MZ36 | Q92817 | Envoplakin |
| Q92820 | Q92820 | Gamma-glutamyl hydrolase |
| Q92834 | Q92834 | X-linked retinitis pigmentosa GTPase regulator |
| Q92841 | Q92841 | Probable ATP-dependent RNA helicase DDX17 |
| Q92845 | Q92845 | Kinesin-associated protein 3 |
| Q92882 | Q92882 | Osteoclast-stimulating factor 1 |
| Q92890 | Q92890 | Ubiquitin recognition factor in ER-associated degradation p |
| Q92896 | Q92896 | Golgi apparatus protein 1 |
| Q92900 | Q92900 | Regulator of nonsense transcripts 1 |
| Q92973 | Q92973 | Transportin-1 |
| Q93045;P16949 | Q93045;P16949 | Stathmin-2;Stathmin |
| Q93050 | Q93050 | V-type proton ATPase 116 kDa subunit a 1 |
| Q93052 | Q93052 | Lipoma-preferred partner |
| Q969H8 | Q969H8 | Myeloid-derived growth factor |
| Q969L2 | Q969L2 | Protein MAL2 |
| Q969S9 | Q969S9 | Ribosome-releasing factor 2, mitochondrial |
| Q969V4 | Q969V4 | Tektin-1 |
| Q96A22 | Q96A22 | Uncharacterized protein C11orf52 |
| Q96A49 | Q96A49 | Synapse-associated protein 1 |
| Q96AG4 | Q96AG4 | Leucine-rich repeat-containing protein 59;Leucine-rich rep |
| Q96AJ1 | Q96AJ1 | Clusterin-associated protein 1 |
| Q96AQ1 | Q96AQ1 | Coiled-coil domain-containing protein 74A |
| Q96AT9;Q2QD12 | Q96AT9 | Ribulose-phosphate 3-epimerase |
| Q96BW5 | Q96BW5 | Phosphotriesterase-related protein |
| Q96C19 | Q96C19 | EF-hand domain-containing protein D2 |
| Q96C23 | Q96C23 | Galactose mutarotase |
| Q96C74 | Q96C74 | Ropporin-1-like protein |
| Q96CD0 | Q96CD0 | F-box/LRR-repeat protein 8 |
| Q96CN7 | Q96CN7 | Isochorismatase domain-containing protein 1 |
| Q96CW1 | Q96CW1 | AP-2 complex subunit mu |

| | | |
|---|---|---|
| Q96CX2 | Q96CX2 | BTB/POZ domain-containing protein KCTD12 |
| Q96DG6 | Q96DG6 | Carboxymethylenebutenolidase homolog |
| Q96DT5 | Q96DT5 | Dynein axonemal heavy chain 11 |
| Q96DX5 | Q96DX5 | Ankyrin repeat and SOCS box protein 9 |
| Q96DY2;REV__Q14571 | Q96DY2 | Dynein regulatory complex protein 10 |
| Q96E40 | Q96E40 | Sperm acrosome-associated protein 9 |
| Q96E66 | Q96E66 | Leucine-rich repeat-containing protein 51 |
| Q96EK6 | Q96EK6 | Glucosamine 6-phosphate N-acetyltransferase |
| Q96EX3 | Q96EX3 | Cytoplasmic dynein 2 intermediate chain 2 |
| Q96FJ2 | Q96FJ2 | Dynein light chain 2, cytoplasmic |
| Q96FQ6 | Q96FQ6 | Protein S100-A16 |
| Q96FT9 | Q96FT9 | Intraflagellar transport protein 43 homolog |
| Q96FV0 | Q96FV0 | Leucine-rich repeat-containing protein 46 |
| Q96FV2 | Q96FV2 | Secernin-2 |
| Q96FW1 | Q96FW1 | Ubiquitin thioesterase OTUB1 |
| Q96G03 | Q96G03 | Phosphopentomutase |
| Q96G28 | Q96G28 | Cilia- and flagella-associated protein 36 |
| Q96GE6 | Q96GE6 | Calmodulin-like protein 4 |
| Q96GG9 | Q96GG9 | DCN1-like protein 1 |
| Q96GY0 | Q96GY0 | Zinc finger C2HC domain-containing protein 1A |
| Q96HC4 | Q96HC4 | PDZ and LIM domain protein 5 |
| Q96HE7;Q86YB8 | Q96HE7 | ERO1-like protein alpha |
| Q96I99 | Q96I99 | Succinate--CoA ligase [GDP-forming] subunit beta, mitoch |
| Q96IJ6 | Q96IJ6 | Mannose-1-phosphate guanyltransferase alpha |
| Q96IM9 | Q96IM9 | DPY30 domain-containing protein 2 |
| Q96IU4 | Q96IU4 | Putative protein-lysine deacylase ABHD14B |
| Q96IX5 | Q96IX5 | ATP synthase membrane subunit K, mitochondrial |
| Q96K30 | Q96K30 | RBPJ-interacting and tubulin-associated protein 1 |
| Q96KP4 | Q96KP4 | Cytosolic non-specific dipeptidase |
| Q96L03 | Q96L03 | Spermatogenesis-associated protein 17 |
| Q96LB3 | Q96LB3 | Intraflagellar transport protein 74 homolog |
| Q96LJ8 | Q96LJ8 | UBX domain-containing protein 10 |
| Q96LP2 | Q96LP2 | Protein FAM81B |
| Q96LQ0 | Q96LQ0 | Protein phosphatase 1 regulatory subunit 36 |
| Q96LR7;REV__Q9Y561 | Q96LR7 | Uncharacterized protein C2orf50 |
| Q96LT9 | Q96LT9 | RNA-binding region-containing protein 3 |
| Q96LX7 | Q96LX7 | Coiled-coil domain-containing protein 17 |
| Q96LY2 | Q96LY2 | Coiled-coil domain-containing protein 74B |
| Q96M27 | Q96M27 | Protein PRRC1 |
| Q96M32 | Q96M32 | Adenylate kinase 7 |
| Q96M63 | Q96M63 | Outer dynein arm-docking complex subunit 1 |
| Q96M69 | Q96M69 | Leucine-rich repeat and guanylate kinase domain-containi |
| Q96M91 | Q96M91 | Cilia- and flagella-associated protein 53 |
| Q96M98 | Q96M98 | Parkin coregulated gene protein |
| Q96MA6 | Q96MA6 | Adenylate kinase 8 |
| Q96MC2 | Q96MC2 | Dynein regulatory complex protein 1 |
| Q96MN2 | Q96MN2 | NACHT, LRR and PYD domains-containing protein 4 |

| | | |
|---|---|---|
| Q96MR6 | Q96MR6 | Cilia- and flagella-associated protein 57 |
| Q96MT7 | Q96MT7 | Cilia- and flagella-associated protein 44 |
| Q96MW1 | Q96MW1 | Coiled-coil domain-containing protein 43 |
| Q96MY7 | Q96MY7 | Protein FAM161B |
| Q96N03 | Q96N03 | V-set and transmembrane domain-containing protein 2-like |
| Q96N06 | Q96N06 | Spermatogenesis-associated protein 33 |
| Q96N23 | Q96N23 | Cilia- and flagella-associated protein 54 |
| Q96NG3 | Q96NG3 | Outer dynein arm-docking complex subunit 4 |
| Q96NH3 | Q96NH3 | Protein broad-minded |
| Q96NY7;Q9NZA1;O152 | Q96NY7 | Chloride intracellular channel protein 6 |
| Q96PK6 | Q96PK6 | RNA-binding protein 14 |
| Q96QK1 | Q96QK1 | Vacuolar protein sorting-associated protein 35 |
| Q96RY7 | Q96RY7 | Intraflagellar transport protein 140 homolog |
| Q96S97 | Q96S97 | Myeloid-associated differentiation marker |
| Q96TA1 | Q96TA1 | Protein Niban 2 |
| Q99102 | Q99102 | Mucin-4;Mucin-4 alpha chain;Mucin-4 beta chain |
| Q99417 | Q99417 | c-Myc-binding protein |
| Q99426 | Q99426 | Tubulin-folding cofactor B |
| Q99436 | Q99436 | Proteasome subunit beta type-7 |
| Q99439 | Q99439 | Calponin-2 |
| Q99447 | Q99447 | Ethanolamine-phosphate cytidylyltransferase |
| Q99460 | Q99460 | 26S proteasome non-ATPase regulatory subunit 1 |
| Q99471 | Q99471 | Prefoldin subunit 5 |
| Q99497 | Q99497 | Parkinson disease protein 7 |
| Q99536 | Q99536 | Synaptic vesicle membrane protein VAT-1 homolog |
| Q99584 | Q99584 | Protein S100-A13 |
| Q99598 | Q99598 | Translin-associated protein X |
| Q99615 | Q99615 | DnaJ homolog subfamily C member 7 |
| Q99623 | Q99623 | Prohibitin-2 |
| Q99627 | Q99627 | COP9 signalosome complex subunit 8 |
| Q99685 | Q99685 | Monoglyceride lipase |
| Q99714 | Q99714 | 3-hydroxyacyl-CoA dehydrogenase type-2 |
| Q99729 | Q99729 | Heterogeneous nuclear ribonucleoprotein A/B |
| Q99733 | Q99733 | Nucleosome assembly protein 1-like 4 |
| Q99798 | Q99798 | Aconitate hydratase, mitochondrial |
| Q99805 | Q99805 | Transmembrane 9 superfamily member 2 |
| Q99816 | Q99816 | Tumor susceptibility gene 101 protein |
| Q99828 | Q99828 | Calcium and integrin-binding protein 1 |
| Q99829 | Q99829 | Copine-1 |
| Q99832 | Q99832 | T-complex protein 1 subunit eta;T-complex protein 1 subun |
| Q99836 | Q99836 | Myeloid differentiation primary response protein MyD88 |
| Q99873;Q9NR22 | Q99873 | Protein arginine N-methyltransferase 1 |
| Q99932 | Q99932 | Sperm-associated antigen 8 |
| Q99941 | Q99941 | Cyclic AMP-dependent transcription factor ATF-6 beta;Proc |
| Q99961;Q99962 | Q99961 | Endophilin-A2 |
| Q9BPW8 | Q9BPW8 | Protein NipSnap homolog 1 |
| Q9BPW9 | Q9BPW9 | Dehydrogenase/reductase SDR family member 9 |

| | | |
|---|---|---|
| Q9BPX5 | Q9BPX5 | Actin-related protein 2/3 complex subunit 5-like protein |
| Q9BQE3 | Q9BQE3 | Tubulin alpha-1C chain;Detyrosinated tubulin alpha-1C cha |
| Q9BR76 | Q9BR76 | Coronin-1B |
| Q9BRA2 | Q9BRA2 | Thioredoxin domain-containing protein 17 |
| Q9BRF8 | Q9BRF8 | Serine/threonine-protein phosphatase CPPED1 |
| Q9BRG1 | Q9BRG1 | Vacuolar protein-sorting-associated protein 25 |
| Q9BRP8 | Q9BRP8 | Partner of Y14 and mago |
| Q9BRQ4 | Q9BRQ4 | Cilia- and flagella-associated protein 300 |
| Q9BRT3 | Q9BRT3 | Migration and invasion enhancer 1 |
| Q9BRX8 | Q9BRX8 | Peroxiredoxin-like 2A |
| Q9BS26 | Q9BS26 | Endoplasmic reticulum resident protein 44 |
| Q9BS40 | Q9BS40 | Latexin |
| Q9BT73 | Q9BT73 | Proteasome assembly chaperone 3 |
| Q9BT92 | Q9BT92 | Trichoplein keratin filament-binding protein |
| Q9BTM9 | Q9BTM9 | Ubiquitin-related modifier 1 |
| Q9BTT0 | Q9BTT0 | Acidic leucine-rich nuclear phosphoprotein 32 family mem |
| Q9BTV4 | Q9BTV4 | Transmembrane protein 43 |
| Q9BU02 | Q9BU02 | Thiamine-triphosphatase |
| Q9BUB5 | Q9BUB5 | MAP kinase-interacting serine/threonine-protein kinase 1 |
| Q9BUD6 | Q9BUD6 | Spondin-2 |
| Q9BUF5 | Q9BUF5 | Tubulin beta-6 chain |
| Q9BUJ2 | Q9BUJ2 | Heterogeneous nuclear ribonucleoprotein U-like protein 1 |
| Q9BUL8 | Q9BUL8 | Programmed cell death protein 10 |
| Q9BUP3 | Q9BUP3 | Oxidoreductase HTATIP2 |
| Q9BUT1 | Q9BUT1 | Dehydrogenase/reductase SDR family member 6 |
| Q9BV20 | Q9BV20 | Methylthioribose-1-phosphate isomerase |
| Q9BV36 | Q9BV36 | Melanophilin |
| Q9BV40 | Q9BV40 | Vesicle-associated membrane protein 8 |
| Q9BVA0 | Q9BVA0 | Katanin p80 WD40 repeat-containing subunit B1 |
| Q9BVG4 | Q9BVG4 | Protein PBDC1 |
| Q9BVK6 | Q9BVK6 | Transmembrane emp24 domain-containing protein 9 |
| Q9BVM2 | Q9BVM2 | Protein DPCD |
| Q9BW04 | Q9BW04 | Specifically androgen-regulated gene protein |
| Q9BW30;P59282 | Q9BW30 | Tubulin polymerization-promoting protein family member 3 |
| Q9BW83 | Q9BW83 | Intraflagellar transport protein 27 homolog |
| Q9BW91 | Q9BW91 | ADP-ribose pyrophosphatase, mitochondrial |
| Q9BWD1 | Q9BWD1 | Acetyl-CoA acetyltransferase, cytosolic |
| Q9BX68 | Q9BX68 | Adenosine 5-monophosphoramidase HINT2 |
| Q9BXC9 | Q9BXC9 | Bardet-Biedl syndrome 2 protein |
| Q9BXF9 | Q9BXF9 | Tektin-3 |
| Q9BXI6 | Q9BXI6 | TBC1 domain family member 10A |
| Q9BXM9 | Q9BXM9 | FSD1-like protein |
| Q9BXY5 | Q9BXY5 | Calcyphosin-2 |
| Q9BY32 | Q9BY32 | Inosine triphosphate pyrophosphatase |
| Q9BY43 | Q9BY43 | Charged multivesicular body protein 4a |
| Q9BYT3 | Q9BYT3 | Serine/threonine-protein kinase 33 |
| Q9BYV8 | Q9BYV8 | Centrosomal protein of 41 kDa |

| | | |
|---|---|---|
| Q9BZL1 | Q9BZL1 | Ubiquitin-like protein 5 |
| Q9BZQ8;REV__Q9UHV! | Q9BZQ8 | Protein Niban 1 |
| Q9BZV1 | Q9BZV1 | UBX domain-containing protein 6 |
| Q9BZW7 | Q9BZW7 | Testis-specific gene 10 protein |
| Q9C005 | Q9C005 | Protein dpy-30 homolog |
| Q9C093 | Q9C093 | Sperm flagellar protein 2 |
| Q9C0B2 | Q9C0B2 | Cilia- and flagella-associated protein 74 |
| Q9C0C2 | Q9C0C2 | 182 kDa tankyrase-1-binding protein |
| Q9C0G6 | Q9C0G6 | Dynein axonemal heavy chain 6 |
| Q9GZN4 | Q9GZN4 | Brain-specific serine protease 4 |
| Q9GZP8 | Q9GZP8 | Immortalization up-regulated protein |
| Q9GZS0 | Q9GZS0 | Dynein axonemal intermediate chain 2 |
| Q9GZS3 | Q9GZS3 | Superkiller complex protein 8;Superkiller complex protein { |
| Q9GZT4 | Q9GZT4 | Serine racemase |
| Q9GZT8 | Q9GZT8 | NIF3-like protein 1 |
| Q9GZX3;Q9GZS9;Q8N( | Q9GZX3;Q9GZS9 | Carbohydrate sulfotransferase 6;Carbohydrate sulfotransf |
| Q9H008 | Q9H008 | Phospholysine phosphohistidine inorganic pyrophosphate |
| Q9H069 | Q9H069 | Dynein regulatory complex subunit 3 |
| Q9H095 | Q9H095 | Dynein regulatory complex protein 9 |
| Q9H098 | Q9H098 | Protein FAM107B |
| Q9H0C1 | Q9H0C1 | Zinc finger MYND domain-containing protein 12 |
| Q9H0E2 | Q9H0E2 | Toll-interacting protein |
| Q9H0F7 | Q9H0F7 | ADP-ribosylation factor-like protein 6 |
| Q9H0I2 | Q9H0I2 | Enkurin domain-containing protein 1 |
| Q9H0I3 | Q9H0I3 | Coiled-coil domain-containing protein 113 |
| Q9H0P0 | Q9H0P0 | Cytosolic 5-nucleotidase 3A |
| Q9H0R4 | Q9H0R4 | Haloacid dehalogenase-like hydrolase domain-containing |
| Q9H0U4;Q92928 | Q9H0U4 | Ras-related protein Rab-1B |
| Q9H0W9 | Q9H0W9 | Ester hydrolase C11orf54 |
| Q9H190 | Q9H190 | Syntenin-2 |
| Q9H1C7 | Q9H1C7 | Cysteine-rich and transmembrane domain-containing prot |
| Q9H1P6 | Q9H1P6 | Ciliary microtubule inner protein 1 |
| Q9H1X1 | Q9H1X1 | Radial spoke head protein 9 homolog |
| Q9H223 | Q9H223 | EH domain-containing protein 4 |
| Q9H299 | Q9H299 | SH3 domain-binding glutamic acid-rich-like protein 3 |
| Q9H2G2;O94804 | Q9H2G2 | STE20-like serine/threonine-protein kinase |
| Q9H2H8 | Q9H2H8 | Peptidyl-prolyl cis-trans isomerase-like 3 |
| Q9H2U2 | Q9H2U2 | Inorganic pyrophosphatase 2, mitochondrial |
| Q9H3K6 | Q9H3K6 | BolA-like protein 2 |
| Q9H3R2 | Q9H3R2 | Mucin-13 |
| Q9H3Z4 | Q9H3Z4 | DnaJ homolog subfamily C member 5 |
| Q9H444;Q96CF2;P590 | Q9H444 | Charged multivesicular body protein 4b |
| Q9H477 | Q9H477 | Ribokinase |
| Q9H4A4 | Q9H4A4 | Aminopeptidase B |
| Q9H4G0 | Q9H4G0 | Band 4.1-like protein 1 |
| Q9H4G4 | Q9H4G4 | Golgi-associated plant pathogenesis-related protein 1 |
| Q9H4K1;Q8IYS0 | Q9H4K1 | RIB43A-like with coiled-coils protein 2 |

| | | |
|---|---|---|
| Q9H4M9 | Q9H4M9 | EH domain-containing protein 1 |
| Q9H5F2 | Q9H5F2 | Cilia- and flagella-associated protein 68 |
| Q9H6L2 | Q9H6L2 | Transmembrane protein 231 |
| Q9H6S3 | Q9H6S3 | Epidermal growth factor receptor kinase substrate 8-like pr |
| Q9H7C9 | Q9H7C9 | Mth938 domain-containing protein |
| Q9H7X7 | Q9H7X7 | Intraflagellar transport protein 22 homolog |
| Q9H832 | Q9H832 | Ubiquitin-conjugating enzyme E2 Z |
| Q9H8H3 | Q9H8H3 | Thiol S-methyltransferase TMT1A |
| Q9H8W4 | Q9H8W4 | Pleckstrin homology domain-containing family F member 2 |
| Q9H8Y8 | Q9H8Y8 | Golgi reassembly-stacking protein 2 |
| Q9H910 | Q9H910 | Jupiter microtubule associated homolog 2 |
| Q9H977 | Q9H977 | WD repeat-containing protein 54 |
| Q9H993 | Q9H993 | Damage-control phosphatase ARMT1 |
| Q9H9B4;Q9BWM7 | Q9H9B4 | Sideroflexin-1 |
| Q9HA64 | Q9HA64 | Ketosamine-3-kinase |
| Q9HAB8 | Q9HAB8 | Phosphopantothenate--cysteine ligase |
| Q9HAE3 | Q9HAE3 | Calaxin |
| Q9HAQ2 | Q9HAQ2 | Kinesin-like protein KIF9 |
| Q9HAT2 | Q9HAT2 | Sialate O-acetylesterase |
| Q9HB07 | Q9HB07 | MYG1 exonuclease |
| Q9HB40 | Q9HB40 | Retinoid-inducible serine carboxypeptidase |
| Q9HB71 | Q9HB71 | Calcyclin-binding protein |
| Q9HBG4 | Q9HBG4 | V-type proton ATPase 116 kDa subunit a 4 |
| Q9HBG6 | Q9HBG6 | Intraflagellar transport protein 122 homolog |
| Q9HBL8 | Q9HBL8 | NmrA-like family domain-containing protein 1 |
| Q9HC38 | Q9HC38 | Glyoxalase domain-containing protein 4 |
| Q9HC84 | Q9HC84 | Mucin-5B |
| Q9HCC0 | Q9HCC0 | Methylcrotonoyl-CoA carboxylase beta chain, mitochondri |
| Q9HCY8 | Q9HCY8 | Protein S100-A14 |
| Q9HD42 | Q9HD42 | Charged multivesicular body protein 1a |
| Q9HD45 | Q9HD45 | Transmembrane 9 superfamily member 3 |
| Q9NP55 | Q9NP55 | BPI fold-containing family A member 1 |
| Q9NP61;Q8N6H7 | Q9NP61 | ADP-ribosylation factor GTPase-activating protein 3 |
| Q9NP79 | Q9NP79 | Vacuolar protein sorting-associated protein VTA1 homolog |
| Q9NP97 | Q9NP97 | Dynein light chain roadblock-type 1 |
| Q9NPA8 | Q9NPA8 | Transcription and mRNA export factor ENY2 |
| Q9NPC4 | Q9NPC4 | Lactosylceramide 4-alpha-galactosyltransferase |
| Q9NPH2 | Q9NPH2 | Inositol-3-phosphate synthase 1 |
| Q9NQ32 | Q9NQ32 | Uncharacterized protein C11orf16 |
| Q9NQ48 | Q9NQ48 | Leucine zipper transcription factor-like protein 1 |
| Q9NQ84 | Q9NQ84 | G-protein coupled receptor family C group 5 member C |
| Q9NQ88 | Q9NQ88 | Fructose-2,6-bisphosphatase TIGAR |
| Q9NQC3 | Q9NQC3 | Reticulon-4 |
| Q9NQC8 | Q9NQC8 | Intraflagellar transport protein 46 homolog |
| Q9NQE9 | Q9NQE9 | Adenosine 5-monophosphoramidase HINT3 |
| Q9NQR4 | Q9NQR4 | Omega-amidase NIT2 |
| Q9NQW7;O00462 | Q9NQW7 | Xaa-Pro aminopeptidase 1 |

| | | |
|---|---|---|
| Q9NR19 | Q9NR19 | Acetyl-coenzyme A synthetase, cytoplasmic |
| Q9NR31 | Q9NR31 | GTP-binding protein SAR1a |
| Q9NR45 | Q9NR45 | Sialic acid synthase |
| Q9NR46 | Q9NR46 | Endophilin-B2 |
| Q9NR56 | Q9NR56 | Muscleblind-like protein 1 |
| Q9NRD9;Q9NRD8 | Q9NRD9 | Dual oxidase 1 |
| Q9NRL2 | Q9NRL2 | Bromodomain adjacent to zinc finger domain protein 1A |
| Q9NRN7 | Q9NRN7 | L-aminoadipate-semialdehyde dehydrogenase-phosphopa |
| Q9NRP0 | Q9NRP0 | Oligosaccharyltransferase complex subunit OSTC |
| Q9NRV9 | Q9NRV9 | Heme-binding protein 1 |
| Q9NRX4 | Q9NRX4 | 14 kDa phosphohistidine phosphatase |
| Q9NS86 | Q9NS86 | LanC-like protein 2 |
| Q9NSC7 | Q9NSC7 | Alpha-N-acetylgalactosaminide alpha-2,6-sialyltransferase |
| Q9NSE4 | Q9NSE4 | Isoleucine--tRNA ligase, mitochondrial |
| Q9NSK0 | Q9NSK0 | Kinesin light chain 4 |
| Q9NT62 | Q9NT62 | Ubiquitin-like-conjugating enzyme ATG3 |
| Q9NTK5 | Q9NTK5 | Obg-like ATPase 1 |
| Q9NTX5 | Q9NTX5 | Ethylmalonyl-CoA decarboxylase |
| Q9NU02 | Q9NU02 | Ankyrin repeat and EF-hand domain-containing protein 1 |
| Q9NUQ6 | Q9NUQ6 | SPATS2-like protein |
| Q9NUQ9;Q9H0Q0 | Q9NUQ9 | CYFIP-related Rac1 interactor B |
| Q9NVA2;Q14141 | Q9NVA2 | Septin-11 |
| Q9NVI7;Q5T9A4;Q5T2N | Q9NVI7;Q5T9A4 | ATPase family AAA domain-containing protein 3A;ATPase fa |
| Q9NVJ2 | Q9NVJ2 | ADP-ribosylation factor-like protein 8B |
| Q9NVP4 | Q9NVP4 | Double zinc ribbon and ankyrin repeat-containing protein 1 |
| Q9NVS9 | Q9NVS9 | Pyridoxine-5-phosphate oxidase |
| Q9NWB7 | Q9NWB7 | Intraflagellar transport protein 57 homolog |
| Q9NWV4 | Q9NWV4 | CXXC motif containing zinc binding protein |
| Q9NXA8 | Q9NXA8 | NAD-dependent protein deacylase sirtuin-5, mitochondrial |
| Q9NY33 | Q9NY33 | Dipeptidyl peptidase 3 |
| Q9NYC9 | Q9NYC9 | Dynein axonemal heavy chain 9 |
| Q9NZ01 | Q9NZ01 | Very-long-chain enoyl-CoA reductase |
| Q9NZ45 | Q9NZ45 | CDGSH iron-sulfur domain-containing protein 1 |
| Q9NZB2 | Q9NZB2 | Constitutive coactivator of PPAR-gamma-like protein 1 |
| Q9NZD2 | Q9NZD2 | Glycolipid transfer protein |
| Q9NZF1 | Q9NZF1 | Placenta-specific gene 8 protein |
| Q9NZL4 | Q9NZL4 | Hsp70-binding protein 1 |
| Q9NZL9 | Q9NZL9 | Methionine adenosyltransferase 2 subunit beta |
| Q9NZM1;O75923 | Q9NZM1 | Myoferlin |
| Q9NZZ3 | Q9NZZ3 | Charged multivesicular body protein 5 |
| Q9P035 | Q9P035 | Very-long-chain (3R)-3-hydroxyacyl-CoA dehydratase 3 |
| Q9P0L0 | Q9P0L0 | Vesicle-associated membrane protein-associated protein A |
| Q9P0R6 | Q9P0R6 | GSK3B-interacting protein |
| Q9P0V9 | Q9P0V9 | Septin-10 |
| Q9P1F3 | Q9P1F3 | Costars family protein ABRACL |
| Q9P1V8 | Q9P1V8 | Sterile alpha motif domain-containing protein 15 |
| Q9P1Y5 | Q9P1Y5 | Calmodulin-regulated spectrin-associated protein 3 |

| | | |
|---|---|---|
| Q9P1Z9 | Q9P1Z9 | Coiled-coil domain-containing protein 180 |
| Q9P209 | Q9P209 | Centrosomal protein of 72 kDa |
| Q9P225 | Q9P225 | Dynein axonemal heavy chain 2 |
| Q9P287 | Q9P287 | BRCA2 and CDKN1A-interacting protein |
| Q9P2B7 | Q9P2B7 | Cilia- and flagella-associated protein 97 |
| Q9P2D7 | Q9P2D7 | Dynein axonemal heavy chain 1 |
| Q9P2H3 | Q9P2H3 | Intraflagellar transport protein 80 homolog |
| Q9P2J5 | Q9P2J5 | Leucine--tRNA ligase, cytoplasmic |
| Q9P2L0 | Q9P2L0 | WD repeat-containing protein 35 |
| Q9P2M1 | Q9P2M1 | LRP2-binding protein |
| Q9P2R7 | Q9P2R7 | Succinate--CoA ligase [ADP-forming] subunit beta, mitocho |
| Q9P2S6 | Q9P2S6 | Ankyrin repeat and MYND domain-containing protein 1 |
| Q9P2T1 | Q9P2T1 | GMP reductase 2 |
| Q9UBC9 | Q9UBC9 | Small proline-rich protein 3 |
| Q9UBG3 | Q9UBG3 | Cornulin |
| Q9UBI1 | Q9UBI1 | COMM domain-containing protein 3 |
| Q9UBI6 | Q9UBI6 | Guanine nucleotide-binding protein G(I)/G(S)/G(O) subunit |
| Q9UBQ0 | Q9UBQ0 | Vacuolar protein sorting-associated protein 29 |
| Q9UBQ7 | Q9UBQ7 | Glyoxylate reductase/hydroxypyruvate reductase |
| Q9UBR2 | Q9UBR2 | Cathepsin Z |
| Q9UBX7 | Q9UBX7 | Kallikrein-11;Kallikrein-11 inactive chain 1;Kallikrein-11 ina |
| Q9UDW1 | Q9UDW1 | Cytochrome b-c1 complex subunit 9 |
| Q9UDX4 | Q9UDX4 | SEC14-like protein 3 |
| Q9UDY2 | Q9UDY2 | Tight junction protein ZO-2 |
| Q9UDY4 | Q9UDY4 | DnaJ homolog subfamily B member 4 |
| Q9UEW8 | Q9UEW8 | STE20/SPS1-related proline-alanine-rich protein kinase |
| Q9UFE4 | Q9UFE4 | Coiled-coil domain-containing protein 39 |
| Q9UG01 | Q9UG01 | Intraflagellar transport protein 172 homolog |
| Q9UGI8 | Q9UGI8 | Testin |
| Q9UH65 | Q9UH65 | Switch-associated protein 70 |
| Q9UHA4 | Q9UHA4 | Ragulator complex protein LAMTOR3 |
| Q9UHB6 | Q9UHB6 | LIM domain and actin-binding protein 1 |
| Q9UHD1 | Q9UHD1 | Cysteine and histidine-rich domain-containing protein 1 |
| Q9UHD8 | Q9UHD8 | Septin-9 |
| Q9UHG0 | Q9UHG0 | Doublecortin domain-containing protein 2 |
| Q9UHG3 | Q9UHG3 | Prenylcysteine oxidase 1 |
| Q9UHL4 | Q9UHL4 | Dipeptidyl peptidase 2 |
| Q9UHP6 | Q9UHP6 | Radial spoke head 14 homolog |
| Q9UHQ9 | Q9UHQ9 | NADH-cytochrome b5 reductase 1 |
| Q9UHR4 | Q9UHR4 | Brain-specific angiogenesis inhibitor 1-associated protein 2 |
| Q9UHV9 | Q9UHV9 | Prefoldin subunit 2 |
| Q9UHY7 | Q9UHY7 | Enolase-phosphatase E1 |
| Q9UI12 | Q9UI12 | V-type proton ATPase subunit H |
| Q9UI30 | Q9UI30 | Multifunctional methyltransferase subunit TRM112-like pro |
| Q9UI46 | Q9UI46 | Dynein axonemal intermediate chain 1 |
| Q9UIC8 | Q9UIC8 | Leucine carboxyl methyltransferase 1 |
| Q9UIF3 | Q9UIF3 | Tektin-2 |

| | | |
|---|---|---|
| Q9UII2 | Q9UII2 | ATPase inhibitor, mitochondrial |
| Q9UJ70 | Q9UJ70 | N-acetyl-D-glucosamine kinase |
| Q9UJC5 | Q9UJC5 | SH3 domain-binding glutamic acid-rich-like protein 2 |
| Q9UJS0;O75746 | Q9UJS0;O75746 | Electrogenic aspartate/glutamate antiporter SLC25A13, mi |
| Q9UJU6 | Q9UJU6 | Drebrin-like protein |
| Q9UJZ1 | Q9UJZ1 | Stomatin-like protein 2, mitochondrial |
| Q9UK22 | Q9UK22 | F-box only protein 2 |
| Q9UK41 | Q9UK41 | Vacuolar protein sorting-associated protein 28 homolog |
| Q9UK76 | Q9UK76 | Jupiter microtubule associated homolog 1;Jupiter microtub |
| Q9UKK9 | Q9UKK9 | ADP-sugar pyrophosphatase |
| Q9UKM9 | Q9UKM9 | RNA-binding protein Raly |
| Q9UKY7 | Q9UKY7 | Protein CDV3 homolog |
| Q9UL16 | Q9UL16 | Cilia- and flagella-associated protein 45 |
| Q9UL46 | Q9UL46 | Proteasome activator complex subunit 2 |
| Q9ULA0 | Q9ULA0 | Aspartyl aminopeptidase |
| Q9ULC3 | Q9ULC3 | Ras-related protein Rab-23 |
| Q9ULC4;A0A3B3IRV3 | Q9ULC4;A0A3B3IRV3 | Malignant T-cell-amplified sequence 1;Malignant T-cell-am |
| Q9ULD2 | Q9ULD2 | Microtubule-associated tumor suppressor 1 |
| Q9ULV4 | Q9ULV4 | Coronin-1C |
| Q9ULZ3 | Q9ULZ3 | Apoptosis-associated speck-like protein containing a CARD |
| Q9UM00 | Q9UM00 | Calcium load-activated calcium channel |
| Q9UM54 | Q9UM54 | Unconventional myosin-VI |
| Q9UMR2 | Q9UMR2 | ATP-dependent RNA helicase DDX19B |
| Q9UMY4 | Q9UMY4 | Sorting nexin-12 |
| Q9UN36 | Q9UN36 | Protein NDRG2 |
| Q9UN76 | Q9UN76 | Sodium- and chloride-dependent neutral and basic amino a |
| Q9UNF0 | Q9UNF0 | Protein kinase C and casein kinase substrate in neurons pr |
| Q9UNH7 | Q9UNH7 | Sorting nexin-6;Sorting nexin-6, N-terminally processed |
| Q9UNL2 | Q9UNL2 | Translocon-associated protein subunit gamma |
| Q9UNM6 | Q9UNM6 | 26S proteasome non-ATPase regulatory subunit 13 |
| Q9UNN5 | Q9UNN5 | FAS-associated factor 1 |
| Q9UNS2 | Q9UNS2 | COP9 signalosome complex subunit 3 |
| Q9UNT1;Q9UBK7 | Q9UNT1;Q9UBK7 | Rab-like protein 2B;Rab-like protein 2A |
| Q9UNZ2 | Q9UNZ2 | NSFL1 cofactor p47 |
| Q9UPY8;Q15555 | Q9UPY8 | Microtubule-associated protein RP/EB family member 3 |
| Q9UPZ9 | Q9UPZ9 | Serine/threonine-protein kinase ICK |
| Q9UQ07 | Q9UQ07 | MAPK/MAK/MRK overlapping kinase |
| Q9UQ80 | Q9UQ80 | Proliferation-associated protein 2G4 |
| Q9UQB8 | Q9UQB8 | Brain-specific angiogenesis inhibitor 1-associated protein 2 |
| Q9UQN3 | Q9UQN3 | Charged multivesicular body protein 2b |
| Q9Y223 | Q9Y223 | Bifunctional UDP-N-acetylglucosamine 2-epimerase/N-ace |
| Q9Y224 | Q9Y224 | RNA transcription, translation and transport factor protein |
| Q9Y230 | Q9Y230 | RuvB-like 2 |
| Q9Y237 | Q9Y237 | Peptidyl-prolyl cis-trans isomerase NIMA-interacting 4 |
| Q9Y238 | Q9Y238 | Deleted in lung and esophageal cancer protein 1 |
| Q9Y259 | Q9Y259 | Choline/ethanolamine kinase |
| Q9Y265 | Q9Y265 | RuvB-like 1 |

| | | |
|---|---|---|
| Q9Y266 | Q9Y266 | Nuclear migration protein nudC |
| Q9Y277 | Q9Y277 | Voltage-dependent anion-selective channel protein 3 |
| Q9Y2A7 | Q9Y2A7 | Nck-associated protein 1 |
| Q9Y2B0 | Q9Y2B0 | Protein canopy homolog 2 |
| Q9Y2E6 | Q9Y2E6 | E3 ubiquitin-protein ligase DTX4 |
| Q9Y2Q3 | Q9Y2Q3 | Glutathione S-transferase kappa 1 |
| Q9Y2S2 | Q9Y2S2 | Lambda-crystallin homolog |
| Q9Y2S6;A0A024R1R8 | Q9Y2S6;A0A024R1R8 | Translation machinery-associated protein 7;Translation ma |
| Q9Y2T3 | Q9Y2T3 | Guanine deaminase |
| Q9Y2V2 | Q9Y2V2 | Calcium-regulated heat-stable protein 1 |
| Q9Y2Z0 | Q9Y2Z0 | Protein SGT1 homolog |
| Q9Y305 | Q9Y305 | Acyl-coenzyme A thioesterase 9, mitochondrial |
| Q9Y365 | Q9Y365 | START domain-containing protein 10 |
| Q9Y366 | Q9Y366 | Intraflagellar transport protein 52 homolog |
| Q9Y371 | Q9Y371 | Endophilin-B1 |
| Q9Y376 | Q9Y376 | Calcium-binding protein 39 |
| Q9Y3A5 | Q9Y3A5 | Ribosome maturation protein SBDS |
| Q9Y3B3 | Q9Y3B3 | Transmembrane emp24 domain-containing protein 7 |
| Q9Y3B8 | Q9Y3B8 | Oligoribonuclease, mitochondrial |
| Q9Y3C6 | Q9Y3C6 | Peptidyl-prolyl cis-trans isomerase-like 1 |
| Q9Y3C8 | Q9Y3C8 | Ubiquitin-fold modifier-conjugating enzyme 1 |
| Q9Y3D6 | Q9Y3D6 | Mitochondrial fission 1 protein |
| Q9Y3E7 | Q9Y3E7 | Charged multivesicular body protein 3 |
| Q9Y3F4 | Q9Y3F4 | Serine-threonine kinase receptor-associated protein |
| Q9Y3I0 | Q9Y3I0 | RNA-splicing ligase RtcB homolog |
| Q9Y3Q3 | Q9Y3Q3 | Transmembrane emp24 domain-containing protein 3 |
| Q9Y3Z3 | Q9Y3Z3 | Deoxynucleoside triphosphate triphosphohydrolase SAMH |
| Q9Y446 | Q9Y446 | Plakophilin-3 |
| Q9Y490;Q9Y4G6 | Q9Y490 | Talin-1 |
| Q9Y496;Q6ZMV9 | Q9Y496 | Kinesin-like protein KIF3A |
| Q9Y4L1 | Q9Y4L1 | Hypoxia up-regulated protein 1 |
| Q9Y4P9 | Q9Y4P9 | Sperm flagellar protein 1 |
| Q9Y4Y9 | Q9Y4Y9 | U6 snRNA-associated Sm-like protein LSm5 |
| Q9Y512 | Q9Y512 | Sorting and assembly machinery component 50 homolog |
| Q9Y547 | Q9Y547 | Intraflagellar transport protein 25 homolog |
| Q9Y570 | Q9Y570 | Protein phosphatase methylesterase 1 |
| Q9Y5B8 | Q9Y5B8 | Nucleoside diphosphate kinase homolog 7 |
| Q9Y5K8 | Q9Y5K8 | V-type proton ATPase subunit D |
| Q9Y5P6 | Q9Y5P6 | Mannose-1-phosphate guanyltransferase beta |
| Q9Y5R8 | Q9Y5R8 | Trafficking protein particle complex subunit 1 |
| Q9Y5Y2 | Q9Y5Y2 | Cytosolic Fe-S cluster assembly factor NUBP2 |
| Q9Y5Z4 | Q9Y5Z4 | Heme-binding protein 2 |
| Q9Y617 | Q9Y617 | Phosphoserine aminotransferase |
| Q9Y678;Q9UBF2 | Q9Y678 | Coatomer subunit gamma-1 |
| Q9Y696 | Q9Y696 | Chloride intracellular channel protein 4 |
| Q9Y6A4 | Q9Y6A4 | Cilia- and flagella-associated protein 20 |
| Q9Y6B6 | Q9Y6B6 | GTP-binding protein SAR1b |

| | | |
|---|---|---|
| Q9Y6C9 | Q9Y6C9 | Mitochondrial carrier homolog 2 |
| Q9Y6E0;O00506 | Q9Y6E0 | Serine/threonine-protein kinase 24;Serine/threonine-prote |
| Q9Y6G9 | Q9Y6G9 | Cytoplasmic dynein 1 light intermediate chain 1 |
| Q9Y6J8 | Q9Y6J8 | Serine/threonine/tyrosine-interacting-like protein 1 |
| Q9Y6M9 | Q9Y6M9 | NADH dehydrogenase [ubiquinone] 1 beta subcomplex sub |
| Q9Y6N5 | Q9Y6N5 | Sulfide:quinone oxidoreductase, mitochondrial |
| Q9Y6R7 | Q9Y6R7 | IgGFc-binding protein |
| Q9Y6U3 | Q9Y6U3 | Scinderin |

T4B;Diphosphoinositol polyphosphate phosphohydrolase 2;Diphosphoinositol polyphosphate phosphoh

ase 1;Sulfate adenylyltransferase;Adenylyl-sulfate kinase;Bifunctional 3-phosphoadenosine 5-phospho ype 1-N;Histone H2B type 1-H;Histone H2B type 2-F;Histone H2B type 1-C/E/F/G/I;Histone H2B type 1-D

sminogen activator long chain A;Urokinase-type plasminogen activator short chain A;Urokinase-type plas ɔmplement C3 alpha chain;C3a anaphylatoxin;Acylation stimulating protein;Complement C3b alpha cha geneous nuclear ribonucleoprotein C-like 4;Heterogeneous nuclear ribonucleoprotein C-like 1;Heteroge ɔus nuclear ribonucleoprotein A1, N-terminally processed;Heterogeneous nuclear ribonucleoprotein A1·

ɔid protein A(2-104);Serum amyloid protein A(3-104);Serum amyloid protein A(2-103);Serum amyloid prc ahydrofolate dehydrogenase;Methenyltetrahydrofolate cyclohydrolase;Formyltetrahydrofolate syntheta

sphoribosylamine--glycine ligase;Phosphoribosylformylglycinamidine cyclo-ligase;Phosphoribosylglycin
hosphoribosylaminoimidazole succinocarboxamide synthetase;Phosphoribosylaminoimidazole carbox

sylaminoimidazolecarboxamide formyltransferase;Inosine 5-monophosphate cyclohydrolase;Bifunctior

:ytochrome b-c1 complex subunit 9;Putative cytochrome b-c1 complex subunit Rieske-like protein 1

se;[Acyl-carrier-protein] S-malonyltransferase;3-oxoacyl-[acyl-carrier-protein] synthase;3-oxoacyl-[acyl

nain death agonist p15;BH3-interacting domain death agonist p13;BH3-interacting domain death agonis

Small ribosomal subunit protein eS31;Ubiquitin-ribosomal protein eL40 fusion protein;Ubiquitin;Large ri rocessed variant 1;Small-inducible cytokine B6, N-processed variant 2;Small-inducible cytokine B6, N-p

ctivated chloride channel regulator 4, 110 kDa form;Calcium-activated chloride channel regulator 4, 30 nst decapentaplegic homolog 3;Mothers against decapentaplegic homolog 5;Mothers against decapenta

ain;Tubulin alpha-3D chain;Detyrosinated tubulin alpha-3D chain;Tubulin alpha-3C chain;Detyrosinated

etylmannosamine kinase;UDP-N-acetylglucosamine 2-epimerase (hydrolyzing);N-acetylmannosamine k

ain;Complement C3c alpha chain fragment 1;Complement C3dg fragment;Complement C3g fragment;(

[illegible]eneous nuclear ribonucleoprotein C-like 3;Heterogeneous nuclear ribonucleoprotein C-like 2 ɔtein A(2-102);Serum amyloid protein A(4-101);Serum amyloid A-2 protein;Amyloid A2 protein

[illegible]-carrier-protein] reductase;3-hydroxyacyl-[acyl-carrier-protein] dehydratase;Enoyl-[acyl-carrier-protein] [illegible]

Complement C3d fragment;Complement C3f fragment;Complement C3c alpha chain fragment 2
