## Supplementary material for "Novel protein networks in motile cilia from human epithelial cells revealed by cryo-ET and proteomics analysis of cilia from PCD patients": SupplTable2

| Protein IDs | Protein names | Gene names | DNAH5 PD122 | DNAH5 PD441 | DNAH5 PD480 | DNAH11 PD442 | DNAH11 PD551 | DNAH9 PD222 | DNAH9 PD272 | IDA PD691 | CP PD311 | CP PD343 | CP PD358 | unknown PD403 | unknown PD428 | unkown PD461 |
|---|---|---|---|---|---|---|---|---|---|---|---|---|---|---|---|---|
| Q9P2D7 | Dynein axonemal heavy chain 1 | DNAH1 | - | - | - | - | - | | | - | | - | | - | - | - |
| Q8IVF4;REV__Q9Y5K6 | Dynein axonemal heavy chain 10 | DNAH10 | - | - | - | - | - | | | | | | | | - | |
| Q96DT5;Q0VDD8 | Dynein axonemal heavy chain 11 | DNAH11 | - | | | - | - | | | | | | | | | |
| Q6ZR08 | Dynein axonemal heavy chain 12 | DNAH12 | - | - | - | - | - | - | | - | | - | | - | - | - |
| Q9P225 | Dynein axonemal heavy chain 2 | DNAH2 | - | - | - | - | - | | | - | | - | | - | | - |
| Q8TD57 | Dynein axonemal heavy chain 3 | DNAH3 | | - | | | | | | | | | | | | |
| Q8TE73;Q9NRZ9 | Dynein axonemal heavy chain 5 | DNAH5 | - | | | - | - | | - | | | | | | | |
| Q9C0G6 | Dynein axonemal heavy chain 6 | DNAH6 | - | - | - | - | - | | | - | | - | | - | | - |
| Q8WXX0 | Dynein axonemal heavy chain 7 | DNAH7 | - | - | - | - | - | - | - | - | | - | | - | | - |
| Q9NYC9 | Dynein axonemal heavy chain 9 | DNAH9 | - | - | - | - | - | | | - | | | | - | | |
| Q9UI46 | Dynein axonemal intermediate chain 1 | DNAI1 | - | - | - | - | - | | | - | | | | - | | - |
| Q9GZS0 | Dynein axonemal intermediate chain 2 | DNAI2 | - | - | - | - | - | | | - | | | | - | | - |
| Q8IWG1 | Dynein axonemal intermediate chain 3 | DNAI3 | - | - | - | - | - | | | - | | | | - | | - |
| Q5VTH9 | Dynein axonemal intermediate chain 4 | DNAI4 | - | - | - | - | - | | | - | - | - | | - | | - |

- Supplementary Table 2. MS data of 14 patients to show proteins which decreased significantly in the patients' cilia compared to WT (marked by -).
