## Supplementary material for "Novel protein networks in motile cilia from human epithelial cells revealed by cryo-ET and proteomics analysis of cilia from PCD patients": SupplTable3

| Protein IDs | Protein names | Gene names | DNAH5 PD122 | DNAH5 PD441 | DNAH5 PD480 | DNAH11 PD442 | DNAH11 PD551 | DNAH9 PD222 | DNAH9 PD272 | IDA PD691 | CP PD311 | CP PD343 | CP PD358 | unknown PD403 | unknown PD428 | unkown PD461 |
|---|---|---|---|---|---|---|---|---|---|---|---|---|---|---|---|---|
| Q8WYR4 | Radial spoke head 1 homolog | RSPH1 | - | - | - | - | - | - | - | | | | | - | | - |
| P0C881 | Radial spoke head 10 homolog B | RSPH10B | | | | | | | | | | | | | | |
| B2RC85 | Radial spoke head 10 homolog B2 | RSPH10B2 | | - | | - | | | | - | | - | | - | | - |
| Q9UHP6 | Radial spoke head 14 homolog | RSPH14 | - | - | - | - | - | - | - | | | | | - | | - |
| Q86UC2 | Radial spoke head protein 3 homolog | RSPH3 | - | - | - | - | - | - | - | | | | | - | | - |
| Q5TD94 | Radial spoke head protein 4 homolog A | RSPH4A | - | - | | - | - | | | | | | | - | | - |
