## Supplementary material for "Novel protein networks in motile cilia from human epithelial cells revealed by cryo-ET and proteomics analysis of cilia from PCD patients": SupplTable4

| Protein IDs | Protein names | Gene names | DNAH5 PD122 | DNAH5 PD441 | DNAH5 PD480 | DNAH11 PD442 | DNAH11 PD551 | DNAH9 PD222 | DNAH9 PD272 | IDA PD691 | CP PD311 | CP PD343 | CP PD358 | unknown PD403 | unknown PD428 | unkown PD461 |
|---|---|---|---|---|---|---|---|---|---|---|---|---|---|---|---|---|
| Q9HBG6 | Intraflagellar transport protein 122 homolog | IFT122 | | - | | - | | | | | | | | | | - |
| Q96RY7 | Intraflagellar transport protein 140 homolog | IFT140 | | - | | - | | - | - | | | | | - | | - |
| Q9UG01 | Intraflagellar transport protein 172 homolog | IFT172 | | - | | - | - | - | - | | | | | - | | - |
| Q8IY31 | Intraflagellar transport protein 20 homolog | IFT20 | | - | | - | - | | | | | | | - | | - |
| Q9H7X7 | Intraflagellar transport protein 22 homolog | IFT22 | | - | | - | - | - | - | | | | | - | | - |
| Q9Y547 | Intraflagellar transport protein 25 homolog | IFT25 | | - | | - | - | | | | - | | | - | | - |
| Q9BW83 | Intraflagellar transport protein 27 homolog | IFT27 | | - | | - | - | - | - | | | | | - | | - |
| Q96FT9 | Intraflagellar transport protein 43 homolog | IFT43 | | - | | - | - | | | | | | | - | | - |
| Q9NQC8 | Intraflagellar transport protein 46 homolog | IFT46 | | - | | - | - | - | - | | - | | | - | | - |
| Q9Y366 | Intraflagellar transport protein 52 homolog | IFT52 | | - | | - | - | - | - | | | | | - | | - |
| A0AVF1 | Intraflagellar transport protein 56 | IFT56 | | - | | - | - | - | | | | | | - | | - |
| Q9NWB7 | Intraflagellar transport protein 57 homolog | IFT57 | | - | | - | - | - | - | | - | | | - | | - |
| Q86WT1 | Intraflagellar transport protein 70A | IFT70A | | - | | - | - | - | | | | | | - | | - |
| Q8N4P2 | Intraflagellar transport protein 70B | IFT70B | | - | | - | - | | | | | | | - | | - |
| Q96LB3 | Intraflagellar transport protein 74 homolog | IFT74 | | - | | - | - | - | - | | - | | | - | | - |
| Q9P2H3 | Intraflagellar transport protein 80 homolog | IFT80 | | - | | - | - | | | | | | | - | | - |
| Q8WYA0 | Intraflagellar transport protein 81 homolog | IFT81 | | - | | - | - | - | - | | | | | - | | - |
